# Developmental origins of thymic B-cell lymphomas

**DOI:** 10.64898/2026.10.17.759359

**Authors:** Shinya Rai, Yifan Yin, Manabu Fujisawa, Makoto Kishida, Michael Yu Li, Jeffrey Niu, Aixiang Jiang, Laura K Hilton, Amos Fong, Gerben Duns, Adèle Telenius, Barbara Meissner, Merrill Boyle, Hisae Nakamura, Alexandra Roos, Shiva Daryani, Luke ÒBrien, Claudia Cassidy, Shaocheng Wu, Eric Lee, Andrew Lytle, Pedro Farinha, Graham Slack, Jonathan W Bush, Ryan D Morin, Leandro Venturutti, Andrew P Weng, Andrew Roth, Brad H Nelson, Kerry J Savage, Tomohiro Aoki, Clementine Sarkozy, David W Scott, Christian Steidl

**Author notes:** Corresponding Authors: Christian Steidl, MD, Centre for Lymphoid Cancer, British Columbia Cancer, Vancouver, BC, Canada 675 West 10th Ave, Room 12-110, Vancouver, BC V5Z 1L3, Canada. Authors contributed equally to this work (Shinya Rai, Yifan Yin).

## Abstract

The thymus provides a specialized niche for lymphocyte differentiation, yet the developmental trajectories of its rare B-cell population and their relationship to thymic B-cell lymphomagenesis remain poorly understood. Here, we integrate single-cell multiomics and spatial profiling across human tissues to delineate a B-cell differentiation program selectively represented in the human thymus. This program links CCL17⁺ and germinal center-like B-cell states along a trajectory distinct from canonical germinal center differentiation and can be recapitulated *in vitro* by signals operating in the thymic medulla. Extending this framework across human B-cell lymphomas, we identify an inherent malignant cell state in PMBCL that shares transcriptional and epigenetic features with CCL17⁺ thymic B cells. This malignant state associates with early clonal positions in reconstructed B-cell receptor phylogenies and persists within tumor microenvironments distinct from the thymic niche. Together, these findings support a model in which thymus-specific B-cell differentiation programs are redeployed during thymic B-cell lymphomagenesis.

**Highlights:**

✓ CCL17⁺ B-cell states are selectively represented in the human thymus
✓ Thymic medullary signals recapitulate CCL17⁺-to-germinal center-like differentiation
✓ Primary mediastinal large B-cell lymphomas harbor a malignant counterpart of CCL17⁺ thymic B cells
✓ The malignant counterpart occupies early nodes in clonal phylogenies

## Introduction

Cancer initiation and progression are driven by a complex interplay between clonally expanding cells and cellular components of tissue-specific tumor microenvironments (TMEs)^1^, in which cellular ecosystems create organ-specific niches for malignant evolution^2–4^. The thymus exemplifies such a niche by providing networks of thymic epithelial cells (TECs) and immune cells in the cortex and medulla that physiologically enable positive and negative T-cell selection for the formation of central immune tolerance^5,6^. Although most primary thymic malignancies are epithelial tumors, including thymoma and thymic carcinoma, B-cell lymphomas can also occur in the thymic niche^7–9^. Yet, how the thymic cellular ecosystem contributes to an environment supporting malignant B-cell transformation and progression, beyond a static cell-of-origin (COO) model, is currently unknown.

Primary mediastinal (thymic) large B-cell lymphoma (PMBCL) is an aggressive large B-cell lymphoma that differs from diffuse large B-cell lymphoma (DLBCL) in clinical presentation, histopathological features, and gene-expression profiles^8,9^. Biologically, many DLBCLs are germinal center (GC)–experienced, and activation-induced cytidine deaminase (AID)-mediated class-switch recombination predisposes to immunoglobulin (IG) translocations involving genes such as *MYC* and *BCL6* in a subset of cases^10,11^. By contrast, such oncogenic rearrangements are uncommon in PMBCL, which is instead characterized by oncogenic Janus kinase–signal transducer and activator of transcription (JAK-STAT) and nuclear factor-κB (NF-κB) signaling, together with immune evasion programs^8,9,12,13^. These hallmarks, along with unique clinico-pathologic features, collectively underpin its classification as a distinct disease entity^14,15^. PMBCL is further characterized by striking tissue-restricted lymphomagenesis, arising in the anterior mediastinum of young adults^8,9^, suggesting a thymic B-cell origin^8,16–19^. However, the relevant normal thymic B-cell states and their relationship to PMBCL phenotypes remain undefined, and, unlike in DLBCL^20,21^, COO-informed therapeutic frameworks have yet to emerge for PMBCL.

Over the past few years, comprehensive analyses of the thymus have broadened our understanding of thymus biology^22–27^. Within this expanding landscape of cellular definitions, thymic B cells have been recognized as a rare and incompletely characterized immune cell population^28^. In mice, thymic B cells function as medullary antigen-presenting cells that promote negative selection and acquire a distinctive phenotype characterized by upregulation of MHC class II, CD80, and AIRE^28,29^. In humans, while single-cell studies have identified activated, chemokine-expressing thymic B-cell populations and evidence of intrathymic plasma-cell differentiation^22,23,30^, their developmental relationships, spatial organization, and links to malignant B-cell states remain unresolved. These knowledge gaps reflect practical challenges, including limited tissue accessibility and the rarity of thymic B cells.

In this study, we assembled a primary human tissue cohort spanning thymus, reactive lymph node (RLN), tonsil, PMBCL, and DLBCL. Using complementary single-cell and single-nucleus transcriptomic, chromatin-accessibility, B-cell receptor (BCR), and spatial profiling, we resolve thymic B-cell developmental programs, their microenvironmental regulation and determine how these programs are redeployed and remodeled in PMBCL.

## Results

### A *CCL17*⁺ B-cell state is selectively enriched in the human thymus

To construct a multi-modal atlas of thymic B-cells, we profiled normal human thymus, RLN, and tonsil samples using complementary single-nucleus and single-cell transcriptomic approaches across formalin-fixed paraffin-embedded (FFPE) and fresh-frozen cohorts. We then extended this framework to B-cell lymphomas, including PMBCL and DLBCL, to assess whether thymus-specific B-cell programs are preserved in malignant contexts. These datasets integrated gene expression, chromatin accessibility, and BCR information to define thymus-specific B-cell states. We further applied high-plex CosMx spatial molecular imaging to both normal thymus and PMBCL tissues to compare the spatial organization and cellular neighborhoods of physiological and tumor-associated B-cell states (Figures 1A and S1, Tables S1-3). Finally, to test whether this thymus-specific phenotype can be induced *in vitro*, we performed cytokine-based stimulation experiments using primary human B cells (Figure 1A).

**Figure 1.**
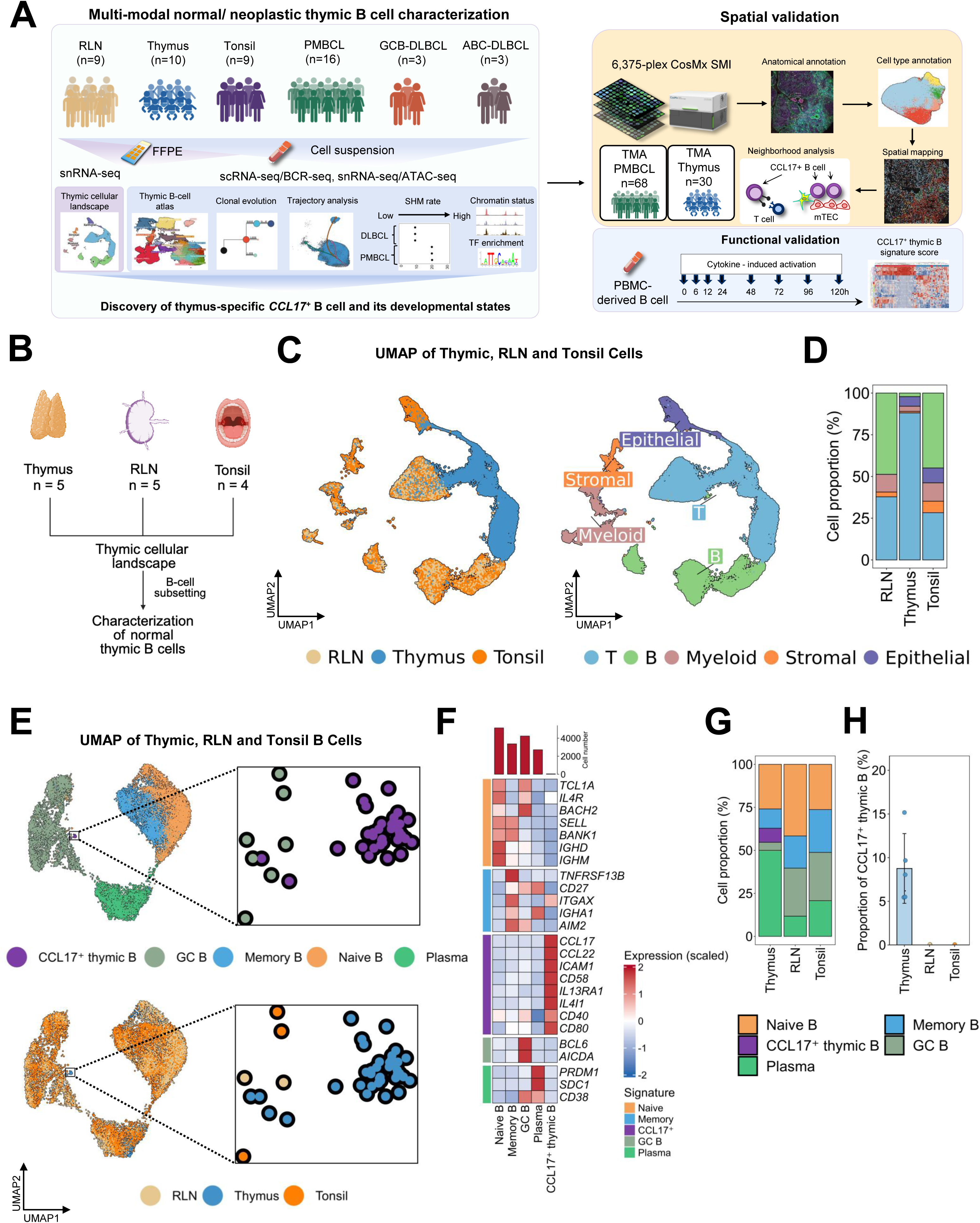
Thymus-specific CCL17⁺ B cells. (A) Overview of tissue sources and multi-modal single-nuclear/cell and spatial profiling strategies. “n” indicates the number of unique cases included. When multiple assays (e.g., single-nucleus RNA-seq, multiome, or BCR sequencing) were performed on the same specimen, the case was counted once. RLN, reactive lymph node; PMBCL, primary mediastinal B-cell lymphoma; GCB-DLBCL, germinal center B-cell–like diffuse large B-cell lymphoma; ABC-DLBCL, activated B-cell–like diffuse large B-cell lymphoma; FFPE, formalin-fixed paraffin-embedded; scRNA-seq, single-cell RNA sequencing; snRNA-seq, single-nucleus RNA sequencing; BCR-seq, B-cell receptor sequencing; ATAC-seq, assay for transposase-accessible chromatin using sequencing; TMA, tissue microarray; PBMC, peripheral blood mononuclear cell; SHM, somatic hypermutation; TF, transcription factor; mTEC, medullary thymic epithelial cell. Figures were created using BioRender.com. (B) Conceptual framework for defining cellular landscapes across thymic, RLN, and tonsil tissues. Figures were created using BioRender.com. (C) UMAP visualization of thymus, RLN, and tonsil samples. Cells are colored by tissue of origin (left) and major cell lineages (right), including T cells, B cells, myeloid cells, stromal cells, and epithelial cells. (D) Proportions of major cell populations in the thymus, RLN, and tonsil samples. (E) UMAP visualization of B cells subsetted from thymus, RLN and tonsil samples, colored by B-cell identity (top) and tissue origin (bottom). Zoomed-in regions highlight a rare B-cell population (CCL17^+^ thymic B) selectively observed in the thymus among the tissues examined. (F) Heatmap showing the canonical marker expression in each B-cell cluster identified by snRNA-seq of thymus, RLN and tonsil samples. Expression values are scaled across clusters (z-score), with red indicating higher and blue indicating lower relative expression. (G) Relative composition of B-cell subsets in RLN, thymus and tonsil samples. Stacked bar plots showing the proportion of B-cell subclusters in each tissue. (H) Quantification of the proportion of CCL17⁺ thymic B cells across thymus, RLN, and tonsil samples.

First, we analyzed single-nucleus RNA sequencing (snRNA-seq) profiles from the FFPE thymus samples (n = 5) to define the cellular composition in contrast to RLN (n = 5) and tonsil controls (n = 4) (Figure 1B). After stringent quality control, 52,343 high-quality nuclei were retained for downstream analysis. Following batch correction, unsupervised clustering resolved five major cell lineages, including T cells, B cells, myeloid cells, stromal cells, and epithelial cells, based on canonical marker expression patterns^24^ (Figures 1C-D and S2A-C). Next, to resolve the heterogeneity within the B-cell compartment, we then subclustered B-lineage cells and annotated the resulting clusters using canonical markers^31^. This approach identified multiple transcriptionally distinct B-cell clusters corresponding to canonical naïve B cells, memory B cells, GC B cells, and plasma cells (Figures 1E-F). While most clusters contained cells from thymus, RLN, and tonsil samples, one cluster characterized by high *CCL17* expression was restricted to the thymus and comprised a median of 8.0% of thymic B cells per sample (range, 5.4–15.2%; Figures 1G–H). This population also expressed *CCL22*, *CD40*, and *CD80* (Figure 1F), resembling the phenotype of B cells previously described as similar to activated dendritic cells^22,30^. To validate the existence of this population, we re-analyzed the B-cell population from a previously published single-cell dataset of the human thymus^24^. Using the top transcriptional markers (*CCL17*, *CCL22*, *ICAM1* [encoding the surface protein CD54], *CD58*, *IL13RA1,* and *IL4I1*) identified in our dataset, we detected a phenotypically similar subset in this external cohort (Figures S3A-B).

### Construction of an integrated B-cell atlas and identification of a thymic B-cell–like PMBCL state

To validate the CCL17⁺ thymic B-cell state at greater cellular depth, we enriched B cells from fresh thymus specimens and generated a fresh-tissue multi-omics cohort comprising thymus (n = 5), RLN (n = 6), tonsil (n = 5), PMBCL (n = 4), ABC-DLBCL (n = 3), and GCB-DLBCL (n = 3) specimens. We integrated 3′ and 5′ gene-expression profiles from thymus, RLN, and tonsil to characterize and annotate non-malignant B-cell states and subsequently integrated these annotated normal B cells with lymphoma-derived B cells for joint analysis. Matched ATAC and BCR data provided orthogonal epigenetic and clonal information (Figure 2A).

**Figure 2.**
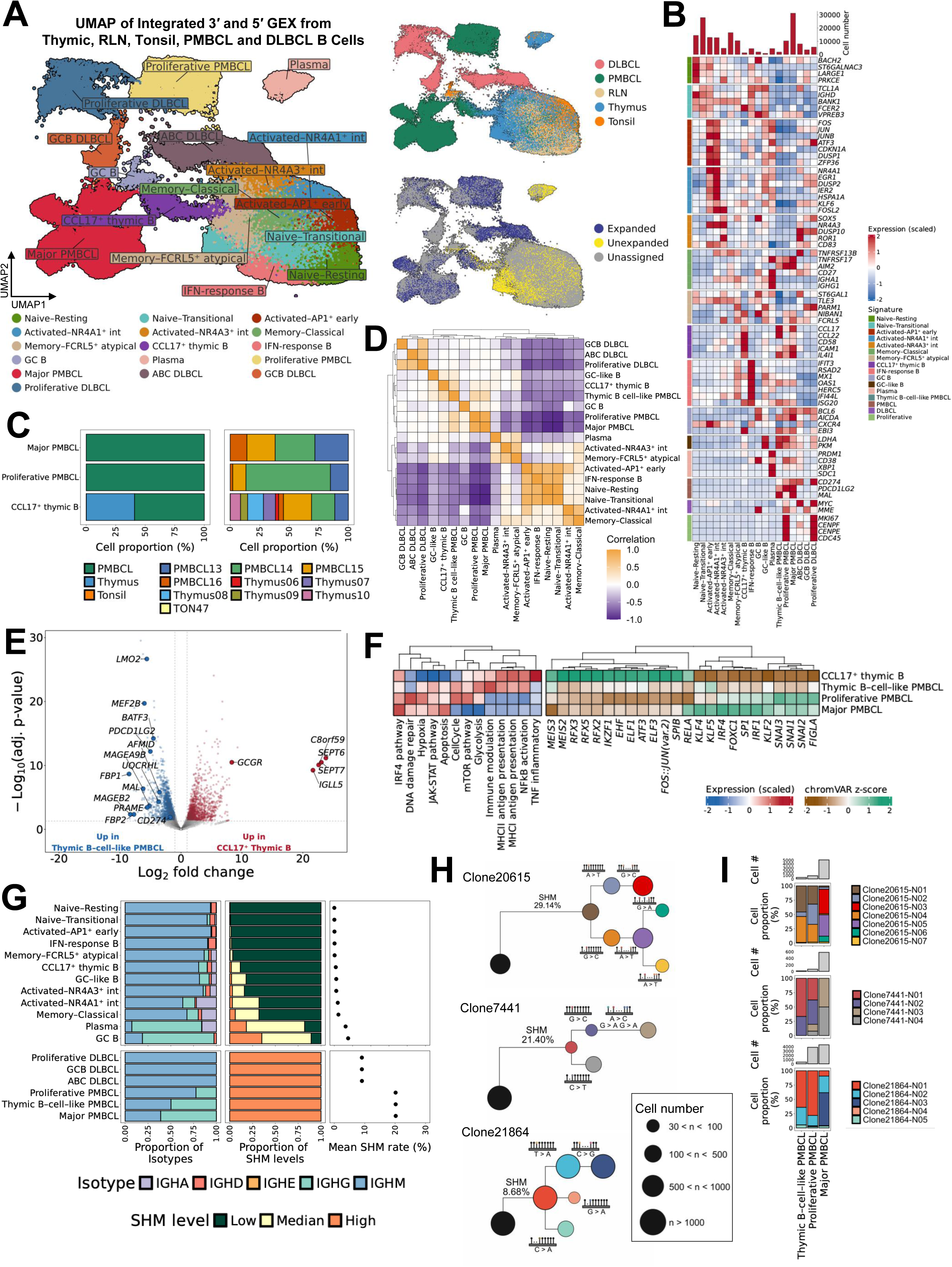
Transcriptional and clonal features of thymic B-cell–like PMBCL. (A) UMAP visualization of integrated gene expression data from single-nucleus multiome (3′ GEX + ATAC) and single-cell (5′ GEX + BCR) profiling of thymus, RLN, tonsil, PMBCL, and DLBCL samples. Left: Each point represents a single cell and is colored by B-cell identity. Right top: Each point represents a single cell and is colored by disease identity. Right bottom: Each point represents a single cell and is colored by clonality. (B) Heatmap showing scaled expression of selected canonical and state-defining markers across B-cell clusters in the integrated atlas. Expression values are scaled across clusters as z-scores, with red indicating higher and blue indicating lower relative expression. (C) Composition of CCL17⁺ thymic and PMBCL B-cell populations across tissues and patients. Left, stacked bar plots showing the proportion of cells derived from samples within CCL17⁺ thymic B cells, proliferative PMBCL, and major PMBCL cells. Right, stacked bar plots showing the contribution of individual samples to each population. (D) Pairwise correlation heatmap of ATAC-derived gene activity profiles across transcriptionally defined normal and malignant B-cell populations in the 3’ snATAC-seq data. Colors indicate correlation coefficients, with orange representing higher similarity and purple representing lower similarity. (E) Volcano plot showing differential gene expression between physiological CCL17⁺ thymic B cells and thymic B-cell–like PMBCL cells. Each point represents a gene, plotted by log₂ fold change and −log₁₀ adjusted *p* value. (F) GSVA and TF motif accessibility analyses comparing physiological CCL17⁺ thymic B cells and the three PMBCL states: thymic B-cell–like, proliferative, and major PMBCL. Left, GSVA heatmap showing relative pathway enrichment across B-cell states. Right, chromVAR heatmap showing motif accessibility z-scores for selected transcription factors. GSVA, gene set variation analysis. (G) Immunoglobulin isotype usage and SHM across normal and malignant B-cell populations. Left, relative proportions of immunoglobulin heavy-chain isotypes (*IGHM*, *IGHD*, *IGHA*, *IGHG*, *IGHE*) across transcriptionally defined B-cell subsets. Middle, distribution of SHM levels classified as Low (mutation rate <1%), Median (1%-5%), or High (> 5%). Right, mean SHM frequency for each population. (H) Phylogenetic trees were reconstructed from single-cell BCR sequences for three representative PMBCL cases (PMBCL14: Clone20615; PMBCL16: Clone7441; PMBCL15: Clone21864). Each node represents a unique BCR sequence, and node size corresponds to the number of cells harboring the identical clonotype (as indicated in the inset of legend). Branch lengths reflect the accumulation of SHM, with SHM frequency (%) indicated relative to the inferred germline sequence. Mutational events are annotated along branches. The black root node represents the inferred germline configuration. (I) Stacked bar plots showing the cellular composition of individual clones across thymic B-cell–like PMBCL, proliferative PMBCL, and major PMBCL cells, illustrating clonal diversification and expansion during tumor evolution.

The normal-tissue B-cell atlas resolved seven major B-cell populations corresponding to naïve, activated, memory, interferon-responsive, GC, and plasma-cell states, together with the CCL17⁺ thymic B-cell state (Figures 2A-B and S4A). The CCL17⁺ population was recovered from all fresh thymus specimens but did not form a comparable population in either RLN or tonsil, independently validating its restriction to the thymus among the non-malignant lymphoid tissues examined (Figure 2C). A subset of B cells from thymus mapped adjacent to conventional GC B cells from RLN and tonsil (Figure S4B). As the thymus lacks conventional GC structures, we operationally refer to these cells as GC-like B cells (Figure S4C). Finer annotation further resolved resting and transitional naïve B cells, AP1⁺ early and NR4A1⁺ or NR4A3⁺ intermediate activated B-cell states, and classical and FCRL5⁺ atypical memory B cells (Figures 2A-B).

Upon integration, lymphoma-derived B cells from different patients converged according to disease entities, forming major PMBCL-, ABC-DLBCL-, and GCB-DLBCL-associated clusters (Figure 2A). Each disease entity additionally contained a distinct, disease-specific proliferative cluster characterized by high expression of *MKI67*, *CENPE*, *CENPF*, and *CDC45* (Figure 2B). In addition to these major and proliferative compartments, a subset of PMBCL-derived B cells co-clustered with physiological CCL17⁺ thymic B cells. We refer to this PMBCL-derived population as the thymic B-cell–like PMBCL state. This shared cluster contained cells from all five thymus donors and all four PMBCL cases, whereas no comparable population was observed in any of the ABC-or GCB-DLBCL cases analyzed (Figures 2A and 2C). Although PMBCL is characterized by a gene-expression profile distinct from conventional DLBCL, two of the three GCB-DLBCL cases analyzed here were unusual in that their bulk transcriptional profiles resembled PMBCL despite arising outside the mediastinum^9,32^. Nevertheless, at single-cell resolution, B cells from these tumors did not co-cluster with physiological CCL17⁺ thymic B cells (Figure 2C), indicating that the thymic B-cell–like PMBCL state is not simply a consequence of an overall PMBCL-like transcriptional program.

We next asked whether the major, proliferative, and thymic B-cell–like PMBCL states were predominantly composed of malignant B cells. To address this, we performed clonotype analysis using scBCR-seq data from three of four PMBCL cases with productive BCR sequences. Across BCR-informative PMBCL cases, cells in all three PMBCL states were almost exclusively derived from expanded B-cell clones, with unrelated unexpanded clonotypes virtually absent, supporting their malignant identity (Figure S4D). Consistent with the clonotype analysis, expression-inferred copy-number profiles revealed patient-specific large-scale chromosomal gains and losses across the major, proliferative, and thymic B-cell–like PMBCL states, providing complementary support for their malignant identity (Figure S4E).

We next asked whether the three PMBCL states defined in the fresh-tissue cohort were reproducible in a larger FFPE snRNA-seq cohort of 13 PMBCL tumors, one of which overlapped with the fresh cohort. All three states, including the major, proliferative, and thymic B-cell–like PMBCL states, were recovered in every case by reference-based label transfer from the fresh-tissue cohort (Figures S5A-E and S6A-C, Table S4).

### Oncogenic rewiring of the thymic B-cell program in PMBCL

Having established the malignant identity of the thymic B-cell–like PMBCL state, we next asked how closely it retained features of physiological CCL17⁺ thymic B cells. Among all states examined, physiological CCL17⁺ thymic B cells were most similar to thymic B-cell–like PMBCL cells at both the transcriptomic (Pearson’s r = 0.518) and chromatin-accessibility (r = 0.506) levels, whereas correlations with the major PMBCL state were more modest (r = 0.252 and 0.280, respectively; Figures 2D and S7A). We next compared the transcriptional programs of physiological CCL17⁺ thymic B cells and thymic B-cell–like PMBCL cells to define features specifically associated with the malignant state (Figure 2E, Table S5). Compared with physiological *CCL17*⁺ thymic B cells, thymic B-cell– like PMBCL cells showed increased expression of oncogenic transcription factors (e.g., *BATF3*, *MEF2B*), metabolic reprogramming genes (e.g., *FBP1*, *FBP2*), immune evasion molecules (e.g., *CD274*, *PDCD1LG2*), and upregulation of cancer-testis antigens, including MAGE family genes and *PRAME* (Figure 2E). These transcriptional changes were further contextualized by gene set variation analysis (GSVA) across normal and malignant clusters. Specifically, physiological CCL17⁺ thymic B cells were enriched for immune-regulatory programs, including TNF signaling, NF-κB activation, and antigen presentation via both MHC class I and class II pathways. The thymic B-cell–like PMBCL cells retained key transcriptional features of their non-malignant counterparts, while additionally showing enrichment of glycolytic and immune modulation pathways. In contrast, major PMBCL cells exhibited marked suppression of antigen presentation programs and were distinguished by enrichment of IRF4/JAK–STAT signaling (Figure 2F). Consistently, chromVAR analysis revealed broader activation of PMBCL-associated transcription factor motif programs, most prominently in major PMBCL cells (Figure 2F).

We next examined BCR isotypes and somatic hypermutation (SHM) levels to further dissect B-cell maturation states across normal and lymphoma subtypes. As expected, naïve B cells showed minimal SHM, whereas classical memory B cells, GC B cells, and plasma cells exhibited progressively higher SHM, consistent with canonical GC maturation. In contrast, physiological CCL17⁺ thymic B cells and GC-like B cells displayed intermediate SHM levels, higher than naïve B cells but lower than classical memory B cells. Notably, thymic B-cell–like PMBCL cells harbored SHM levels comparable to major PMBCL cells, clearly separating them from their physiological counterparts (Figure 2G). To resolve the evolutionary relationships among PMBCL subsets, we reconstructed BCR lineage trees across three independent cases (Figure 2H). To test whether thymic B-cell–like PMBCL cells occupy earlier clonal positions than major PMBCL cells, we mapped cell-state composition onto the BCR phylogeny of each expanded clone and quantified cell-count-weighted node depth from the germline root (Figures 2H–I and S7B). Across all three clones, thymic B-cell– like PMBCL cells occupied earlier phylogenetic positions than major PMBCL cells (mean node depth: 2.02 versus 2.61, 1.33 versus 2.50, and 1.36 versus 1.91, respectively), with a significant difference across clones by within-clone permutation analysis (Δ = 0.77 nodes, *p* = 0.015). Together, these findings associate the thymic B-cell–like PMBCL state with earlier clonal positions and the major PMBCL state with later positions.

### Multimodal dissection of the CCL17⁺ thymic B-cell program reveals its developmental trajectory toward a GC-like B-cell state

Having identified a thymic B-cell–like malignant state in PMBCL, we next examined the physiological thymic B-cell compartment to define the developmental context of the corresponding normal program. We systematically compared B-cell populations across thymus, RLN, and tonsil samples. Quantification of B-cell subsets revealed a pronounced divergence in tissue composition, with the thymus enriched for FCRL5⁺ atypical memory B cells (versus RLN, adjusted *p* < 0.01; versus tonsil, adjusted *p* < 0.01) and plasma cells (versus RLN, adjusted *p* < 0.01; versus tonsil, adjusted *p* < 0.05; Figure 3A). These differences indicated that the thymic microenvironment sustained B-cell states that were underrepresented in the secondary lymphoid organs, suggesting a fundamentally distinct developmental landscape. Integrated transcriptional and chromatin analyses revealed that each thymic B-cell subset was governed by a distinct and internally coherent regulatory program (Figure 3B). While canonical GC B cells exhibited classical GC hallmarks, including robust *AICDA* and *BCL6* expression, GC-like B cells in the thymus are only partially engaged in this program, showing attenuated GC signatures and minimal *AICDA* activation. In addition, CCL17⁺ thymic B cells were defined by a distinct NF-κB-centered regulatory architecture, with coordinated upregulation of inflammatory and antigen-presentation pathways, including *CCL17*, *CCL22*, *CD40*, *CD80*, *NFKB1*, and *NFKB2*. This regulatory program was not observed in any B-cell population derived from RLN and tonsil samples, indicating a thymus-imprinted activation state that is distinct from canonical GC differentiation. Consistent with these observations, chromatin accessibility-based motif analysis showed that GC-like B cells exhibit only partial and uncoordinated activation of GC-associated transcription factor networks, whereas CCL17⁺ thymic B cells are dominated by NF-κB motif activity (Figure 3C, Table S6). Gene Ontology (GO) enrichment analysis further revealed that CCL17⁺ thymic B cells are uniquely enriched for NF-κB signaling and antigen processing and presentation pathways, consistent with an immune-interactive, antigen-presenting B-cell state (Figure 3D, Table S7).

**Figure 3.**
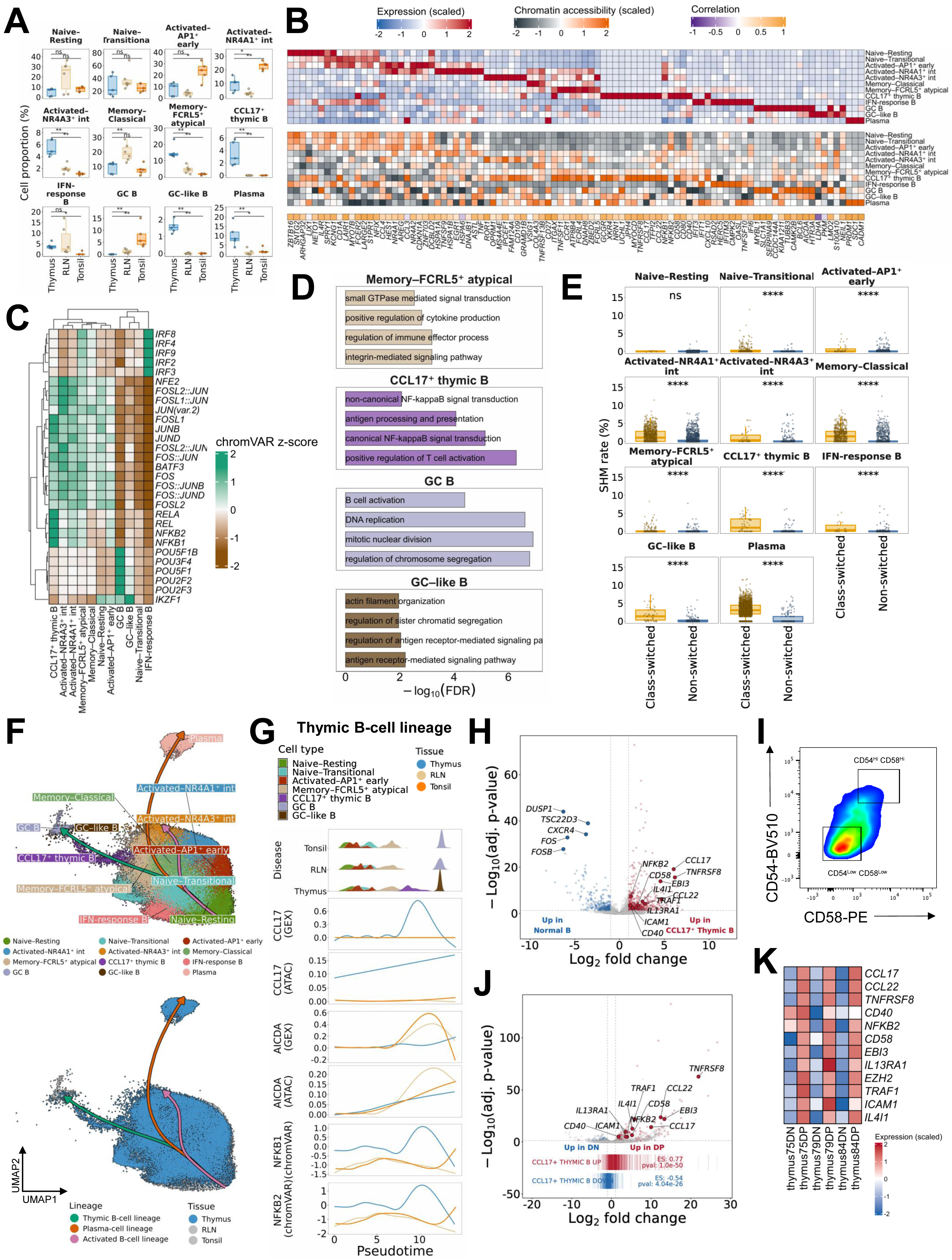
Epigenetic and developmental features of thymic B cells. (A) Boxplots showing the proportion of each transcriptionally defined B-cell subset in thymus, RLN and tonsil samples. Each point represents an individual sample. Statistical significance was assessed using two-sided Mann–Whitney U tests with Benjamini– Hochberg correction for multiple comparisons. Benjamini–Hochberg-adjusted *p* < 0.05 is indicated by (*), adjusted *p* < 0.01 by (**), and ns denotes not significant. (B) Heatmap showing scaled RNA expression (top) and corresponding chromatin accessibility (middle) and correlation (bottom) for selected regulatory and lineage-defining genes across transcriptionally defined thymic B-cell subsets. (C) Heatmap showing chromVAR-derived motif accessibility (Z-scores) for selected transcription factors across normal B-cell subsets. Colors indicate motif accessibility, with green representing higher motif accessibility and brown representing lower motif accessibility. (D) Gene Ontology (GO) enrichment analysis showing significantly enriched biological processes across normal B-cell subsets. (E) Distribution of SHM frequencies in class-switched (IGHA/IGHG/IGHE) versus non-class-switched (IGHM/IGHD) cells across thymic B-cell populations. Statistical significance was assessed using two-sided Mann–Whitney U tests; *P* < 0.0001 is indicated by four asterisks (****). (F) UMAP visualization of normal B-cell populations in thymus samples overlaid with inferred differentiation trajectories reconstructed by pseudotime analysis. (G) Dynamic transcriptional and epigenetic changes along normal B-cell pseudotime. Top, density distribution of thymic, RLN and tonsil B-cell populations along pseudotime, colored by cell type. Middle and bottom panels show smoothed trends of gene expression and chromatin accessibility for *CCL17* and *AICDA* and *NFKB1*/*NFKB2* TF activity along pseudotime. (H) Volcano plot showing differential gene expression between physiological CCL17⁺ thymic B cells and other normal B cells. Each point represents a gene, plotted by log₂ fold change and −log₁₀ adjusted *p* value. (I) Representative flow cytometry density plot showing CD54 and CD58 expression. Cells are color-coded by event density (red to blue, low to high). Gated regions define CD54^low^CD58^low^ DN cells and CD54^high^CD58^high^ DP cells. Numbers indicate the percentage of total cells within each gate. DN, double-negative; DP, double-positive. (J) Volcano plot showing differential gene expression between sorted CD54highCD58high and CD54lowCD58low thymic B cells, together with GSEA showing enrichment of the CCL17⁺ thymic B-cell program in the CD54highCD58high fraction (enrichment score (ES) = 0.77, *P* = 1.0 × 10⁻⁵⁰). GSEA, gene set enrichment analysis. (K) Heatmap showing expression of genes defining single-cell-derived thymic B-cell signatures in sorted CD54highCD58high and CD54lowCD58low B-cell fractions. Expression values are scaled across rows (z-score), with red indicating higher and blue indicating lower relative expression.

Analysis of SHM patterns further delineated the differentiation context of thymic B-cell states (Figure 3E). Substantial SHM was observed specifically within class-switched (*IGHA*⁺ and *IGHG*⁺) CCL17⁺ thymic B cells and GC-like B cells, indicating that SHM preferentially occurs in class-switched thymic B-cell subsets (Figures 3E and S7C). Pseudotime reconstruction further demonstrated that thymic B-cell differentiation follows trajectories distinct from canonical RLN and tonsil GC maturation. One major branch (thymic B-cell lineage) linked CCL17⁺ thymic B cells with GC-like B cells and was characterized by a peak of NF-κB TF activity and *CCL17* accessibility that was not observed in RLN-or tonsil-derived cells (Figures 3F-G and S7D). In contrast, *AICDA* expression and chromatin accessibility steadily increased at late pseudotime stages, indicating that thymic B cells can engage in *AICDA*-associated programs at advanced differentiation stages, without adopting a canonical GC program (Figures 3F-G and S7D).

Finally, we isolated CCL17⁺ thymic B cells from normal human thymus tissue using CD54 and CD58, two surface molecules robustly and consistently upregulated in this B-cell population (Figures 3H-I). Gene set enrichment analysis (GSEA) confirmed that the CD20^+^CD38^-^CD54^high^CD58^high^ fraction is selectively enriched for the CCL17⁺ thymic B-cell transcriptional program (ES = 0.77, *p* = 1.0 × 10^-50^), whereas the CD54^low^CD58^low^ fraction exhibited the opposing gene signature (ES = –0.54, *p* = 4.04 × 10^-26^; Figures 3J–K), supporting the existence of a transcriptionally distinct thymic B-cell population that diverges from canonical GC transcriptional programs.

### CD40L and IL-4 signaling sequentially reconstitute the CCL17⁺ to GC-like B-cell trajectory

To delineate the upstream regulatory signals underlying the NF-κB–centered CCL17⁺ thymic B-cell and GC-like B-cell states, we first performed Ingenuity Pathway Analysis (IPA) on the transcriptional profiles of each subset. This analysis predicted a convergent set of upstream activators, including CD40L, IL-4, TLR4, and NF-κB, as the top predicted upstream regulators of CCL17⁺ thymic B-cell and GC-like B-cell transcriptional states (Figure 4A, Table S8). CytoSig^33^, a cytokine-specific signaling inference framework, independently corroborated these findings and highlighted CD40L and IL-4 as top predicted inducers (Figure 4A, Table S9). We also compared baseline receptor expression across lymphoma cell lines. Consistent with the availability of these signaling axes, PMBCL cell lines showed *CD40* expression comparable to ABC-DLBCL, whereas expression levels of *IL4R* and *IL2RG* were higher than in ABC-and GCB-DLBCL cell lines (Figure S8A). Based on these predictions, we stimulated peripheral blood mononuclear cell (PBMC)-derived CD19⁺ B cells with CD40L, IL-4, LPS (a TLR4 agonist), and BCR engagement via F(ab′)₂ anti-IgM + IgG, administered either individually or in selected combinations (Figure 4B). IFN-γ was included to assess FCRL5-associated programs (Figures 4A–B). Transcriptional profiling revealed that CD40L alone or CD40L + IL-4 induced a gene-expression program closely resembling that of CCL17⁺ thymic B cells. This included robust upregulation of hallmark genes such as *CCL17*, *CCL22*, *IL4I1*, *TRAF1*, *TNFRSF8*, *EBI3*, *NFKB1*, and *NFKB2*. In contrast, stimulation with F(ab′)₂ anti-IgM + IgG or LPS, both of which activate NF-κB, or IL-4 alone, generated only minimal induction of these genes (Figure 4C). To systematically assess cytokine responsiveness, we generated subset-specific transcriptional scores capturing CCL17⁺ thymic B-cell and GC-like B-cell states (termed CCL17^+^ thymic B and GC-like B-cell signature score, respectively) based on the signatures defined by our single-cell B-cell atlas (Tables S10-11) and applied them to each stimulation condition. Both scores were markedly increased following CD40L + IL-4 stimulation, indicating that this combination can induce not only the CCL17⁺-like phenotype (Dunnett-adjusted *p* < 0.001 for all comparisons) but also a GC-like B-cell transcriptional state (Dunnett-adjusted *p* < 0.001 for all comparisons; Figure 4D). The CCL17⁺ thymic B signature score peaked at 24 hours, whereas the GC-like B-cell score gradually increased and reached a maximum at 96–120 hours (Figure 4E), suggesting a temporal progression from a CCL17⁺-like to a GC-like B-cell phenotype using the same cytokine cocktail. These results support a model in which CCL17⁺ and GC-like B cells lie along a shared trajectory recapitulated by CD40L and IL-4 signaling. Notably, addition of F(ab′)₂ anti-IgM + IgG to CD40L and IL-4 stimulation significantly attenuated the CCL17⁺ thymic B-cell program and showed a trend toward reduced GC-like signature scores (Figure S8B). We next investigated whether the CCL17⁺ and GC-like B-cell programs can arise from distinct peripheral B-cell compartments. Stimulation of purified naïve and memory B cells from peripheral blood with CD40L and IL-4 revealed that both subsets were capable of generating a CCL17⁺-like phenotype, although with different kinetics (Figure S8C), indicating that the CCL17⁺ program is not restricted to thymic B cells but can be recapitulated in peripheral counterparts when exposed to the same receptor stimulation.

**Figure 4.**
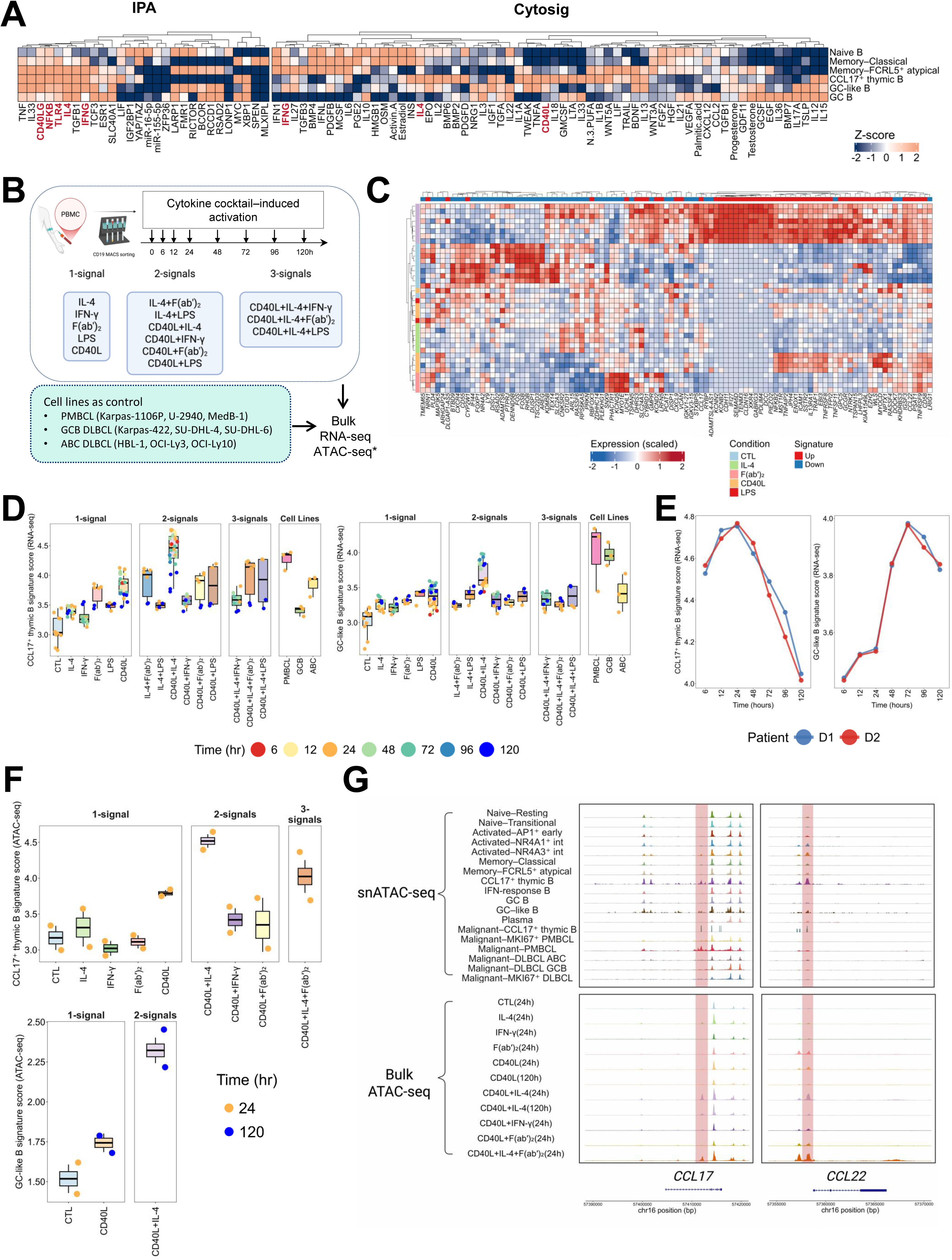
*In vitro* modeling of cytokine-driven induction of the thymic B-cell program. (A) Left: IPA identifying predicted upstream regulators of the CCL17⁺ thymic B-cell and GC-like B-cell transcriptional programs. CD40LG, NFKB, IL4, IFNG and TLR4 are highlighted as dominant predicted drivers. Right: Cytokine signaling inference using CytoSig independently corroborating IPA predictions and identifying CD40L and IL4 as top inducers of CCL17⁺ and GC-like B-cell transcriptional states. IPA, Ingenuity Pathway Analysis. (B) Experimental design for in vitro stimulation of PBMC-derived CD19⁺ B cells. BCR engagement was achieved using F(ab′)₂ anti-IgM + IgG. **\*** Bulk ATAC-seq was performed for selected stimulation conditions at 24 and 120 hours, as indicated in panel F. (C) Heatmap showing scaled expression of CCL17⁺ thymic B-cell signature genes 24 h after in vitro stimulation of PBMC-derived CD19⁺ B cells with control (CTL), IL-4, CD40L, CD40L + IL-4, LPS, or BCR engagement using F(ab′)₂ anti-IgM + IgG. Genes are hierarchically clustered and annotated according to signature direction (up-or downregulated), and columns are annotated by stimulation condition. (D) Left: CCL17⁺ thymic B-cell signature scores across stimulation conditions and time points following *in vitro* activation of PBMC-derived CD19⁺ B cells. Right: GC-like B-cell signature scores across the same stimulation conditions and time points. Boxplots summarize distributions across time points, with individual points representing biological replicates. Lymphoma cell lines are shown as transcriptional reference controls. (E) Line plots showing the kinetics of CCL17⁺ thymic B-cell (left) and GC-like B-cell (right) signature scores following combined CD40L + IL-4 stimulation of PBMC-derived CD19⁺ B cells. Lines represent independent donors. (F) Top: CCL17⁺ thymic B-cell signature scores calculated from ATAC-seq data across one-, two-, and three-signal stimulation conditions. Bottom: GC-like B-cell signature scores calculated from ATAC-seq data across one-, two-, and three-signal stimulation conditions. Boxplots summarize distributions across replicates, with points representing individual samples. (G) Genome browser tracks showing chromatin accessibility at the *CCL17* (left), and *CCL22* (right) loci. Top panels display aggregated snATAC-seq profiles across thymic B-cell subsets and malignant B-cell populations. Bottom panels show bulk ATAC-seq profiles of PBMC-derived B cells following cytokine stimulation under indicated conditions and time points.

To assess whether induced states recapitulate thymic epigenetic programs, we performed ATAC-seq on samples collected at 24-and 120-hour post-stimulation. Chromatin-accessibility signatures were consistent with a CCL17⁺ thymic B-cell program at 24 hours and a GC-like program at 120 hours (Figure 4F, Tables S12-13). Locus-level analysis supported this progression: at 24 hours, NF-κB–associated regulatory elements at the *CCL17* promoter and *CCL22* enhancer were accessible, mirroring CCL17⁺ thymic B cells (Figure 4G), while at 120 hours, the *AICDA* locus showed GC-like chromatin accessibility (Figure S8D). Together, these results indicate that CD40L and IL-4 stimulation sequentially reconstitute the epigenetic transition from the CCL17⁺ to the GC-like B-cell state, recapitulating the thymic trajectory *in vitro*.

### Spatial transcriptomic profiling identifies a medullary microenvironment enriched in CD40L⁺ and IL-4⁺ cells surrounding CCL17⁺ thymic B cells

To define the spatial organization of thymic B-cell subsets, we performed 6,175-plex CosMx spatial molecular imaging with an additional 200 customized add-on genes (total 6,375 genes) (Table S14) on 30 FFPE human thymus samples (0–18 years) and obtained 741,248 cells after stringent quality control. Label transfer from matched FFPE snRNA-seq enabled robust annotation of immune and stromal populations, including rare thymic B-cell subsets, with consistent cell-type proportions across donors (Figure S9A). To enable cohort-wide spatial analyses, all cells were assigned to thymic anatomical regions (Figures 5A–B). Spatial mapping showed that CCL17⁺ thymic B cells and GC-like B cells localized almos exclusively to the medulla, with minimal presence in the cortex or Hassall’s corpuscles (Figure 5B). Other B-cell subsets showed broader spatial distributions, whereas CCL17⁺ thymic B cells were reproducibly detected across donors (Figures 5B–C).

**Figure 5.**
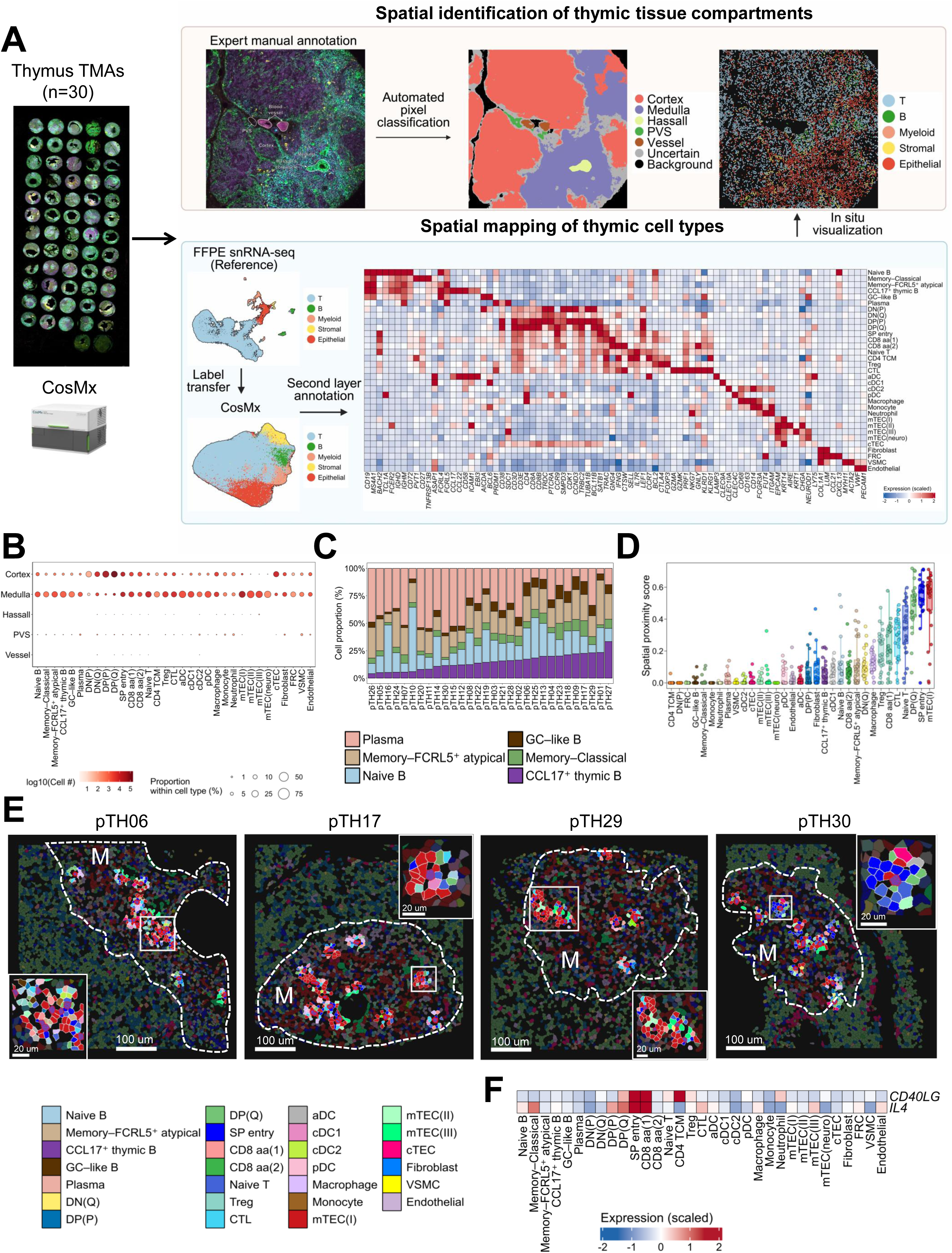
Spatial transcriptomic mapping of thymic B-cell states. (A) Overview of the analytical workflow used to define the spatial organization of thymic B-cell subsets. Top, expert manual annotation of major thymic anatomical regions on representative immunofluorescence images was used to train a supervised pixel-level classifier, enabling automated and standardized region assignment across all CosMx FOVs, followed by in situ visualization. Bottom, FFPE snRNA-seq data served as a reference for label transfer to CosMx spatial transcriptomic profiles, enabling robust annotation of immune and stromal cell populations and second-layer refinement of rare thymic B-cell subsets. This integrated framework enabled high-resolution mapping of cell identities and spatial contexts across 30 FFPE human thymus samples. FOV, field of view. Figures were created using BioRender.com. (B) Spatial distribution of thymic cell types across annotated anatomical regions (cortex, medulla, Hassall’s corpuscles, PVS, and vessels). Dot size indicates the proportion of each cell type within a given region, and color intensity represents the log-transformed absolute cell number. PVS, perivascular space. aDC, activated dendritic cell. cDC, conventional dendritic cell. (C) Proportions of B-cell subsets across individual thymus donors profiled by CosMx spatial molecular imaging. Stacked bars show the relative contribution of each B-cell population per donor. (D) Spatial proximity analysis centered on CCL17⁺ thymic B cells. Boxplots show proximity scores for neighboring cell types within a 50 μm radius. Hgher scores indicate closer local proximity to CCL17⁺ thymic B cells. (E) Representative CosMx spatial molecular imaging section illustrating the spatial distribution of annotated thymic cell types (scale bar, 100 µm) in four individual cases. Individual segmented cells are colored by cell identity. CCL17⁺ thymic B cells are highlighted (outlined in red) and all cells located within a 50 µm radius of these cells are indicated. M denotes the medulla. Insets depict magnified views of the boxed regions (scale bar, 20 µm). (F) Heatmap showing scaled expression of *CD40LG* and *IL4* across thymic cell populations within the niche of CCL17⁺ thymic B cells. Expression values are scaled across clusters (z-score), with red indicating higher and blue indicating lower relative expression.

Spatial proximity analysis^34^ centered on CCL17⁺ thymic B cells identified mTEC(I), SP-entry T cells, and quiescent DP (Q) thymocytes among the closest neighboring populations (Figures 5D–E, S9B). Neighboring cells with CCL17⁺ thymic B cells exhibited increased expression of *CD40LG*, *IL-4*, and chemokine ligands (*CXCL12*, *CXCL13*, *CCL21*, *CCL22*), indicating a cytokine-rich, NF-κB–associated niche (Figure S9C). To further resolve the cellular sources of these signals within the niche, we examined lineage-specific expression patterns. *CD40LG* was mainly expressed by CD4 central memory T cells (TCM), SP-entry, and CD8 αα (1) T cells, whereas *IL4* originated primarily from DP (Q), CD8 αα (1) and SP-entry subsets (Figure 5F). Together, these data define a medullary CD40L and IL-4 signaling niche associated with the CCL17⁺ program.

### PMBCL B-cell states occupy a microenvironment distinct from the physiological thymic niche

To determine whether the cellular niches associated with physiological thymic B cells were preserved in PMBCL, we performed CosMx spatial molecular imaging on tissue microarrays from 68 PMBCL cases, all derived from biopsies of mediastinal lesions, and obtained 432,747 cells after stringent quality control. Reference-guided annotation identified malignant B cells and accompanying lymphoid, myeloid, stromal, and epithelial populations (Figure 6A). Compared with normal thymus, PMBCL showed increased representation of myeloid and stromal populations and reduced representation of T cells, non-malignant B cells, and epithelial cells, together with a prominent malignant B-cell compartment (Figures 6B and S10A). A similar shift in cellular composition was also observed in the FFPE snRNA-seq cohort (Figures S10B-C). Notably, TECs were rarely detected in profiled tumor regions. Spatial proximity analysis centered on malignant PMBCL cells showed the strongest proximity to other malignant B cells, followed by IL18-expressing macrophages and interferon-responsive fibroblasts (Figure 6C). This tumor-centered organization contrasted with the epithelial-and thymocyte-associated neighborhoods of physiological CCL17⁺ thymic B cells. Representative spatial maps illustrated these distinct cellular arrangements (Figure 6D).

**Figure 6.**
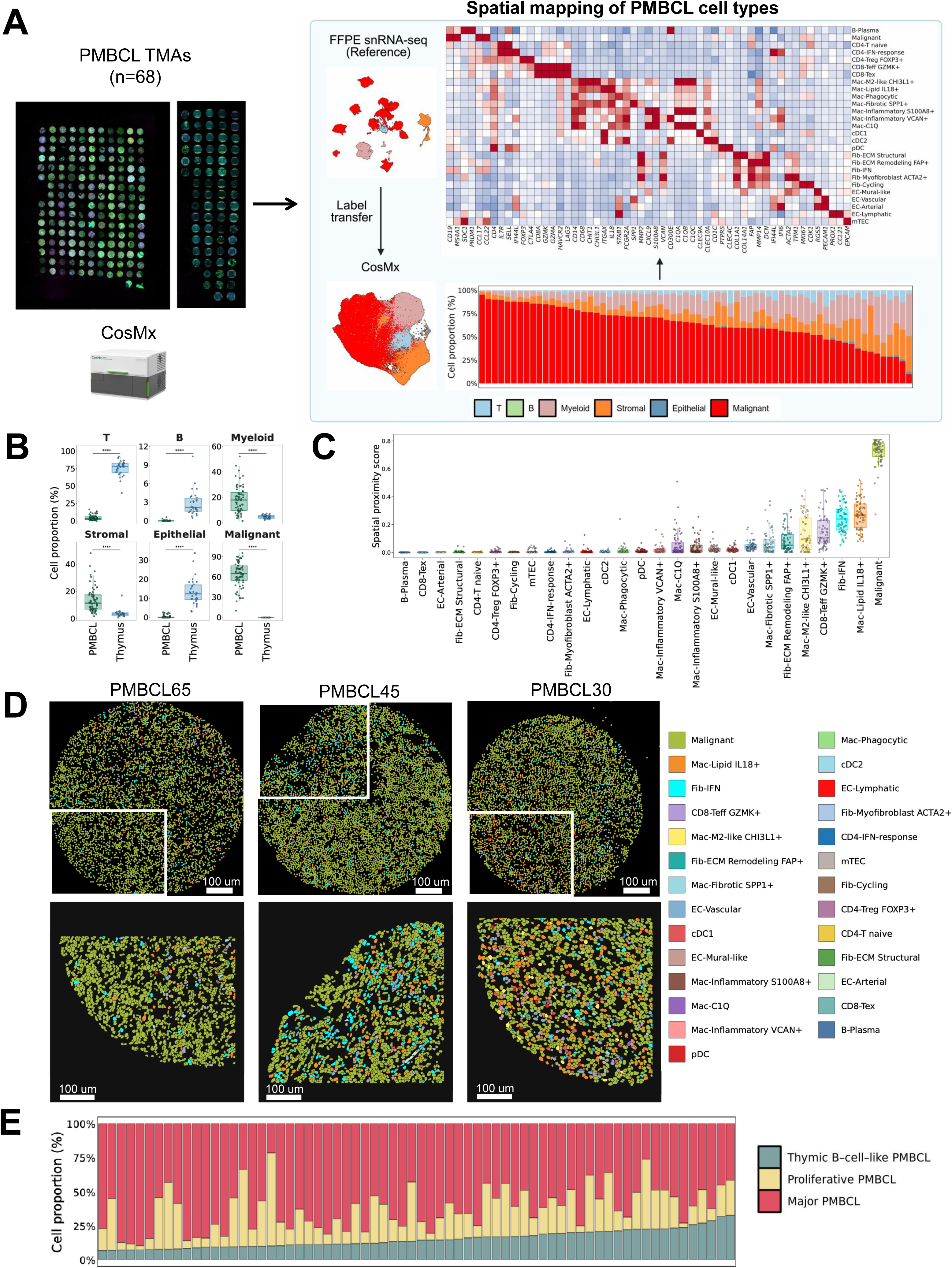
**Spatial transcriptomic mapping of PMBCL malignant B-cell states**. (A) Overview of the analytical workflow used to define the spatial organization of PMBCL samples. Left, overview of PMBCL TMAs. Middle, FFPE snRNA-seq data were used as a reference for label transfer to CosMx profiles, enabling annotation of major immune, stromal, epithelial, and malignant cell populations. Right, cell-type annotations were validated by the expression of lineage-specific marker genes (top), and their relative proportions across PMBCL cores were quantified (bottom). Figures were created using BioRender.com. (B) Boxplots showing the proportion of major cell lineages in thymus, and PMBCL samples. Each point represents an individual sample. Statistical significance was assessed using two-sided Mann–Whitney U tests; * *p* < 0.05, ** *p* < 0.01, *** *p* < 0.001, ** *p* < 0.0001; ns, not significant. (C) Spatial proximity analysis centered on PMBCL malignant B cells. Boxplots show proximity scores for neighboring cell types within a 50 μm radius. Higher scores indicate closer local proximity to PMBCL malignant B cells. (D) Representative CosMx spatial molecular imaging section illustrating the spatial distribution of annotated PMBCL cell types (scale bar, 300 µm) in three individual cases. Individual segmented cells are colored by cell identity. (E) Stacked barplot showing the proportion of malignant B-cell states in PMBCL samples.

Using matched FFPE snRNA-seq data from 13 of the 68 cases, we next mapped the major, proliferative, and thymic B-cell–like PMBCL states in the spatial dataset and subsequently extended these annotations across the full cohort. All three states were identified in every case, although their relative proportions varied among tumors (Figures 6E and S10D). No obvious state-specific differences in local cellular neighborhoods were observed (representative examples in Figure S10E).

## Discussion

In this study, we delineate the molecular organization of a CCL17⁺ thymic B-cell program, its relationship to a GC-like state, and its representation within PMBCL. The physiological CCL17⁺ and GC-like programs can be induced *in vitro* from primary B cells through stimulation with cytokines that are characteristically expressed by specific immune cell subsets in the medulla microenvironment of the thymus. Transcriptional and chromatin-accessibility similarities between physiological CCL17⁺ thymic B cells and the thymic B-cell–like PMBCL state establish a molecular link between a niche-associated CCL17⁺ thymic B-cell program and an inherent malignant cell state in PMBCL. Despite this molecular resemblance, spatial profiling revealed that thymic B-cell–like PMBCL cells reside within a TME markedly different from the physiological thymic niche.

A three-state transcriptional architecture comprising major, proliferative, and thymic B-cell–like PMBCL states was consistently observed across FFPE, fresh tissue, and spatial datasets. In BCR-informative cases, the thymic B-cell–like PMBCL cells were preferentially associated with earlier nodes in reconstructed phylogenies than major PMBCL cells. Combined with the transcriptional differences between these states, this pattern suggests that PMBCL clonal evolution is accompanied by attenuation of antigen-presentation programs and reinforcement of IRF4/JAK–STAT-associated programs.

A key distinction between physiological CCL17⁺ thymic B cells and thymic B-cell– like PMBCL cells lies in their SHM burden. Although SHM is classically associated with GC responses, where AID is the principal driver^35,36^, several studies showed that SHM can also occur in the thymus^28,30,37,38^. In our study, physiological CCL17⁺ and GC-like thymic B cells showed lower SHM burdens than memory B cells and plasma cells. Nevertheless, the parallel increase in *AICDA* expression and SHM along the inferred CCL17⁺–to–GC-like trajectory raises the possibility that this differentiation program provides a permissive window for AID-associated mutational accumulation. The relatively low SHM burden in these physiological states may partly reflect the young age of the thymus donors, consistent with reports that SHM levels increase during childhood^39^. In contrast, thymic B-cell–like PMBCL cells harbored a high SHM burden. This may reflect age-related SHM accumulation^40^, aberrant AID activity during malignant evolution, or both.

Our spatial analyses reveal a thymus-specific architecture in which rare CCL17⁺ B cells reside within T cell-dense medullary zones enriched for multiple cellular sources of CD40L and IL-4. In contrast, RLN contains abundant B cells organized into GCs, where CD40L and IL-4 are provided primarily by a relatively small subset of T follicular helper cells^41,42^. The lack of a comparable CCL17⁺ B-cell state in RLN and tonsil may reflect differences in niche organization and the integration of CD40L/IL-4 signals with BCR signaling input, as suggested by our *in vitro* experiments. In the thymus, the CCL17⁺ program is associated with CD40L-dependent, NF-κB-centered signaling, with IL-4 likely acting as an amplifier rather than a primary driver. In fully established PMBCL, however, the thymic B-cell–like PMBCL cells were detected within a markedly different cellular environment, with few detectable thymic epithelial cells. This raises the possibility that the program can be maintained with reduced dependence on the physiological thymic niche, potentially through cell-intrinsic reinforcement of NF-κB and JAK–STAT signaling by recurrent genetic alterations.

Collectively, our study highlights the human thymus as an active niche that shapes a distinct B-cell developmental program that diverges from canonical GC trajectories, linking normal thymic differentiation to thymic B-cell lymphomagenesis.

## Resource availability Data availability

The raw sequencing data generated in this study will be deposited in the European Genome– phenome Archive (EGA). The processed single-cell data will be deposited at the cellxgene database(https://cellxgene.cziscience.com/). The processed CosMx data will be deposited at zenodo. The data will be made publicly available upon publication.

## Code availability

Scripts used for data analysis are available at https://github.com/LCR-BCCRC/Developmental_Origin_Thymic_Lymphoma and will be available upon publication.

## Acknowledgements

This study is supported by Program Project Grant funding from the Terry Fox Research Institute (Grant No.1061 and 1108), Large Scale Applied Research Project funding from Genome Canada (Grant No. 13124), Genome BC (Grant No. 271LYM) and CIHR (Grant No.GP1-155873, Grant No. 191809), the Canadian Cancer Society Research Institute (Grant No. 705288), a Foundation grant from CIHR (Grant No. 148393), the BC Cancer Foundation and the Paul G. Allen Frontiers Group (Distinguished Investigator award to C.S., Grant No.12829). S.R. was supported by the Randy Gascoyne lymphoma fellowship, and by a physician scientist fellow award from the Leukemia & Lymphoma Society of Canada. M.F. was supported by Gilead’s Research Scholars Program and a grant from The Leukemia & Lymphoma Society (Grant ID: 3442-25). T.A. was supported by a fellowship from CIHR and the Lymphoma Research Foundation. T.A. has received research funding support from V Foundation, Hold‘em life program and Princess Margaret Cancer Foundation. T.A. is the recipient of ASH scholar Award. L.V. was supported by a Michael Smith Health Research Foundation Scholarship SCH-2022-2607, and Canada Foundation for Innovation John R. Evans Leaders Fund #43630 and Innovation Fund #46036 (L.V.).

## Visualization

Parts of the figure were created using BioRender.com.

## Authors’ contributions

Study design: S Rai, Y Yin, M Fujisawa, T Aoki, C Steidl.

Writing: All Authors.

Manuscript review: All Authors.

Data interpretation: S Rai, Y Yin, M Fujisawa, T Aoki, C Sarkozy, C Steidl.

*In vitro* experiments: S Rai, M Fujisawa, M Kishida, M Li.

Data analysis: Y Yin, J Niu, A Jiang, L Hilton, A Fong, G Duns, S Wu, E Lee.

Sequencing processing: S Rai, A Telenius, M Boyle, L ÒBrien, C Cassidy.

Pathological review: A Lytle, P Farinha, G Slack, J Bush.

Case identification: S Rai, Y Yin, M Fujisawa, T Aoki, C Steidl.

Administrative (i.e., reporting or organizing data, constructing databases): A Telenius, M Boyle, H Nakamura, S Daryani, B Meissner, A Roos.

Supervision: R Morin, L Venturutti, A Weng, A Roth. B Nelson, K Savage, T Aoki, C Sarkozy, D Scott, C Steidl.

## Declaration of Generative AI and AI-assisted technologies

During the preparation of this work, the authors used ChatGPT (OpenAI) to improve the language and readability of the manuscript. After using this tool, the authors reviewed and edited the content as needed and took full responsibility for the content of the publication.

## Materials and Methods

### Key Resources Table

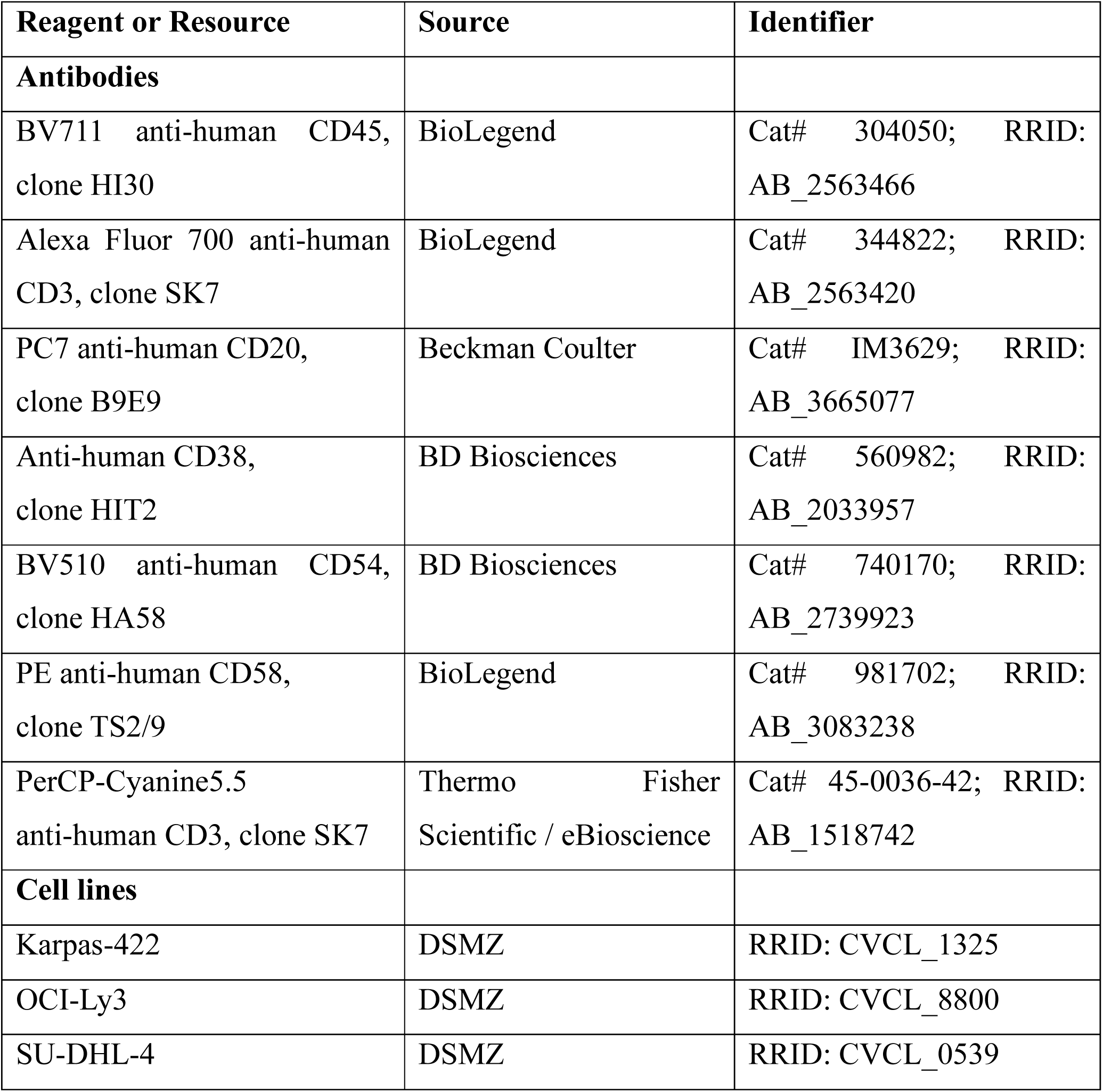
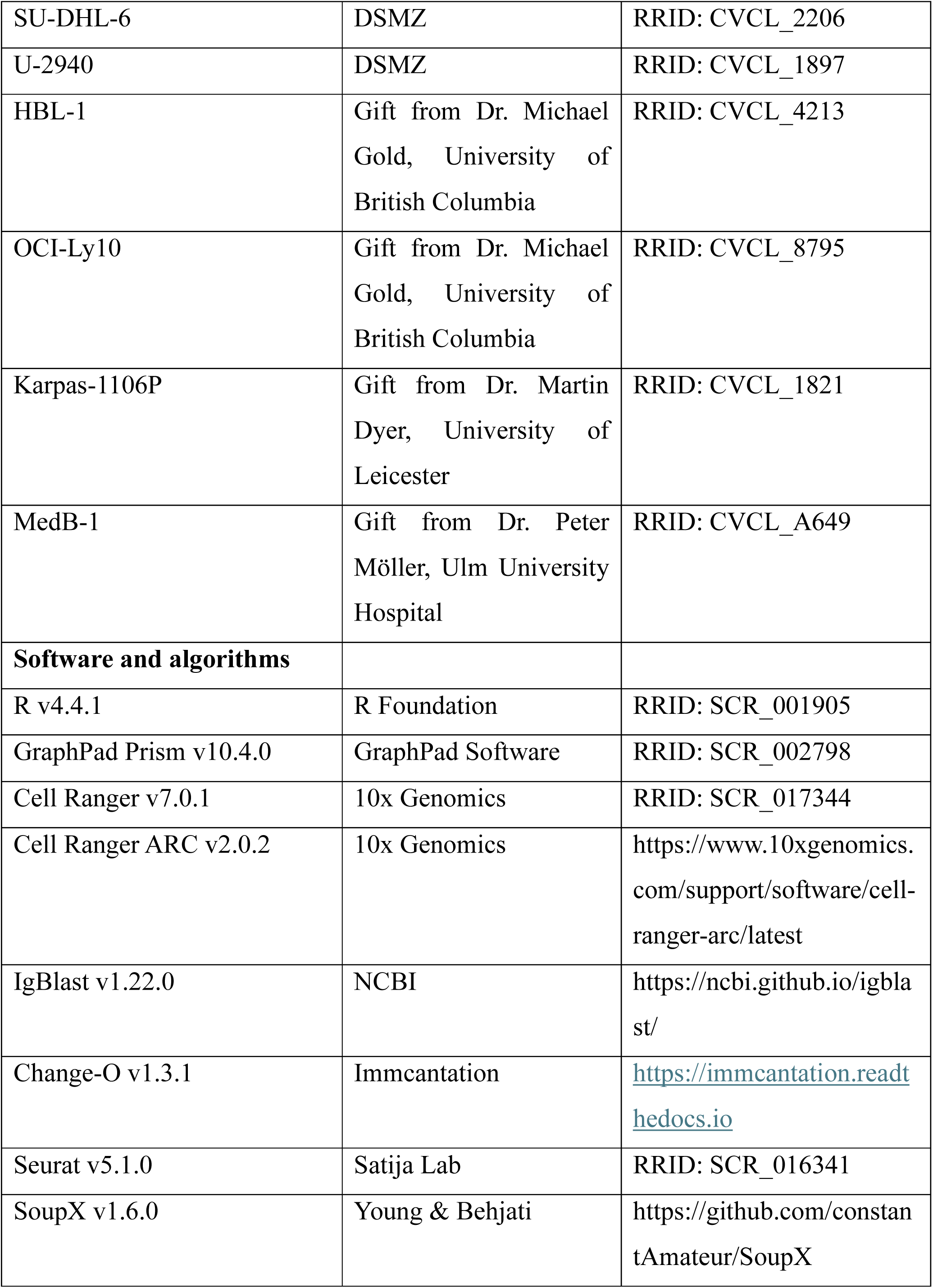
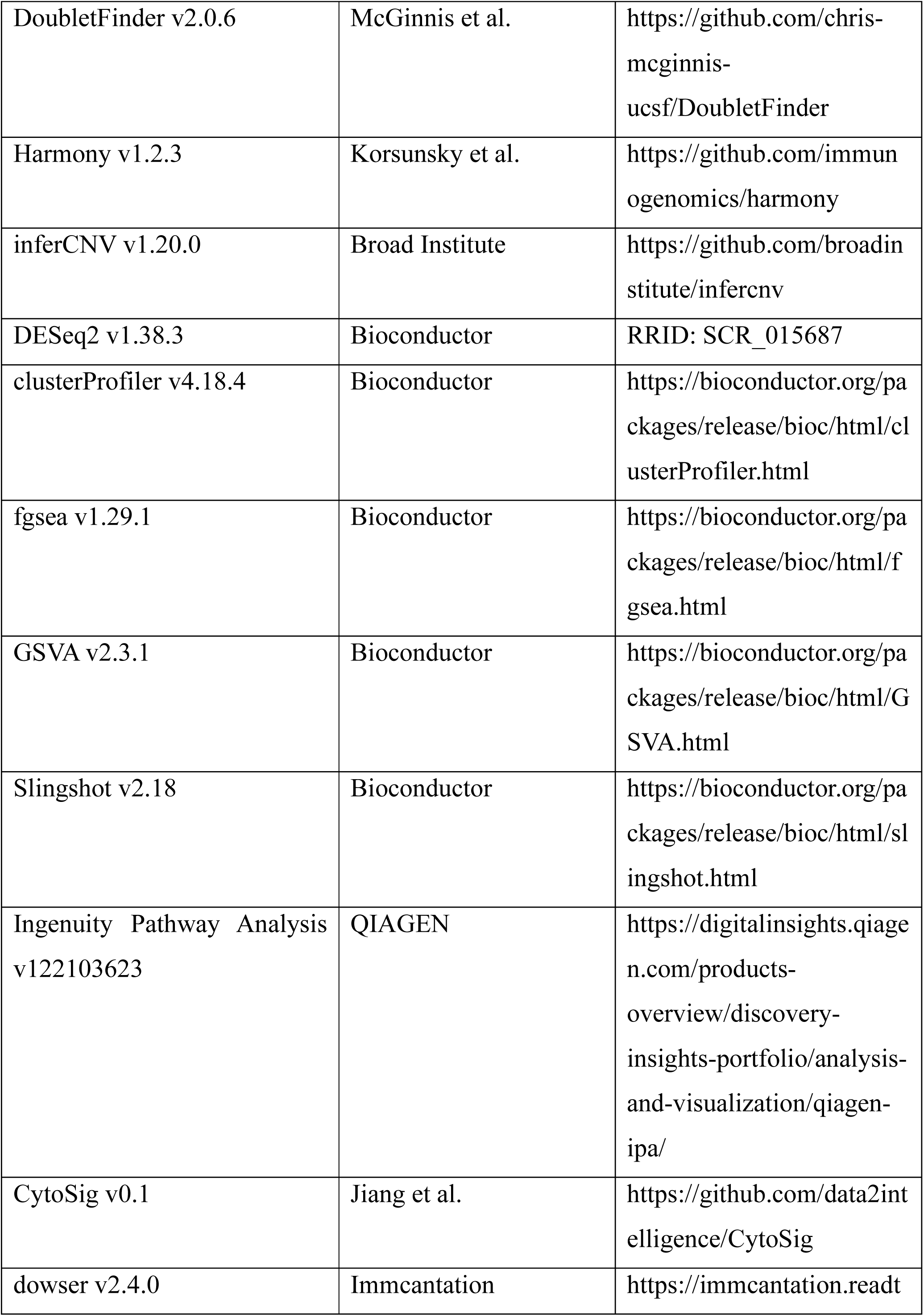
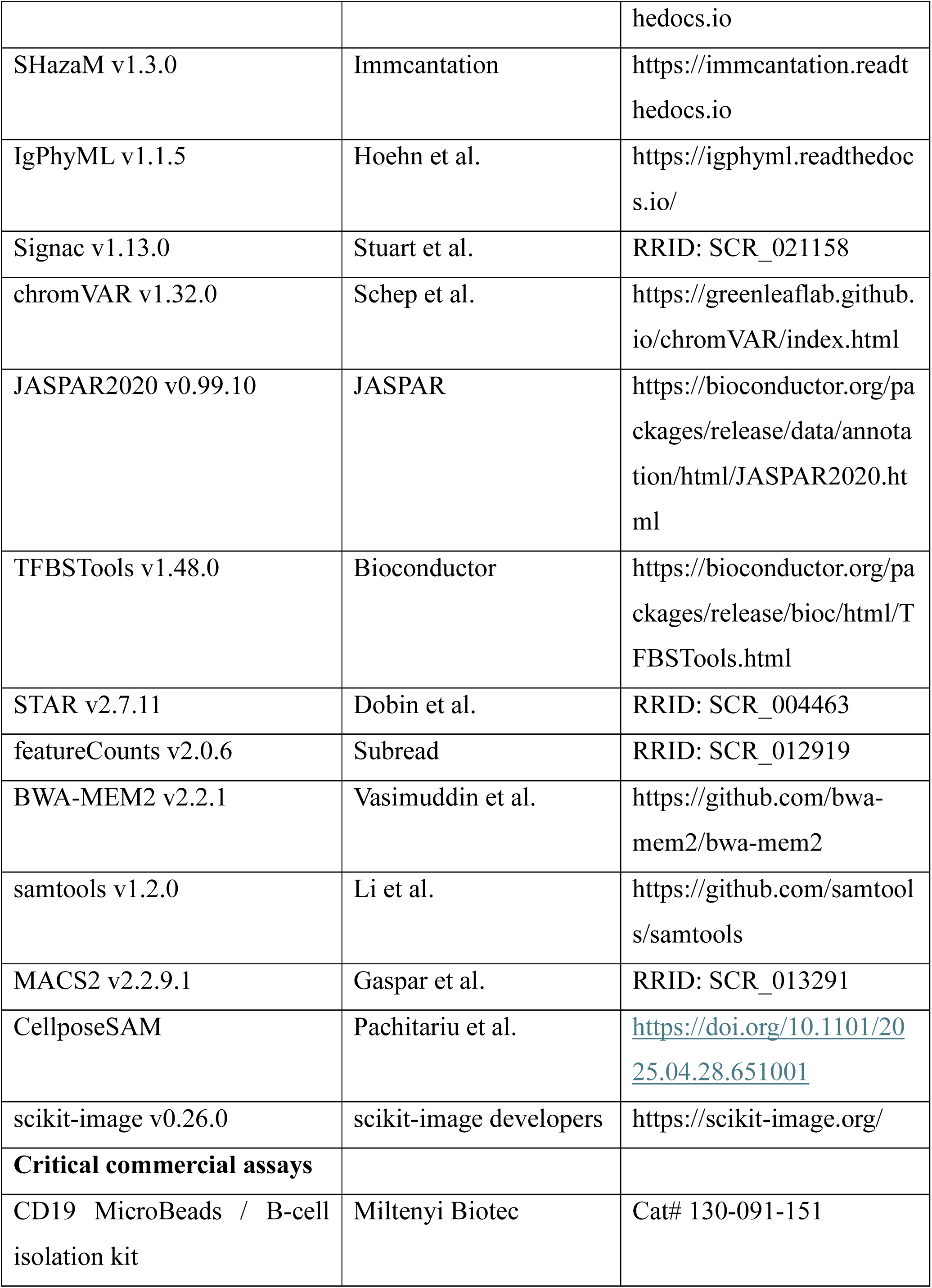
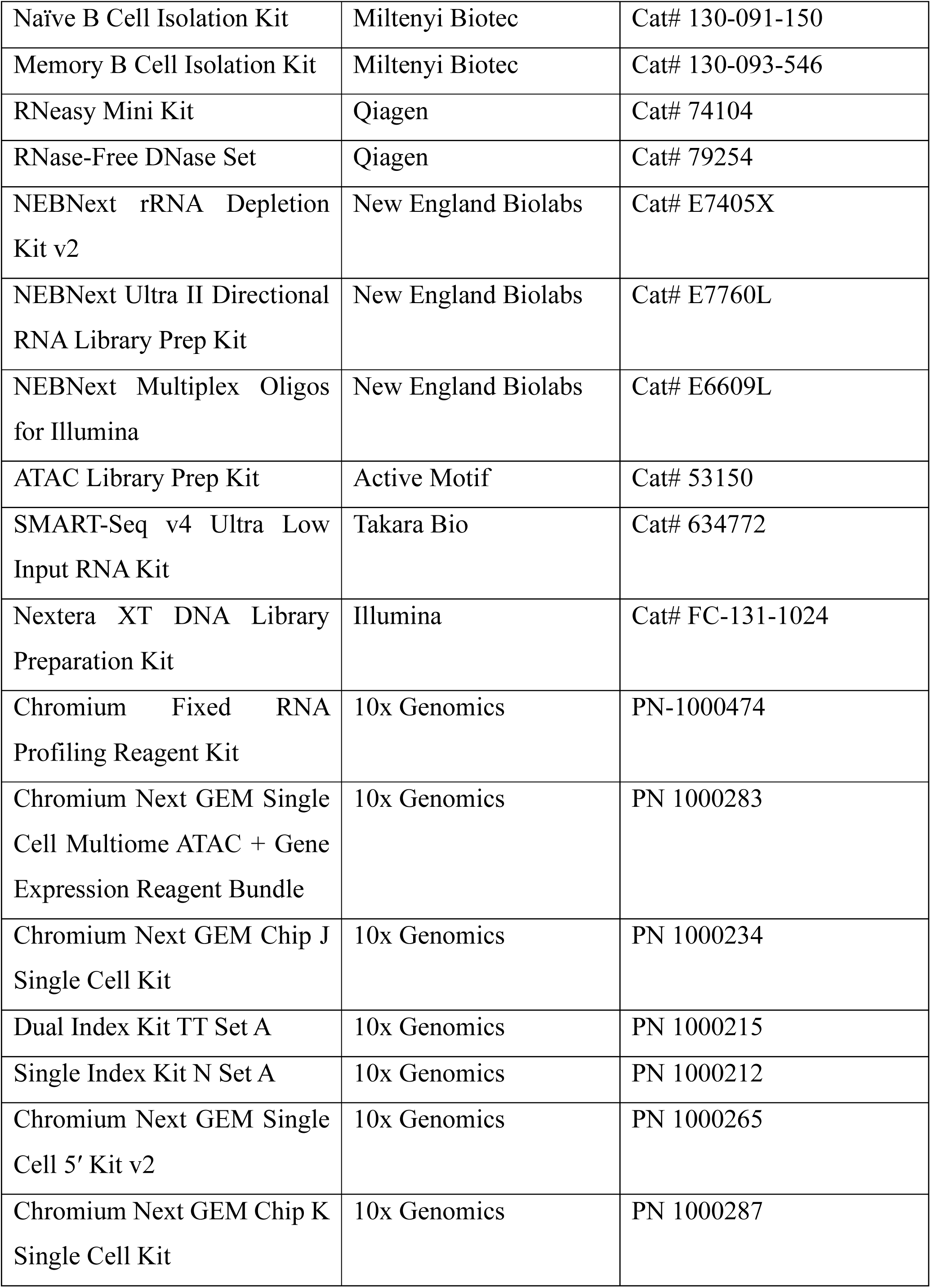
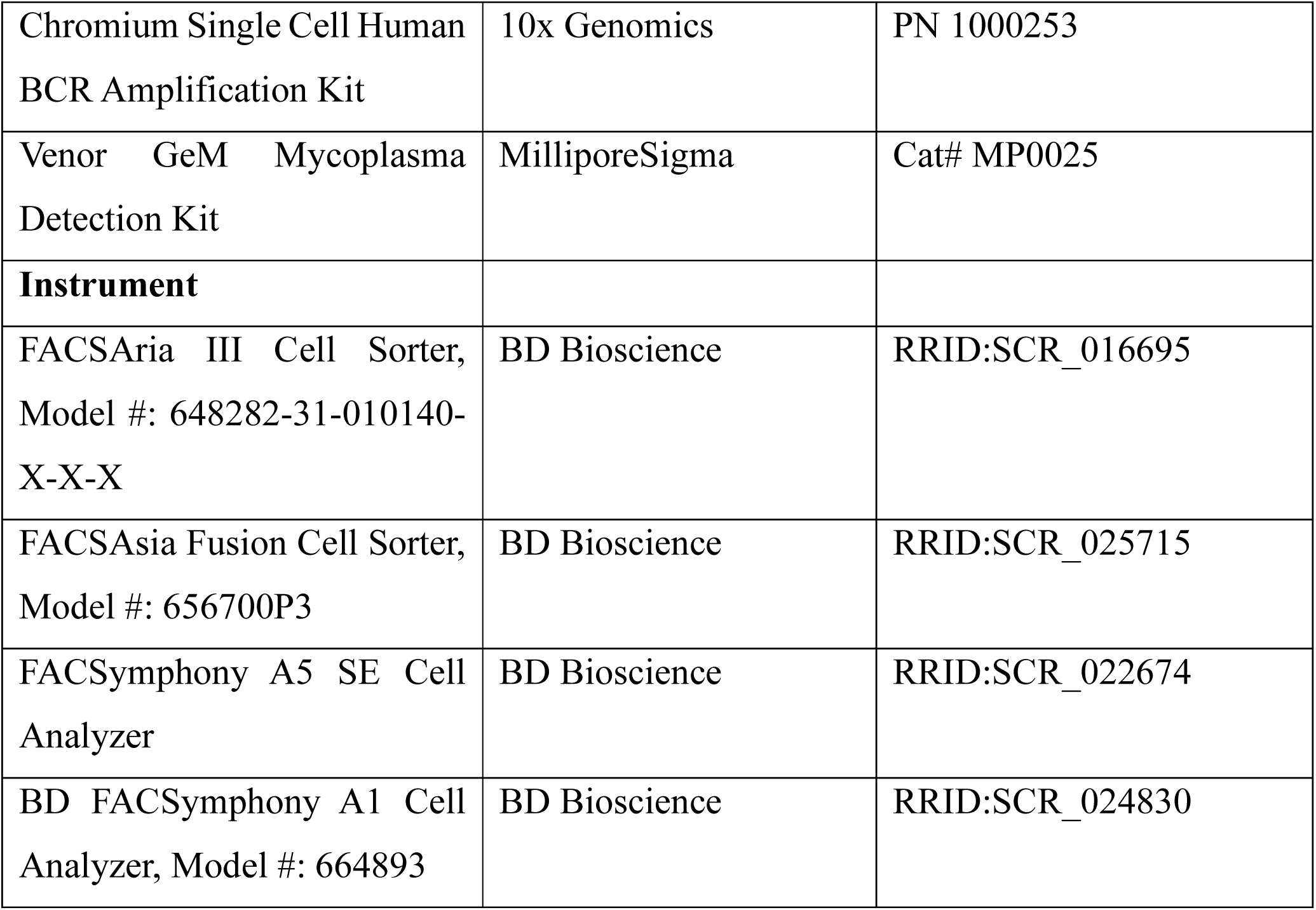

### Study protocols

This study received ethical approval from the University of British Columbia (UBC)–BC Cancer Research Ethics Board (H18-00469, and H22-03661), adhering to the principles of the Declaration of Helsinki. Written informed consent was obtained from patients, or the need for consent was waived by the Research Ethics Board.

### Human samples

Formalin-fixed paraffin-embedded (FFPE) and fresh-frozen reactive lymph node (RLN) samples were retrospectively collected following lymph node resection. RLN samples were defined as lymph nodes showing reactive histopathologic changes without evidence of lymphoma or other lymphoid malignancy on expert hematopathologic review. Normal thymus tissues were obtained from non-neoplastic thymic tissue collected during cardiothoracic surgery, and tonsil samples were obtained from tonsillectomy specimens. All specimens were reviewed by experienced hematopathologists. PMBCL and DLBCL cases were selected from diagnoses established by expert hematopathologists in close multidisciplinary collaboration with lymphoma clinicians, integrating clinical presentation, radiologic findings, histopathologic features, and immunophenotype according to established diagnostic criteria. PMBCL diagnosis required concordant clinical, morphologic, and immunophenotypic features, including a primary mediastinal mass. DLBCL cases were classified as ABC-or GCB-DLBCL using the NanoString-based Lymph3Cx gene-expression assay, which also provides a PMBCL-associated molecular signature.

### Biological materials / sample preparation Cell lines

The following human B-cell lymphoma lines (Karpas-422, OCI-Ly3, SU-DHL-4, SU-DHL-6, U-2940) were purchased from DSMZ (German Collection of Microorganisms and Cell Cultures). HBL-1 and OCI-Ly10 were a kind gift from Dr. Michael Gold (The University of British Columbia, Vancouver, Canada). Karpas-1106P was a kind gift from Dr. Martin Dyer (The University of Leicester, Leicester, UK). Med-B1 was kindly provided by Dr. Peter Möller (Ulm University Hospital, Ulm, Germany). HBL-1, OCI-Ly3 and U-2940 cells were cultured in Roswell Park Memorial Institute-1640 medium (RPMI-1640) supplemented with 10% heat-inactivated Fetal Bovine Serum (FBS) (Corning, catalog no. 35-077-CV). Karpas-1106P, SU-DHL-4, SU-DHL-6 cells were cultured in RPMI-1640 + 20% FBS. OCI-Ly10 was cultured in Iscove’s Modified Dulbecco Medium + 20% FBS. All cell lines were verified through short tandem repeat (STR) profiling conducted by the Centre for Applied Genomics (Toronto, Canada) and confirmed to be free of mycoplasma contamination using the PCR-based Venor GenM Mycoplasma Detection Kit (MilliporeSigma, catalog no. MP0025).

### B-cells isolation and stimulation assays

Peripheral blood mononuclear cells (PBMCs) were isolated from healthy donor-derived buffy coats (Miltenyi Biotec, catalog no. 150-000-572) by density-gradient centrifugation using Ficoll-Paque Plus (MilliporeSigma, catalog no. GE17-1440-02) and LeucoSep tubes (VWR, catalog no. 89048-938), according to the manufacturers’ protocols. PBMCs were washed once in PBS + 2 mM EDTA (Invitrogen, catalog no. 15575020) and centrifuged at 300 × g for 5 min. The red blood cells were lysed using 5 mL RBC lysis buffer (BioLegend, catalog no. 420301) for 15 min at room temperature. PBMCs were washed twice in PBS + 2 mM EDTA, centrifuged at 200 × g for 10 min and resuspended in 1 mL FACS buffer for cell counting. Subsequently, B cells were purified from PBMCs using magnetic-activated cell sorting (MACS, Miltenyi Biotec) using the following kits: CD19^+^ B cells (catalog no. 130-091-151), naïve B cells (catalog no. 130-091-150), and memory B cells (catalog no. 130-093-546). 5 × 10⁵ B cells were seeded in 200 µL of RPMI-1640 + 10% FBS at 37°C in a 96-well plate and stimulated with individual or combined cytokines at the indicated concentrations for 6 – 120 hrs: 20 ng/mL IL-4 (Invitrogen, catalog no. 200-04), 100 ng/mL MEGACD40L (Enzo, catalog no. ALX-522-110), 100 ng/mL LPS (List Labs, catalog no. 201), 10 µg/mL F(ab′)₂ anti-IgG + IgM (Jackson Immunoresearch, catalog no. 109-006-127), 100 ng/mL IFN-γ (Peprotech, catalog no. 300-02).

### B-cells sorting from human thymus

Cell suspension from healthy donor-derived thymus was rapidly defrosted at 37°C, washed with 10 mL of RPMI-1640 with 10% fetal bovine serum (FBS) solution containing DNase I (MilliporeSigma, DN25-10MG) and for 15 min at 37°C. The cell suspension was filtered with MACS SmartStrainers (70 µm) (Miltenyi Biotec, catalog no. 130-098-462) to obtain single-cell suspension. The single-cell suspension was washed once with FACS buffer, centrifuged at 300 × g for 5 min, and resuspended in FACS buffer for cell counting.

Cells were blocked with the Fc block (BD Bioscience, catalog no. 564220) for 15 min on ice in the dark and subsequently stained with the brilliant staining buffer (BD Bioscience, catalog no. 566349) containing fluorochrome-conjugated antibodies against CD45 (BioLegend, catalog no. 304050), CD3 (BioLegend, catalog no. 344822), CD20 (Beckman Coulter, catalog no. IM3629), CD38 (BD Bioscience, catalog no. 560982), CD54 (BD Bioscience, catalog no. 740170), and CD58 (BioLegend, catalog no. 981702) for 30 min on ice in the dark. Cells were washed in FACS buffer, stained with DAPI (Invitrogen, catalog no. D3571) on ice in the dark for 20 min, and subsequently sorted using FACS Aria Fusion Cell Sorter (BD Bioscience, model no. 656700P3). CD20+CD3-CD38-CD54^high^CD58^high^ cells and CD20+CD3-CD38-CD54^low^CD58^low^ cells were collected into 384-well plate (Supplementary Data Fig. 1).

### Live cell isolation for single-cell / single-nucleus library preparation

Live cell isolation and single-cell library preparation were performed. Frozen cell suspensions were rapidly thawed in a 37°C water bath, washed with 10 mL of RPMI-1640 with 10% fetal bovine serum (FBS) solution containing DNase I and for 15 min at 37°C. The cell suspension was filtered with MACS SmartStrainers (70µm) to obtain single-cell suspension. The single-cell suspension was washed once with the FACS buffer, centrifuged at 300 × g for 5 min, and resuspended in the FACS buffer for cell counting. For DLBCL samples, live cells were directly sorted based on DAPI (Sigma, catalog no. D9542) exclusion. To enrich viable B-cell populations in RLN, Thymus, tonsil, and PMBCL, cells were blocked with the Fc block for 15 min on ice in the dark and subsequently stained with the brilliant staining buffer containing fluorochrome-conjugated antibodies against CD3 (Invitrogen, catalog no. 45-0036-42) and CD20 for 30 min on ice in the dark. Cells were washed in FACS buffer, stained with DAPI on ice for 20 min, and subsequently sorted using FACS Aria Fusion Cell Sorter or FACS Aria III Cell Sorter (BD Bioscience, model no. 648282-31-010140-X-X-X, Supplementary Data Fig. 2).

### Library preparation & sequencing (bulk / single cell)

#### RNA-seq of stimulated B cells and cell lines

Stimulated primary B cells and B-cell lymphoma cell lines were washed once with PBS, centrifuged at 300 × g for 5 min, and immediately processed for RNA extraction using the RNeasy Mini Kit (Qiagen, catalog no. 74104) with on-column DNase digestion (Qiagen, catalog no. 79254), according to the manufacturer’s instructions. RNA-seq was performed on biological replicates of stimulated primary B cells.

RNA quality and integrity were assessed, and then ribosomal RNA was depleted from total RNA using the NEBNext® rRNA Depletion Kit v2 (Human/Mouse/Rat) with RNA Sample Purification Beads (New England Biolabs, catalog no. E7405X). rRNA-depleted RNA was then used for stranded library preparation with the NEBNext® Ultra™ II Directional RNA Library Prep Kit for Illumina (New England Biolabs, catalog no. E7760L), following the manufacturer’s protocol. Libraries were indexed using NEBNext® Multiplex Oligos for Illumina® (96 Index Primers) (New England Biolabs, catalog no. E6609L). Final libraries were purified, quantified, pooled at equimolar concentrations, and sequenced on an AVITI system (Element Biosciences) using a High Output paired-end kit.

#### ATAC-seq of stimulated B cells

Stimulated primary B cells were washed once in PBS and centrifuged at 300× g for 5 min and immediately proceeded to ATAC-seq library preparation using the ATAC Library Prep Kit (Active Motif, catalog no. 53150) according to the manufacturer’s instructions. Briefly, 200,000 cells per sample were pelleted and washed with cold PBS. The cells were lysed to release nuclei, which were immediately subjected to transposition reactions. The transposition reaction was performed using the enzyme and buffer provided in the kit to tag and fragment accessible chromatin regions. After incubation at 37°C for 60 min, transposed DNA was purified using the purification columns included in the kit. The purified DNA was amplified using PCR to generate sequencing-ready libraries. The final libraries were purified, and their quality and size distribution were assessed using a Bioanalyzer 2100. Libraries were quantified using a Qubit fluorometer (Thermo Fisher Scientific) before being sequenced on an AVITI using a High Output paired end kit.

#### RNA-seq of sorted thymic B cells

For transcriptomic profiling, approximately 2,500∼4,000 FACS-sorted cells were collected for each thymic B-cell fraction and processed using the SMART-Seq® v4 Ultra Low Input RNA Kit for Sequencing (Takara Bio, catalog no. 634772) according to the manufacturer’s instructions. Briefly, sorted cells were lysed, and polyadenylated RNA was reverse-transcribed using a template-switching mechanism to generate full-length cDNA, followed by PCR amplification. Amplified cDNA was purified, quantified, and assessed for size distribution prior to library preparation.

Sequencing libraries were generated from amplified cDNA using the Nextera XT DNA Library Preparation Kit (Illumina, catalog no. FC-131-1024) following the manufacturer’s protocol. Libraries were pooled at equimolar ratios and sequenced on an AVITI system (Element Biosciences) using a High Output paired end kit.

#### Deparaffinization and nuclear extraction of FFPE samples

FFPE tissue blocks were serially sectioned at 25 µm thickness, with four sections collected per sample (total tissue thickness, 100 µm). Deparaffinization was performed according to the manufacturer’s protocol (10x Genomics, CG000553 Rev B). Briefly, tissue sections were treated twice with xylene for 10 min each, followed by sequential rehydration in 100%, 95%, and 70% ethanol and a final 5-min incubation in distilled water. Deparaffinized tissue was then enzymatically digested and mechanically dissociated using a gentleMACS Octo Dissociator with Heaters (Miltenyi Biotec).

#### 10x Chromium snRNA-seq and library preparation

Single-cell nuclei extracted from FFPE were converted into barcoded snRNA-seq libraries using Chromium Fixed RNA Profiling Reagent kits (10x Genomics, catalog no. PN-1000474), according to the manufacturer’s instructions (CG000477 Rev B), aiming for ∼5,000 nuclei per sample. All libraries were subjected to quality control and quantified using the High Sensitivity DNA Kit using the 2100 Bioanalyzer (Agilent Technologies). Libraries were sequenced on the Nextseq 550 platform (Illumina) using a High Output paired end kit.

#### 10x Chromium sn/scRNA-seq

For single-nucleus chromatin accessibility and gene expression profiling, nuclei were isolated from 250,000 sorted cells using the Low Cell Input Nuclei Isolation protocol for Single Cell Multiome ATAC + Gene Expression sequencing (10x Genomics, CG000365 Rev C). Transposition, GEM generation and barcoding, library pre-amplification, and ATAC and gene expression library construction were performed according to the Chromium Next GEM Single Cell Multiome ATAC + Gene Expression User Guide (CG000338 Rev G), using the Chromium Next GEM Single Cell Multiome ATAC + Gene Expression Reagent Bundle (PN 1000283) in combination with the Chromium Next GEM Chip J Single Cell Kit (PN 1000234), targeting a recovery of 5,000 nuclei. Gene expression and ATAC libraries were indexed using the Dual Index Kit TT Set A (PN 1000215) and the Single Index Kit N Set A (PN 1000212), respectively.

For single-cell gene expression and B-cell receptor (BCR) profiling, 5,000 cells per sample were applied for library preparation using the Chromium Next GEM Single Cell 5′ Kit v2 (PN 1000265) in combination with the Chromium Next GEM Chip K Single Cell Kit (PN 1000287), following the manufacturer’s protocol (CG000331 Rev F). Paired BCR libraries were generated from the same amplified single-cell cDNA using the Chromium Single Cell Human BCR Amplification Kit (PN 1000253). Gene expression and BCR libraries were dual indexed using the Dual Index Kit TT Set A (PN 1000215). Library quality was assessed using the High Sensitivity DNA Kit using the 2100 Bioanalyzer (Agilent Technologies), and libraries were sequenced on an Illumina NextSeq 550 platform using a High Output paired end kit.

#### Primary data preprocessing of single-cell data modalities

All raw sequencing data were demultiplexed using *cellranger mkfastq* (v7.0.1; 10x Genomics), except for 10x Chromium 3′ snRNA-seq + snATAC-seq (multiome) libraries, which were demultiplexed and converted to FASTQ format using *cellranger-arc mkfastq* (v2.0.2; 10x Genomics). The resulting FASTQ files were processed using modality-specific pipelines from the Cell Ranger suite, as described below.

#### Data preprocessing of 10x Chromium Fixed snRNA-seq

FASTQ files were processed using *cellranger multi* (v7.0.1) with the Chromium Fixed RNA Profiling configuration. Reads were aligned to the human reference genome (hg38) using the Chromium Human Transcriptome Probe Set v1.0.1, followed by probe assignment and generation of gene–barcode count matrices. Gene-level UMI counts were generated by aggregating confidently assigned probe-ligation events.

#### Data preprocessing of 10x 5’ Chromium scRNA-seq

FASTQ files were processed using *cellranger count* (v7.0.1) for read alignment and quantification against the human reference genome (GRCh38-2020-A), generating gene– barcode count matrices.

#### Data preprocessing of 10x 5’ Chromium scBCR-seq

FASTQ files were processed using cellranger vdj (v7.0.1) to generate assembled BCR contigs for downstream immunoglobulin repertoire analysis. Assembled BCR contigs were imported into the Immcantation framework and converted to AIRR-compliant format. Sequence alignment and V(D)J gene assignment were performed using IgBlast^43^ against the IMGT reference database using the Change-O package (v1.3.1)^44^. Non-productive rearrangements, including sequences containing stop codons, frameshifts, or ambiguous base calls, were excluded. Cells with multiple heavy-chain sequences or no detectable productive IGH sequence were excluded from BCR-based analyses. BCR-informative cells were defined as cells with a single productive IGH sequence.

#### Data preprocessing of 10x 3’ Chromium snRNA-seq + snATAC-seq

FASTQ files were processed using *cellranger-arc count* (v2.0.2) for joint alignment and annotation of gene expression and chromatin accessibility data, generating gene expression count matrices and chromatin accessibility fragment files.

#### Overview of single cell/single nucleus transcriptome analysis

This study comprised multiple single-cell/single-nucleus transcriptomic data regimes: FFPE-derived 10x Chromium Fixed snRNA-seq, fresh-derived 10x Chromium 5′ scRNA-seq and 3′ snRNA-seq. Due to differences in library preparation chemistry and RNA integrity, FFPE-derived snRNA-seq data were analyzed independently, whereas fresh-derived datasets were jointly integrated for downstream analyses. Modality-specific preprocessing and analysis of pipelines were applied accordingly, as described below.

### Downstream analysis of FFPE-derived 10x Chromium Fixed snRNA-seq

#### Quality control

Low-quality cells were excluded prior to downstream analyses. Specifically, cells were removed if ≥20% of sequencing reads aligned to mitochondrial genes, or if they represented extreme outliers based on global quality metrics, defined as cells exceeding 3 median absolute deviations from the median number of detected genes (Supplementary Data Fig. 3; <u>Supplementary Note 1</u>).

#### Ambient RNA removal

Ambient RNA contamination was estimated and corrected separately for each sample using SoupX (v1.6.0)^45^. Raw and filtered count matrices generated by Cell Ranger were used as input, and contamination fractions were estimated using the *autoEstCont* function with cluster information provided from *preliminary* clustering. Ambient RNA–corrected count matrices generated using *adjustCounts* were used for subsequent normalization, clustering, and differential-expression analyses.

#### Doublet detection

Putative doublets were initially identified using DoubletFinder (v2.0.6)^46^ with default parameters. However, due to the probe-based nature of Fixed RNA Profiling and the increased prevalence of ambient RNA contamination in FFPE-derived datasets, algorithmic doublet detection alone was insufficient to reliably distinguish true biological doublets from transcriptionally contaminated singlets. Therefore, additional lineage-aware quality control was performed based on canonical marker gene expression. Cells were first provisionally assigned to major lineage categories (B cells, T cells, myeloid cells, epithelial cells, and stromal cells) based on preliminary unsupervised clustering and established marker gene expression patterns. Within each lineage, cells were screened for inappropriate expression of markers characteristic of other lineages. B cells expressing T-cell markers (*CD3D* or *CD3E*), stromal markers (*COL1A1* or *LUM*), or myeloid markers (*CD68*) were excluded. Similarly, T cells expressing B-cell markers (*CD19* or *MS4A1*), stromal markers (*COL1A1* or *LUM*), or myeloid markers (*CD68*) were removed. Myeloid cells expressing T-cell markers (*CD3D* or *CD3E*) or B-cell markers (*CD19* or *MS4A1*), and stromal cells expressing lymphoid markers (*CD19, MS4A1, CD3D*, or *CD3E)* were also filtered out. Any detectable expression of lineage-inappropriate markers above background levels was considered evidence of contamination.

## Data processing

The SoupX-corrected expression matrices were loaded and processed using the Seurat^47^ framework. First, they were log-normalized using *NormalizeData* and scaled using *ScaleData*. Highly variable genes were identified using *FindVariableFeatures*, and the top 2,000 variable genes were used for downstream analyses. Principal component analysis (PCA) was performed using *RunPCA*, and the first 30 principal components were used for the following batch effect evaluation and correction.

### Batch effect evaluation and correction

Quantitative assessment of batch effects was performed using established batch-mixing metrics, including the local inverse Simpson’s index (LISI), average silhouette width (ASW), and adjusted rand index (ARI), as previously described^48^. These metrics evaluate batch mixing at local and global scales, respectively, while accounting for preservation of biological structure (Supplementary Data Fig. 4, Supplementary Note 2). Batch effects arising from differences between batches were corrected using the *RunHarmony* function from Harmony (v1.2.3)^49^. The first 30 Harmony components were used to construct a shared nearest-neighbor graph with the *FindNeighbors* function, followed by unsupervised clustering with the *FindClusters* function at a resolution of 0.8. UMAP embeddings were generated from the same 30 Harmony components using the *RunUMAP* function.

### Cell type annotation

Clusters generated from unsupervised clustering were manually assigned to a cell type by comparing the mean expression of known markers across cells in a cluster. Markers used to annotate cells included *CD19, CD20* (B cells), *CD3, CD4, CD8* (T cells), *CD14, CD68* (myeloid cells), *COL1A1, LUM, PECAM1* (stromal cells).

Malignant B cells were identified using an integrated approach combining clustering behavior, canonical marker expression, and inferred copy number profiles. Normal B cells from RLN and tonsil samples were used as a reference population. B cells from PMBCL samples that were co-clustered with normal control derived B cells and lacked malignant-associated transcriptional features were annotated as non-malignant. In contrast, B cells from PMBCL samples that formed transcriptionally distinct clusters separate from RLN and tonsil B cells and exhibited expression of established PMBCL-associated markers, including *CCL17*, *CCL22*, *CD274*, and *PDCD1LG2*, were annotated as malignant.

### Copy number inference

To further validate malignant and non-malignant B-cell populations, inferred copy number variation (CNV) profiles were generated using the inferCNV pipeline^50^. For each sample, PMBCL-derived B cells were compared against a reference panel of transcriptionally normal B cells, which included B cells from all RLN and tonsil samples.

Gene expression values were ordered according to chromosomal position (hg38) and smoothed across adjacent genomic regions to infer large-scale copy number gains and losses. CNV profiles were used as complementary evidence of malignant identity rather than as a sole classification criterion.

## Differential analysis

Differential expression (DE) analyses were performed using a pseudobulk strategy. For each sample, raw counts from cells belonging to the same subcluster were aggregated to generate sample-level count matrices. Differential expression testing was then conducted using DESeq2 (v1.38.3)^51^, which models count data using a negative binomial generalized linear model framework. Size-factor normalization was applied to account for differences in sequencing depth across samples. Statistical significance was determined using the Wald test, and *p* values were adjusted for multiple testing using the Benjamini–Hochberg method. Differentially expressed genes were defined as those exhibiting an average expression level at least 2.0-fold higher in a given subcluster compared with other subclusters of the same cell type and an adjusted *p* value < 0.05.

### Correlation analysis

To assess transcriptional similarity between cell populations across tissues and pathological conditions, correlation-based analyses were performed using average gene expression profiles. For each annotated cell cluster, average expression values were calculated across all genes retained after quality control. Pairwise correlations were then computed to quantify the degree of similarity between B-cell subsets derived from normal thymus, reactive lymph nodes, tonsils, and lymphoma samples.

### Integrated analysis of 10x Chromium 5’ scRNA-seq and 10x 3’snRNA-seq

Unless otherwise specified, quality control (Supplementary Data Fig. 5-6, Supplementary Note 3-4), doublet removal, normalization, dimensionality reduction, clustering, copy number inference, correlation analysis, differential expression analyses and batch effect evaluation (Supplementary Data Fig. 7, Supplementary Note 5) was performed using the same procedures described above.

### Data integration

After initial preprocessing, unsupervised clustering was performed separately for each dataset, followed by manual annotation of cell clusters based on canonical lineage marker expression. In B-cell enriched samples, residual non-B-cell populations were identified and excluded. In B-cell unenriched DLBCL samples, only clusters exhibiting robust expression of B cell-specific markers and absence of non-B-cell lineage markers were retained. All downstream analyses were conducted exclusively on the curated B-cell populations from each tissue and disease context.

Given the high abundance of immunoglobulin (Ig) transcripts in B cells, we excluded Ig genes prior to integration analyses to prevent their dominance from obscuring biologically meaningful variation in non-Ig gene expression, following a previously validated integration strategy on a B-cell atlas study^31^. In addition, mitochondrial genes, ribosomal genes, mitochondrial ribosomal genes, and heat-shock protein genes were also excluded to reduce technical noise. The processing procedures were the same as described above. Batch effects arising from differences between data modalities and individual samples were corrected using Harmony (v1.2.3)^49^, with batch correction applied across both data modalities and sample variables using the *RunHarmony* function. The top 30 Harmony-corrected components were retained for downstream analyses.

### Cell type annotation

Cell type annotation was performed using a label transfer approach implemented in Seurat, followed with manual correction. Briefly, a B-cell single cell atlas dataset with established cell type annotations^31^ was used to identify shared transcriptional features with query datasets using the *FindTransferAnchors* function. These anchors were then used to transfer cell type labels from the reference to the query data using the *TransferData* function.

Transferred labels were assigned to individual cells based on prediction scores and subsequently evaluated by manual inspection of canonical marker gene expression to ensure consistency with known biological cell identities. Where necessary, annotations were refined based on marker expression patterns.

### Gene Ontology enrichment analysis

Gene Ontology (GO) enrichment analysis was then performed using the *enrichGO* function implemented in the clusterProfiler (v.4.18.4) package^52^, focusing on the Biological Process (BP) ontology. Gene symbols were mapped using the org.Hs.eg.db annotation database (v3.16.0). Multiple testing correction was applied using the Benjamini–Hochberg method.

### Gene set enrichment analysis

To assess whether the gene signatures of CCL17^+^ thymic B cells were enriched in the sorted double-positive (DP) versus double-negative (DN) comparison, pre-ranked gene set enrichment analysis was performed using the R package fgsea (v.1.29.1) (https://bioconductor.org/packages/release/bioc/html/fgsea.html). Genes were ranked according to average log2 fold change from the DP vs DN differential expression analysis.

### Gene set variation analysis

Gene set variation analysis (GSVA) was performed to assess pathway activity at the transcriptional level across CCL17^+^ thymic B cell and malignant B-cell populations. GSVA was conducted using averaged gene expression matrices derived from normalized expression data. Enrichment scores were computed using the *gsva* function in the GSVA package (v2.3.1)^53^.

### Trajectory analysis

Trajectory inference was performed to reconstruct developmental relationships among normal B-cell populations from thymus, RLN and tonsil samples using the Slingshot package (v2.18)^54^. Dimensionality reduction embeddings derived from harmony-corrected matrix were used as input for trajectory analysis, together with refined B-cell cluster annotations. To characterize molecular dynamics along the inferred trajectories, gene expression, chromatin accessibility, and transcription factor activity were examined as functions of pseudotime. Trends were visualized separately for each tissue using generalized additive models (GAMs), with a spline basis dimension of 10 (k = 10).

### Ingenuity Pathway Analysis (IPA)

IPA (v122103623; QIAGEN Inc) was performed to identify canonical pathways, potential upstream regulators, and gene networks. Default settings were used for core analysis, and significant pathways and regulators were selected using Fisher’s exact test (*p* < 0.05) and z-score thresholds, as implemented in IPA.

### CytoSig

To complement Ingenuity Pathway Analysis and independently infer upstream signaling activities, cytokine signaling inference was performed using CytoSig^33^. CytoSig was applied to pseudo-bulk gene expression profiles derived from normal B-cell populations. Raw gene expression count matrices were aggregated at the population level to generate pseudo-bulk profiles and normalized transcripts per million (TPM). TPM values were then scaled (TPM/10), log₂-transformed [log₂(TPM/10 + 1)] and input into CytoSig analysis. CytoSig inferred relative cytokine activity scores for each curated cytokine, reflecting upstream signaling programs associated with transcriptional states of normal B-cell subsets.

### B-cell receptor repertoire and somatic hypermutation analyses

#### Clonal Type Construction

BCR clonotypes were constructed separately for each patient. Clones were defined based on shared V and J gene usage and junctional sequence similarity of the immunoglobulin heavy-chain (IGH) CDR3 region. Pairwise distances were calculated using a nucleotide Hamming-distance metric using the *distToNearest* function, and clonal thresholds were determined from the distance-to-nearest-neighbor (DTNN) distribution using the *findThreshold* function in the SHazaM package (v1.3.0)^44^, enabling objective delineation of clonal relationships^55–57^. Expanded clonotypes were defined as clonotypes represented by more than 20 BCR-informative cells within an individual patient.

#### Somatic hypermutation frequency calculation

IGH sequences within each clone were aligned to their inferred germline sequences using the *createGermlines* function implemented in the dowser package (v2.4.0)^58^, with reference germline alleles obtained from the IMGT database. Somatic hypermutation (SHM) frequencies were calculated as the percentage of nucleotide divergence from the inferred germline sequence for each productive heavy chain using the *observedMutations* function in the SHazaM package (v1.3.0)^44^. Only productive rearrangements with unambiguous V(D)J assignments were included in SHM analyses.

#### Phylogenetic analysis with BCR sequences

Lineage trees were reconstructed from clonally related IGH sequences using maximum-likelihood phylogenetic inference implemented in IgPhyML (v1.1.5)^59^, with inferred germline sequences used as the root. Tree topologies were inferred from aligned IGH sequences, and branch lengths reflected the accumulation of SHM. GEX-defined PMBCL transcriptional states were linked to BCR sequences using matched single-cell barcodes and mapped onto the reconstructed lineage trees. Early versus late clonal positions were defined according to SHM-derived evolutionary distance from the inferred germline root, with lower SHM corresponding to earlier clonal positions.

#### BCR Phylogenetic Node Depth Analysis

We define the depth d_n_of a node n as its hop count from the germline root along the directed tree:

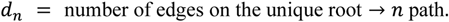

The root itself has depth 0; its immediate children have depth 1, and so on. Hop count is a topology-only measure that preserves the *earlier vs terminal* ordering of nodes without requiring branch-length estimates (which are only partially available from Figure 2H).

For clone *c* and cell type *t*, we summarise the spatial distribution of that cell type across the tree as the cell-count-weighted mean depth:

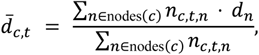

Where n_c,t,n_ is the number of cells of type t at node n in clone *c*. A larger indicates that the cell type is preferentially localised at deeper (more terminal) nodes.

We evaluated the ordered hypothesis

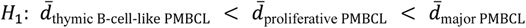

using four complementary approaches.

(a) Within-clone permutation test (primary). The observed effect size is

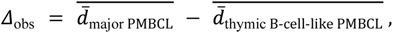

the difference of clone-level means across the three clones. The permutation null is generated by shuffling cell-type labels *within each clone* (preserving the paired structure and the per-clone cell-type counts), recomputing *Δ* on 5,000 permutations, and taking

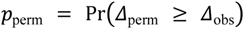

as a one-sided test in the direction of the hypothesis.

(b) Linear mixed model with clone as a random intercept (primary).

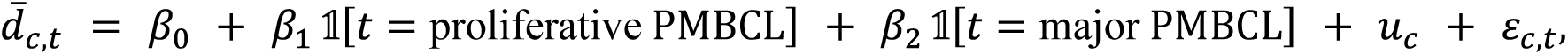 with 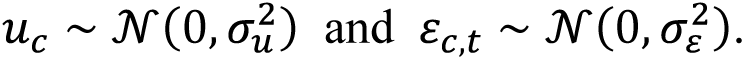 Clone-level random intercepts absorb between-clone variation in baseline depth. Pairwise contrasts use Kenward–Roger degrees of freedom with a Tukey adjustment for multiplicity.

(c) Ordered polynomial contrasts on the cell-type fixed effect:

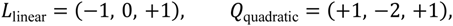

which respectively test for a monotonic trend and for curvature around the intermediate (proliferative PMBCL) cell type.

(d) Jonckheere–Terpstra trend test (secondary). A rank-based test for an ordered alternative across the three cell-type groups. It is reported as a supporting analysis but is treated as secondary because it ignores the within-clone pairing and is therefore anticonservative with only three clones.

### Chromatin accessibility and regulatory landscape analysis

#### Quality control

Downstream chromatin accessibility analyses were performed using the Signac framework (v1.13.0)^60^. Cells were retained if they exhibited a nucleosome banding signal < 4, a transcription start site (TSS) enrichment score > 2, and between 1,000 and 100,000 fragments mapped within peaks (Supplementary Data Fig. 8, Supplementary Note 6). Cells failing to meet any of these criteria were excluded from downstream analyses.

#### Gene activity score

To estimate gene-level regulatory activity from chromatin accessibility data, gene activity scores were computed using the *GeneActivity* function in the Signac^60^ package.

Gene activity matrices were subsequently normalized using standard Seurat normalization workflows and were used for downstream analyses.

#### Correlation analysis

To assess chromatin status similarity between cell populations across tissues and pathological conditions, correlation-based analyses were performed using average gene activity scores. For each annotated cell cluster, average gene activity scores were calculated across all genes retained after quality control. Pairwise correlations were then computed to quantify the degree of similarity between B-cell subsets derived from normal thymus, reactive lymph nodes, tonsils, and lymphoma samples.

#### Transcription factor motif activity analysis

Transcription factor (TF) motif activity was inferred from single-cell chromatin accessibility data using the chromVAR package (v1.32.0)^60,61^. Human transcription factor binding motifs were obtained from the JASPAR2020 database (v0.99.10)^62^, restricted to non-redundant motifs for *Homo sapiens*. Position frequency matrices were retrieved using the TFBSTools package (v1.48.0)^63^.

Motif annotations were added to the chromatin accessibility peak set using the *AddMotifs* function implemented in Signac^60^, with the BSgenome.Hsapiens. UCSC.hg38 reference genome (v1.4.5) used to determine genomic sequence content. chromVAR deviation scores were subsequently computed using the *RunChromVAR* function, which quantifies bias-corrected variability in chromatin accessibility associated with each TF motif while accounting for technical confounders.

The resulting chromVAR deviation matrix, representing relative TF motif activity at single-cell resolution, was used for downstream differential TF motif activity analysis. Differential activity was quantified based on differences in average chromVAR deviation scores between cell populations. TF motifs were defined as differentially active if they met both an adjusted *p* value < 0.05 and an average deviation score difference > 1.

### Projection of single-cell–derived programs onto bulk transcriptomic and epigenomic data

#### Bulk RNA-seq

Raw sequencing reads were aligned to the human reference genome (hg38) using STAR (v2.7.11)^64^. Only uniquely mapped reads were retained. Gene-level read counts were quantified using featureCounts (v2.0.6)^65^ based on gene annotations corresponding to the hg38 reference genome. The resulting gene count matrices were used for downstream normalization and differential expression analyses using DESeq2 (v1.38.3)^51^.

To evaluate the activity of single-cell-derived transcriptional programs in bulk RNA-seq samples, CCL17⁺ thymic B-cell and GC-like B-cell signatures were derived by comparing each respective population with all other normal B-cell populations. For each signature, genes were selected based on significant upregulation in the corresponding single-cell population (adjusted *p* value < 0.05 and average log2 fold change > 0.5). To account for both the magnitude and robustness of differential expression, each gene was assigned a weight calculated as the product of its average *log2* fold change, and the negative logarithm of the adjusted *p*-value. Gene weights were subsequently normalized such that the sum of weights for each signature equaled one. Only genes present in both the single-cell–derived signature and the bulk RNA-seq dataset were retained for scoring. For each bulk RNA-seq sample, the signature score was calculated as the sum of gene expression values multiplied by their corresponding normalized weights:

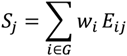

Where s_j_ represents the signature score for sample j, w represents the normalized weight of gene i,E_ij_ represents the expression value of gene i in sample j, and G denotes the set of signature genes shared with the bulk RNA-seq dataset.

#### Bulk ATAC-seq

Raw sequencing reads were aligned to the human reference genome (hg38) using BWA-MEM2 (v2.2.1)^66–68^. Reference genome indexing was performed once prior to alignment.

Alignment was carried out in paired-end mode using multi-threaded execution. Aligned reads were converted to BAM format, coordinate-sorted, and indexed using samtools (v1.2.0)^67,68^. Low-quality reads (mapping quality < 30) and reads mapping to the mitochondrial genome were removed. PCR duplicates were identified and removed followed by mate fixing and duplicate marking, with duplicate reads excluded from downstream analyses. Accessible chromatin regions were identified by peak calling using MACS2 (v2.2.9.1)^69^. Peaks were called independently for each sample using paired-end mode without model building, with a fixed shift of zero and a false discovery rate threshold of *q* < 0.01. The resulting peak sets were used for downstream chromatin accessibility analyses.

Gene activity scores were calculated by assigning chromatin accessibility peaks to genes based on proximity to transcription start sites (±2 kb). For each gene, accessibility signals from all assigned peaks were aggregated to generate a gene-level activity score, reflecting the cumulative chromatin accessibility associated with the gene locus.

To assess the activity of single-cell–defined chromatin accessibility programs in bulk ATAC-seq samples, weighted gene activity scores were calculated using gene activity signatures derived from differential accessibility analysis of snATAC-seq data. Genes were retained if they were significantly upregulated in the corresponding single-cell population (adjusted *p* < 0.05 and average log₂ fold change > 1).

Gene weights were calculated analogously to the bulk RNA-seq analysis, based on the product of average log₂ fold change, and the negative logarithm of the adjusted *p* value. Weights were normalized such that the sum of weights within each signature equaled one. Only genes shared between the single-cell–derived signatures and bulk ATAC-seq gene activity matrices were retained.

For each bulk ATAC-seq sample, a weighted signature score was calculated as the sum of gene activity values multiplied by their corresponding normalized weights. These scores represent the relative activity of single-cell–defined chromatin accessibility programs in bulk chromatin accessibility data and were used for downstream comparative analyses.

### Spatial transcriptomic analysis of thymus and PMBCL TMAs

#### TMA construction

FFPE tissue microarrays (TMAs) were sectioned at 5 μm thickness using a microtome and mounted onto VWR Superfrost Plus Micro Slides (cat. no. 48311-703) to ensure optimal adherence. Slides were dried overnight at 37 °C. All TMAs were processed using the same standardized CosMx Spatial Molecular Imager workflow. Briefly, slides underwent deparaffinization, heat-induced target retrieval using a standardized retrieval time, followed by proteinase-mediated permeabilization. Procedures were performed according to the manufacturer’s protocol for the CosMx platform (https://nanostring.com/products/cosmx-spatial-molecular-imager/single-cell-imaging-overview). RNA in situ hybridization was performed using the Human 6K Discovery Panel (6K-plex, RNA) together with a 200-gene Custom Add-On panel (Table S14). Hybridization was carried out overnight at 37 °C, followed by stringent washing. A flow cell was assembled on the slide, and cyclic RNA readout was performed on the CosMx instrument. Prior to RNA detection cycling, morphology and segmentation markers were applied. Nuclear staining was performed using DAPI. Segmentation markers included human CD298/B2M and a combined human PanCK/CD45 reagent, with human CD68 used as an additional a la carte marker to assist immune cell identification. Thymus TMAs consisted of 1.0 mm cores, whereas PMBCL TMAs consisted of 0.6mm cores. Fields of view (FOVs), each covering an area of approximately 0.5 × 0.5 mm², were selected by expert pathologists based on histopathological evaluation of the tissue sections. Up to four FOVs were collected per core, depending on tissue coverage and run constraints. Immunofluorescence (IF) images and transcript localization were obtained from the AtoMx platform (NanoString Technologies).

#### Segmentation

Cell segmentation of CosMx spatial transcriptomics images was performed using the CellposeSAM framework^70^. IF images were processed from multi-channel TIFF files, in which membrane and nuclear signals were used to guide cell boundary detection. Specifically, membrane marker channels (a pan-membrane marker, PanCK, CD45 and CD68) were merged into a single composite membrane signal, and the nuclear (DNA) channel was used to aid cell center identification during segmentation. Following segmentation, individual transcript coordinates were spatially mapped to the revised CellposeSAM segmentation masks. Transcripts located within a segmented cell were assigned to the corresponding cell identifier, and a new cell-by-gene count matrix was generated by aggregating transcript counts for each cell. The resulting expression matrix and segmentation-derived cell metadata were used for downstream quality control, cell-type annotation and spatial analyses.

#### Annotation of thymic architecture in CosMx spatial data

Cells were annotated as cortical, medullary, perivascular space (PVS), vessel, or Hassall’s corpuscle regions following the TissueTag approach^23^. Expert pathologists first annotated regions in 11 out of 60 cores. A random forest pixel classifier (50 estimators with maximum depth 10) was trained on the 11 annotated cores and predicted the regions in the unannotated cores. The random forest pixel classifier input was augmented with intensity, edge, and local structure features using *skimage.feature.multiscale_basic_features*. Cells were assigned to a region based on the classification of the cells’ centroid pixels. Cells whose assigned region had a probability of less than 0.66 were excluded.

#### Quality control

QC was performed at both the single-cell and FOV levels. For each patient, cell-level RNA counts were evaluated to identify outlier cells. Specifically, the MAD of total RNA counts was calculated separately for each patient, and cells with RNA counts exceeding 3 MAD from the patient-specific median were classified as low quality. In addition, cells with fewer than 50 detected RNA counts were excluded from downstream analyses (Supplementary Data Fig. 9, Supplementary Note 7).

At the FOV level, segmentation quality was manually assessed by expert inspection through overlaying segmented cell masks with corresponding IF images. FOVs exhibiting poor IF staining quality or suboptimal cell segmentation performance were excluded. In parallel, the proportion of low-quality cells was calculated for each FOV, and FOVs in which more than 50% of cells were classified as low quality were removed.

#### Cell type annotation

Cell type annotation was performed using a hierarchical, reference-based label transfer strategy implemented in Seurat. First, major cell types were identified using annotations derived from FFPE snRNA-seq reference data. Cells with a maximum prediction probability below 70% were considered low-confidence assignments and were assigned as uncertain. Cells within each major lineage were then subsetted and subjected to a second round of reference-based label transfer to assign more fine-grained cell type identities.

#### Identification of PMBCL cell states

Following the identification of major cell lineages, B cells were subsetted for further identification of malignant and non-malignant B cells. Using the annotated fresh scRNA-seq and snRNA-seq dataset as a reference, malignant and non-malignant B cells were first distinguished in the 13-case FFPE snRNA-seq cohort through Seurat-based label transfer. Within the malignant B-cell compartment, three PMBCL cell states initially defined in the fresh scRNA-seq and snRNA-seq dataset were subsequently assigned to FFPE snRNA-seq cells using the same reference-based label-transfer approach. To extend these annotations to the CosMx spatial transcriptomics dataset, FFPE snRNA-seq samples with matched CosMx data were first used as references to transfer malignant cell-state annotations to their corresponding CosMx samples. The resulting annotated CosMx cells were then used as a reference to project these annotations to the remaining CosMx samples, enabling consistent identification of the three malignant PMBCL states across the spatial cohort.

#### Spatial score

Spatial relationships between cells were quantified to characterize local cellular organization and define tumor microenvironment ecosystems. For each segmented cell, Euclidean distances were calculated to the five nearest neighboring cells of each phenotypic cell type. Distances were capped at 50 μm to restrict the analysis to biologically relevant local neighborhoods and to limit the influence of long-range spatial effects^34^.

Distance values were subsequently scaled to a 0–1 range and inverted such that higher values reflected greater local enrichment of a given cell type in the immediate vicinity of each cell. These transformed distance metrics therefore represent proximity-based enrichment scores that describe the local cellular context surrounding individual cells.

## Statistical analyses

Statistical analyses were performed using R (v4.4.1) and GraphPad Prism (v10.4.0). Statistical tests were selected according to the experimental design and are specified in the corresponding figure legends. Multiple-comparison correction was applied where indicated. All statistical tests were two-sided unless otherwise specified, and *p* < 0.05 was considered statistically significant.

