## Supplementary Figures for "Developmental origins of thymic B-cell lymphomas"

* Authors contributed equally to this work.

This PDF file includes:

**Supplementary Notes:**

**Figure S1. Age distribution and multi-modal profiling of analyzed cases.**

**Figure S2. Global cell-type annotation of thymic FFPE single-nucleus RNA-seq data.**

**Figure S3. Validation of thymic B-cell states in an independent external dataset.**

**Figure S4. Integrated analysis of single-cell B-cell atlas across tissues and lymphomas.**

**Figure S5. Lineage annotation and malignant cell identification in FFPE snRNA-seq samples.**

**Figure S6. Identification of malignant B-cell state within PMBCL malignant cells.**

**Figure S7. Transcriptional relationships, SHM patterns, and inferred differentiation trajectories across B-cell states.**

**Figure S8. Cytokine- and BCR-dependent regulation of CCL17$^{+}$ and GC-like B-cell programs.**

**Figure S9. Spatial compartmentalization and proximity-associated gene expression in the thymus.**

**Figure S10. Spatial compartmentalization of PMBCL states within a remodeled thymic microenvironment.**

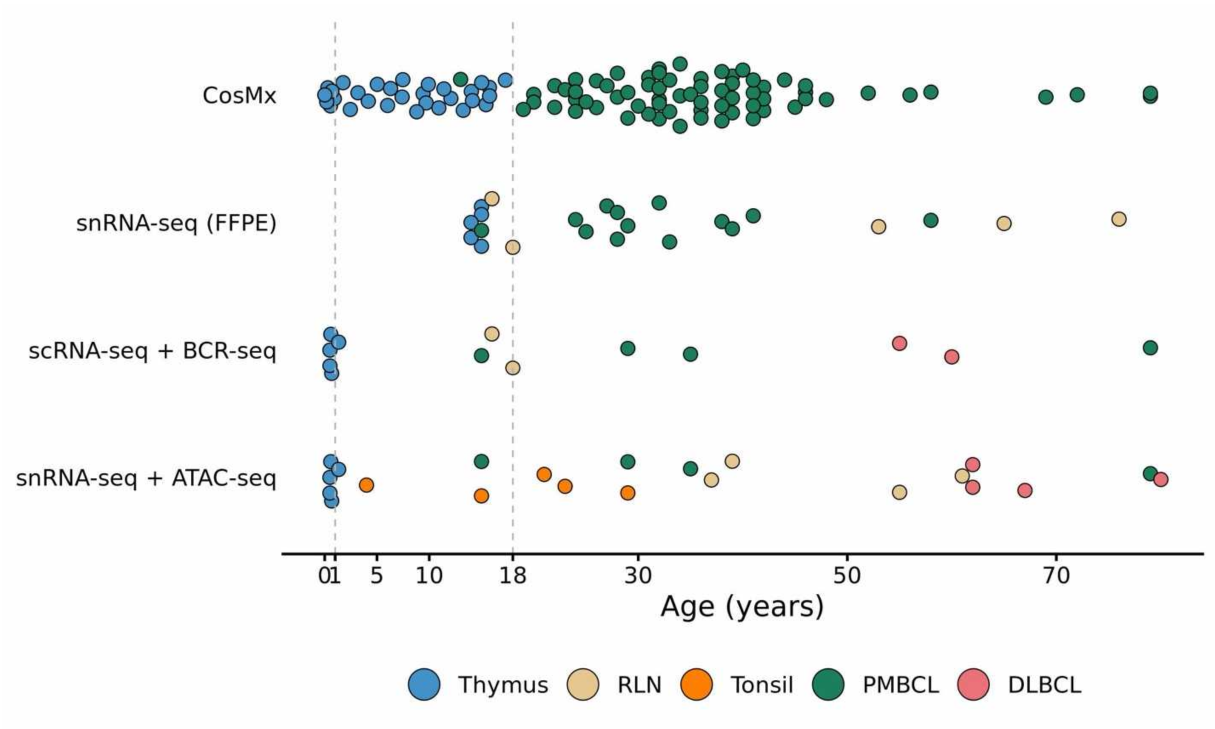


**Figure S1. Age distribution and multi-modal profiling of analyzed cases.**
Scatter plot showing the age of donors or patients (x-axis) across different experimental platforms (y-axis), including CosMx spatial transcriptomics, snRNA-seq (FFPE), scRNA-seq with BCR-seq, and snRNA-seq with ATAC-seq. Each dot represents one sample. Samples are color-coded by tissue or disease of origin. Dashed vertical lines indicate key age thresholds (e.g., pediatric and adult cutoffs). Age information was unavailable for four tonsil samples that were sequenced with FFPE snRNA-seq.

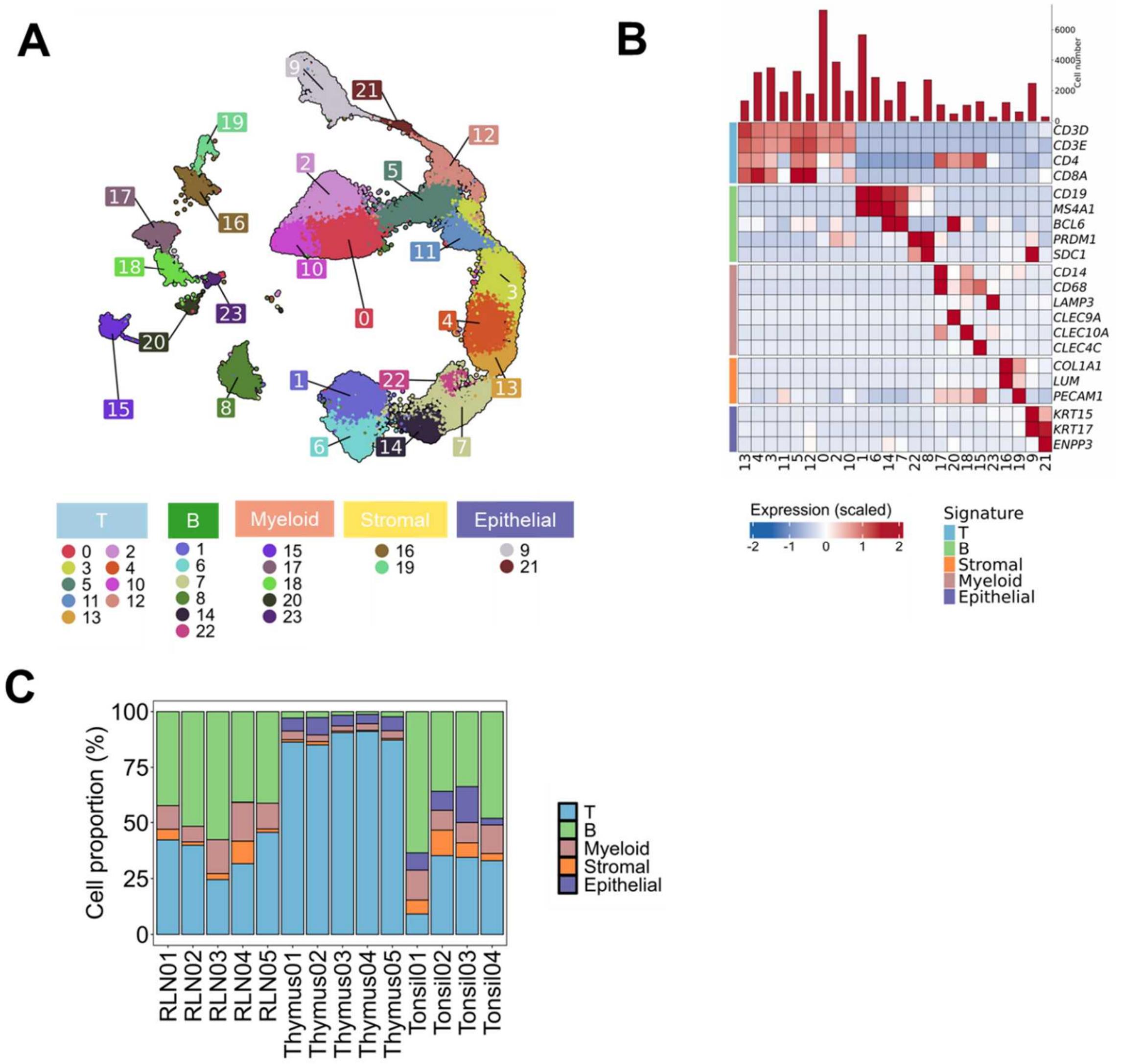


**Figure S2. Global cell-type annotation of thymic FFPE single-nucleus RNA-seq data.**

(A) UMAP visualization of FFPE thymus, RLN and tonsil snRNA-seq data. Each point represents a single cell, colored by cluster identity.

(B) Heatmap showing the canonical marker expression in each cluster identified by snRNA-seq of thymus, RLN and tonsil samples. Expression values are scaled across clusters (z-score), with red indicating higher and blue indicating lower relative expression.

(C) Proportions of major cell populations in the individual thymus, RLN and tonsil samples.

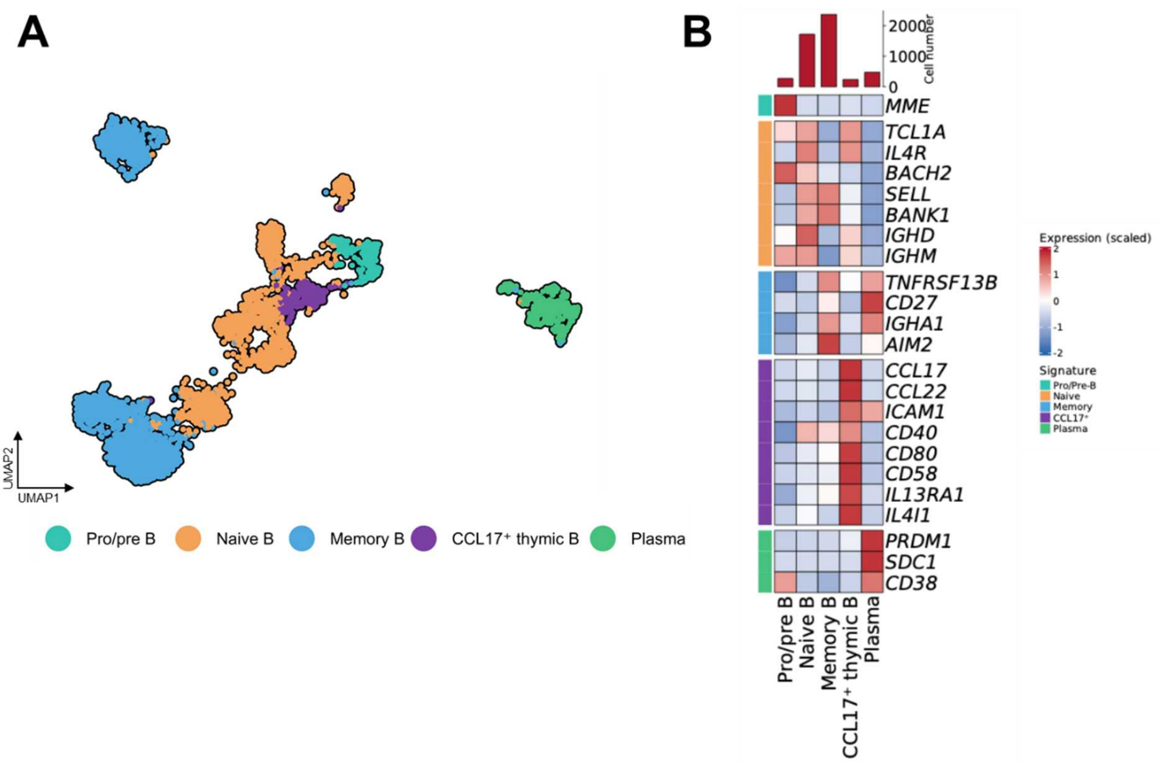


**Figure S3. Validation of thymic B-cell states in an independent external dataset.**

(A) UMAP visualization of public thymus scRNA-seq data. Each point represents a single cell, colored by B-cell identity.

(B) Heatmap showing the canonical marker expression in each cell cluster. Expression values are scaled across clusters (z-score), with red indicating higher and blue indicating lower relative expression.

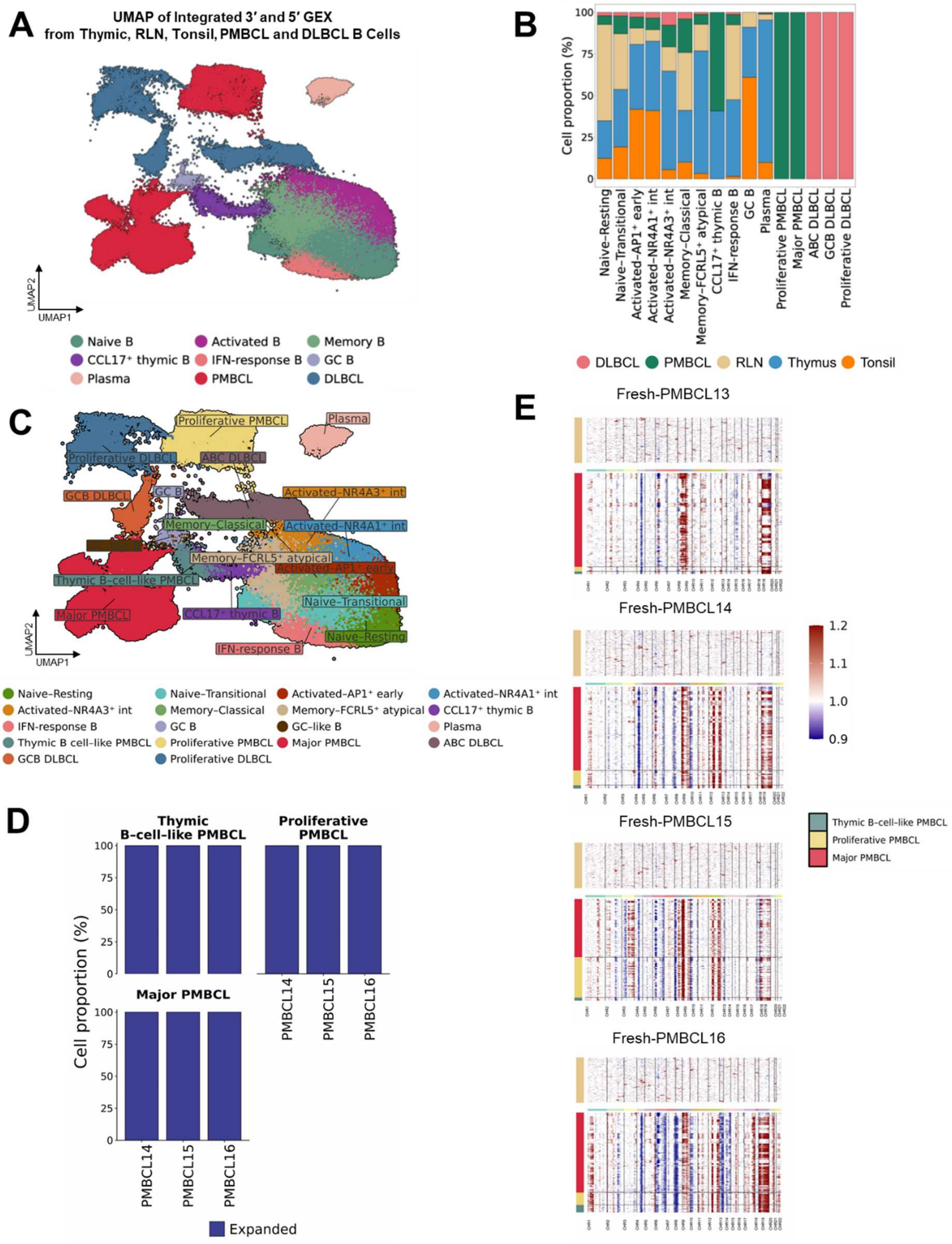


**Figure S4. Integrated analysis of single-cell B-cell atlas across tissues and lymphomas.**

(A) UMAP visualization of integrated gene expression data from single-nucleus multiome (3′ GEX + ATAC) and single-cell (5′ GEX + BCR) profiling of thymus, RLN, tonsil, PMBCL, and diffuse DLBCL samples. Each point represents a single cell and is colored by first-level B-cell identity.

(B) Composition of B-cell populations across tissues.

(C) UMAP visualization of refined annotation of B-cell populations revealing transcriptional heterogeneity within major lineages.

(D) Stacked bar plots showing, for each of three PMBCL cases (PMBCL14–16), the proportions of BCR-informative cells assigned to expanded versus unexpanded clonotypes within each of the three transcriptionally defined PMBCL states: major, proliferative, and thymic B-cell–like.

(E) Heatmaps showing large-scale copy-number variation (CNV) patterns inferred using inferCNV in Fresh-PMBCL13–16. Cells are grouped according to the three transcriptionally defined malignant states: thymic B-cell–like PMBCL, proliferative PMBCL, and major PMBCL. Each row represents an individual cell, and genomic regions are ordered by chromosome along the x-axis. Relative copy-number gains and losses are indicated by red and blue, respectively.

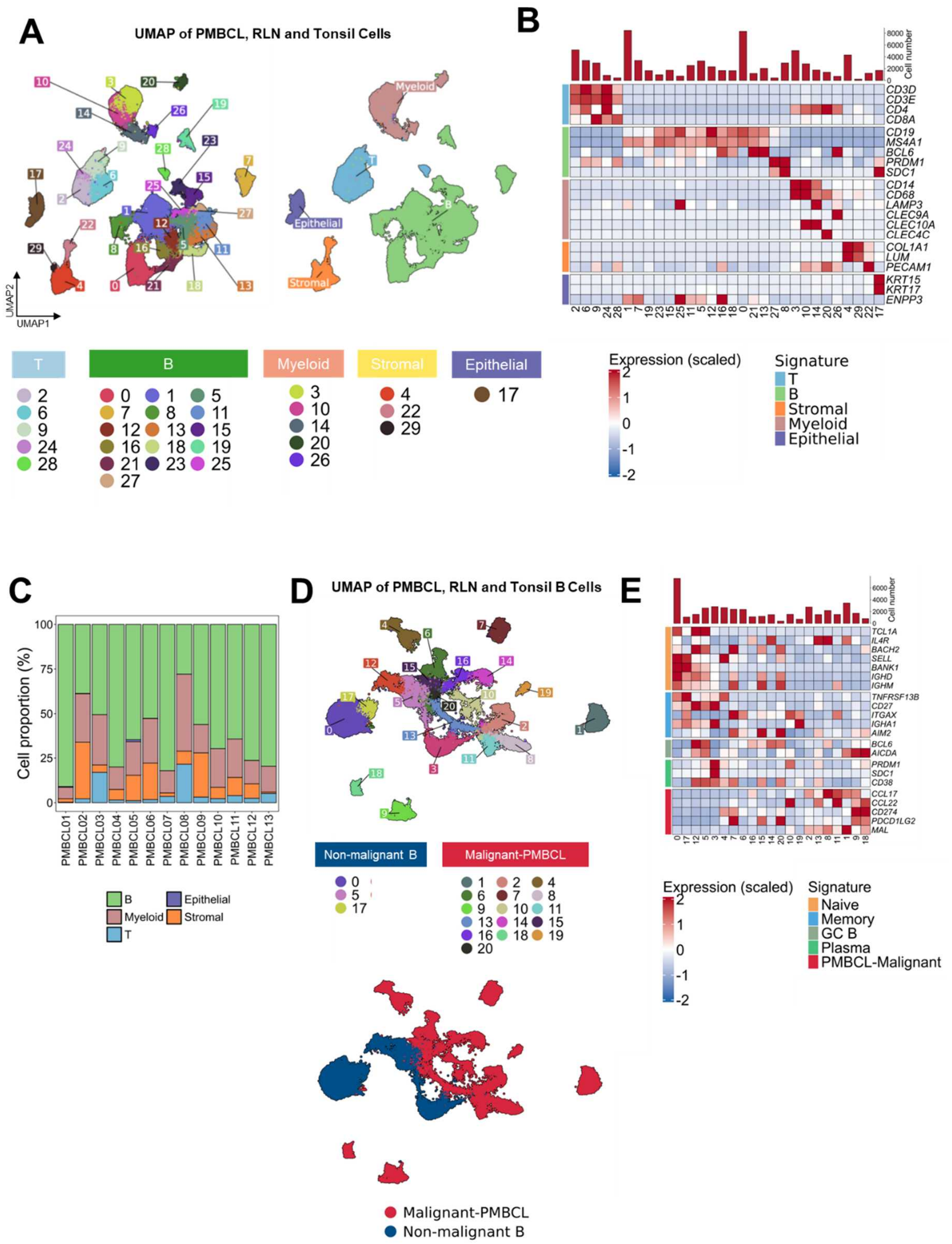


**Figure S5. Lineage annotation and malignant cell identification in FFPE snRNA-seq samples.**

(A) UMAP visualization of FFPE PMBCL, RLN and tonsil snRNA-seq data. Left: Each point represents a single cell, colored by cluster identity. Right: Each point represents a single cell, color coded by cell lineages.

(B) Heatmap showing the canonical marker expression in each cell cluster identified by single-nucleus RNA sequencing of PMBCL, RLN and tonsil samples. Expression values are scaled across clusters (z-score), with red indicating higher and blue indicating lower relative expression.

(C) Proportions of major cell populations in individual PMBCL samples.

(D) UMAP visualization of B-cell subsets in the FFPE PMBCL and RLN snRNA-seq data. Top: Each point represents a single cell, colored by cluster identity. Bottom: Each point represents a single cell, color coded by malignant B cell and normal B cell identity.

(E) Heatmap showing the canonical marker expression in B-cell clusters identified by snRNA-seq of PMBCL, RLN and tonsil samples. Rows represent selected lineage-defining genes, and columns represent transcriptionally defined cell clusters. Expression values are scaled across clusters (z-score), with red indicating higher and blue indicating lower relative expression.

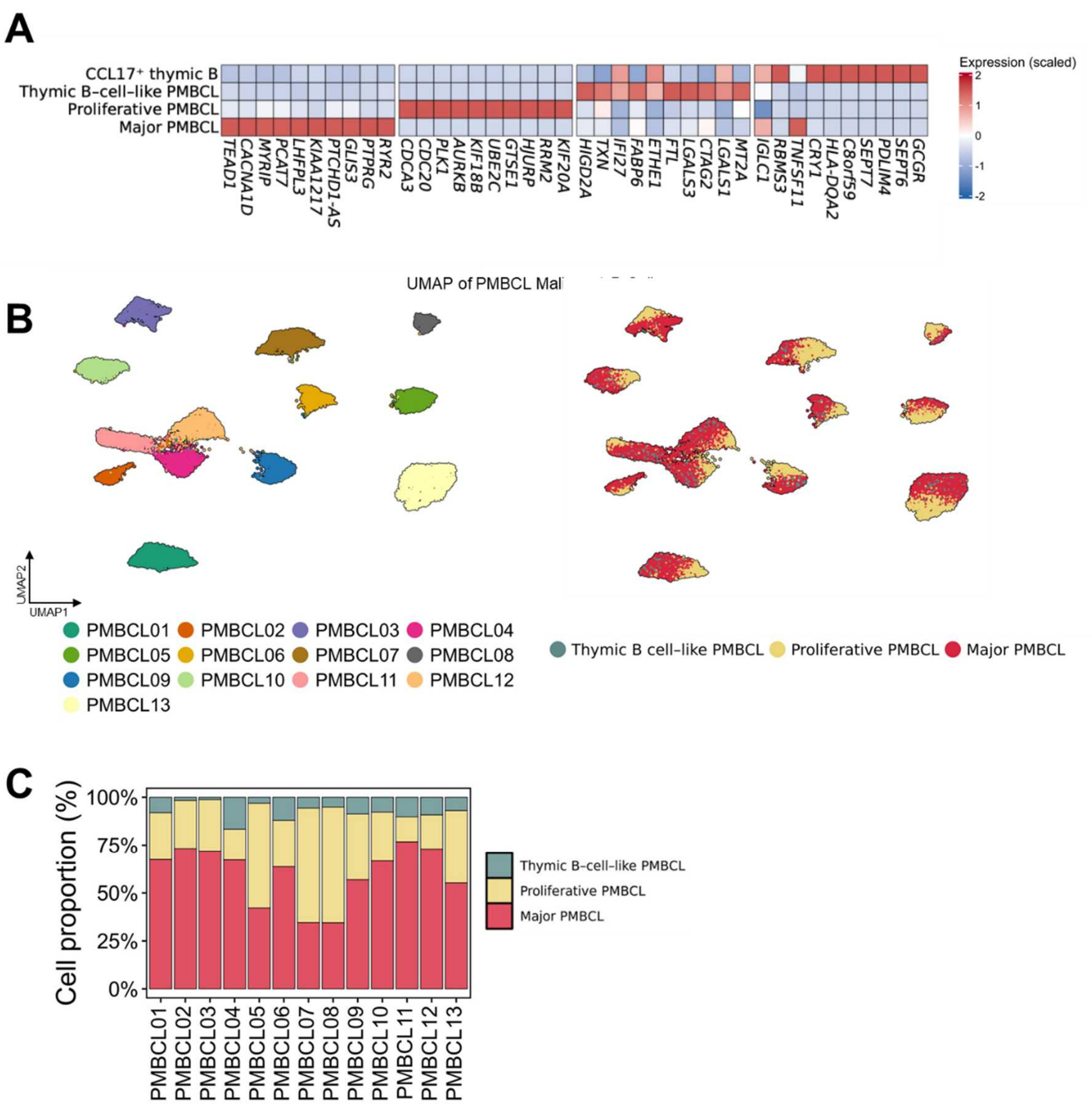


**Figure S6. Identification of malignant B-cell state within PMBCL malignant cells.**

(A) Heatmap showing the top 10 differentially expressed genes for each cell state across physiological CCL17⁺ thymic B cells and the three PMBCL states (major, proliferative, and thymic B-cell–like). Gene expression is scaled by row (z-score) to highlight relative enrichment patterns across states.

(B) UMAP visualization of malignant B-cell subsets in the FFPE PMBCL snRNA-seq data. Left: Each point represents a single cell, colored by patient identity. Right: Each point represents a single cell, colored by malignant B-cell states.

(C) Stacked barplot showing the proportion of malignant B-cell states in PMBCL samples.

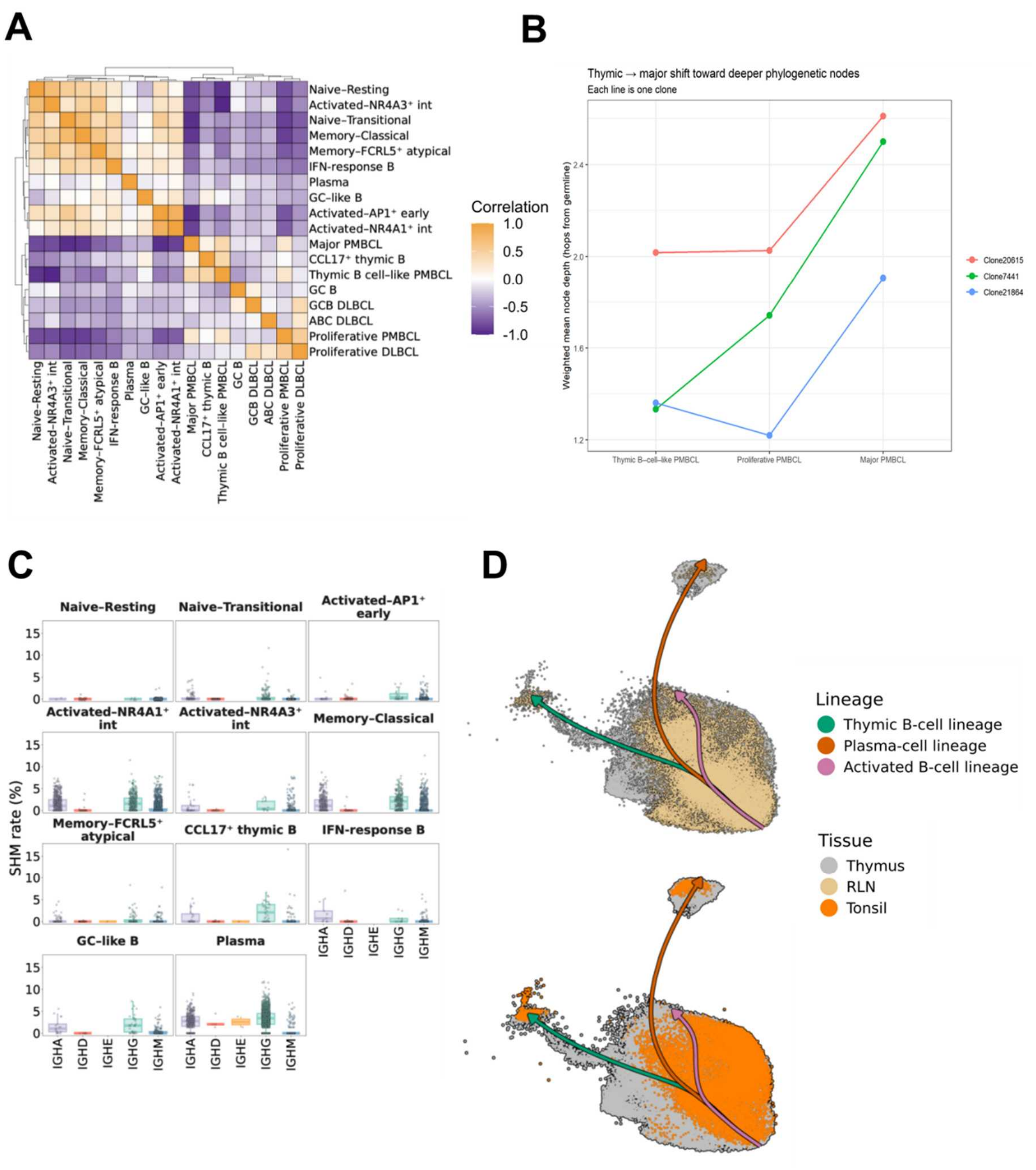


**Figure S7. Transcriptional relationships, SHM patterns, and inferred differentiation trajectories across B-cell states.**

(A) Pairwise correlation heatmap of gene expression profiles across transcriptionally defined normal and malignant B-cell populations. Colors indicate correlation coefficients, with orange representing higher similarity and purple representing lower similarity.

(B) Cell-count-weighted mean node depth per clone × cell type. Each line is one expanded clone. All three clones show major PMBCL cells at deeper nodes than thymic B-cell-like PMBCL cells; Clone21864 is the only clone in which the proliferative PMBCL cells are not intermediate. A linear mixed-effects model with clone as a random intercept showed a significant ordered increase in node depth across thymic B-cell–like, proliferative, and major PMBCL states (linear trend, $p = 0.0072$; quadratic component, $p = 0.091$). The thymic B-cell–like versus major PMBCL contrast remained significant after Tukey adjustment ($p = 0.0157$).

(C) Distribution of SHM frequencies stratified by immunoglobulin isotype across thymic B-cell populations.

(D) UMAP visualization of normal B-cell populations in RLN and tonsil samples overlaid with inferred differentiation trajectories reconstructed by pseudotime analysis.

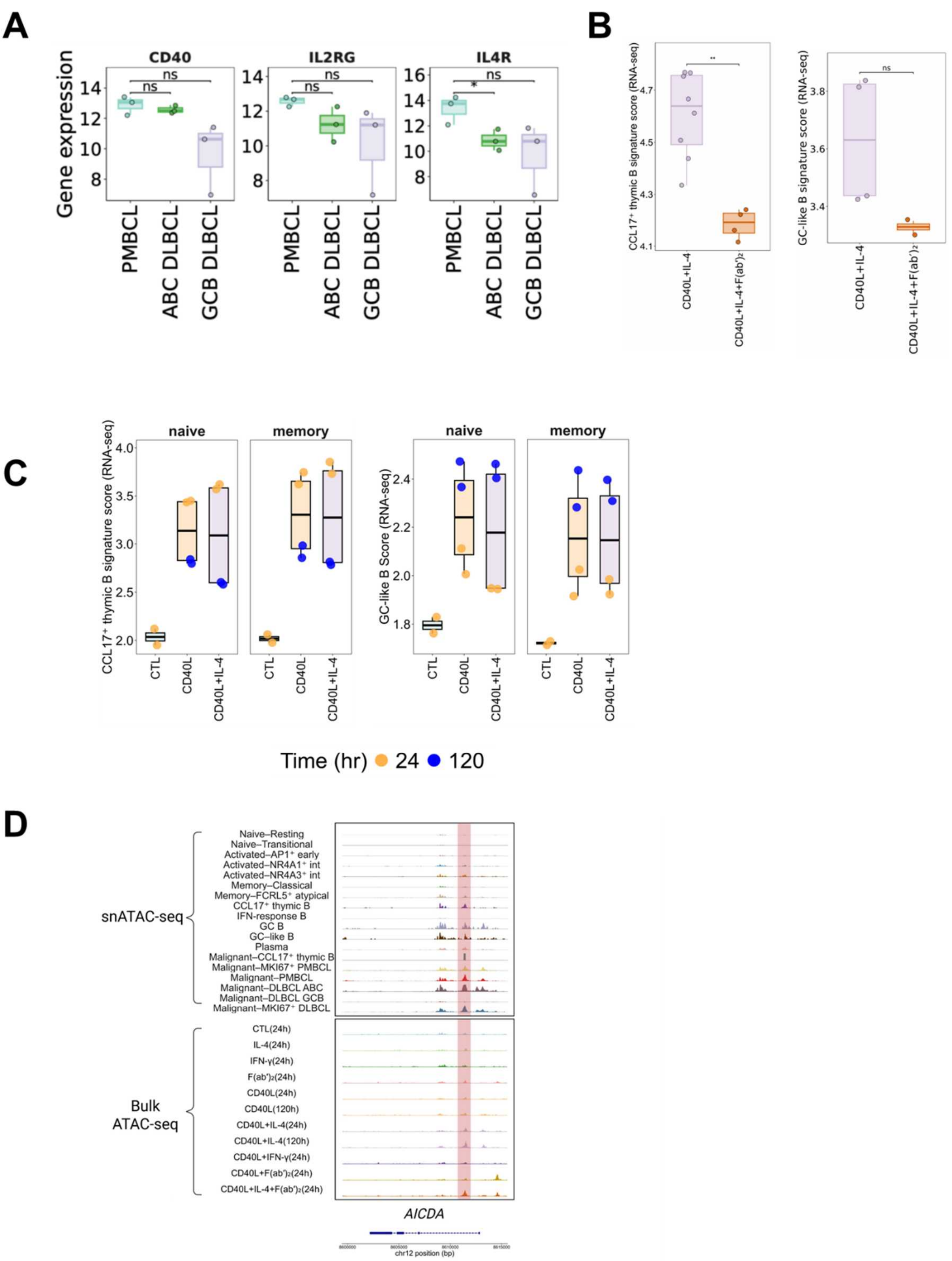


**Figure S8. Cytokine- and BCR-dependent regulation of CCL17$^{+}$ and GC-like B-cell programs.**

(A) Boxplots showing baseline relative expression of CD40 and the IL-4 receptor components

IL4R and IL2RG across the ABC-DLBCL, GCB-DLBCL, and PMBCL cell lines used in this study. Each point represents an individual cell line; boxes indicate the median and interquartile range. Statistical significance was assessed using two-sided Mann–Whitney U tests; $p < 0.05$; ns, not significant.

(B) CCL17$^{+}$ thymic B-cell (left) and GC-like B-cell (right) signature scores following *in vitro* stimulation. B cells were stimulated with CD40L and IL-4 in the presence or absence of BCR engagement (Fab). Signature scores derived from RNA-seq are shown for CD40L+IL-4 and CD40L+IL-4+Fab conditions. Boxplots summarize distributions across time points, with individual points representing biological replicates. Concurrent BCR engagement significantly reduced the CCL17$^{+}$ thymic B-cell signature score ($p < 0.01$) and produced a trend toward lower GC-like signature scores; ns, not significant. Statistical significance was assessed using two-sided Mann–Whitney U tests.

(C) Left: CCL17$^{+}$ thymic B-cell signature scores across stimulation conditions and time points following *in vitro* activation of naïve and memory B cells. Right: GC-like B-cell signature scores across the same stimulation conditions and time points. Boxplots summarize distributions across time points, with individual points representing biological replicates.

(D) Genome browser tracks showing chromatin accessibility at the *AICDA* locus.

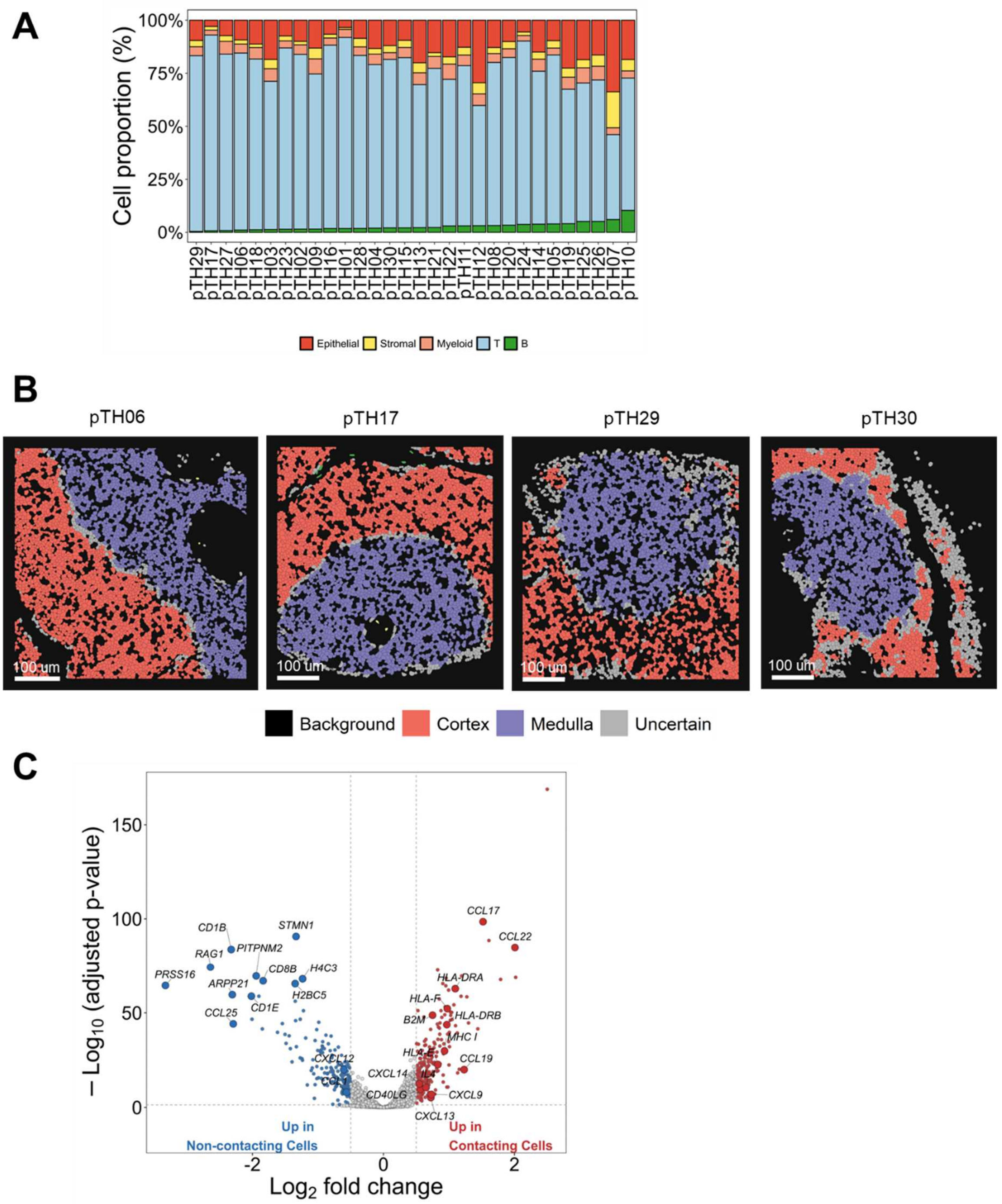


**Figure S9. Spatial compartmentalization and proximity-associated gene expression in the thymus.**

(A) Stacked barplot showing the cell-type composition across 30 human thymus samples profiled by CosMx spatial transcriptomics.

(B) Representative CosMx spatial molecular imaging section illustrating the spatial distribution of annotated thymic anatomical regions.

(C) Volcano plot showing the differential gene expression results comparing cells within 50 μm of CCL17$^{+}$ thymic B cells (Contacting Cells) versus more distant neighboring cells (Non-contacting Cells).

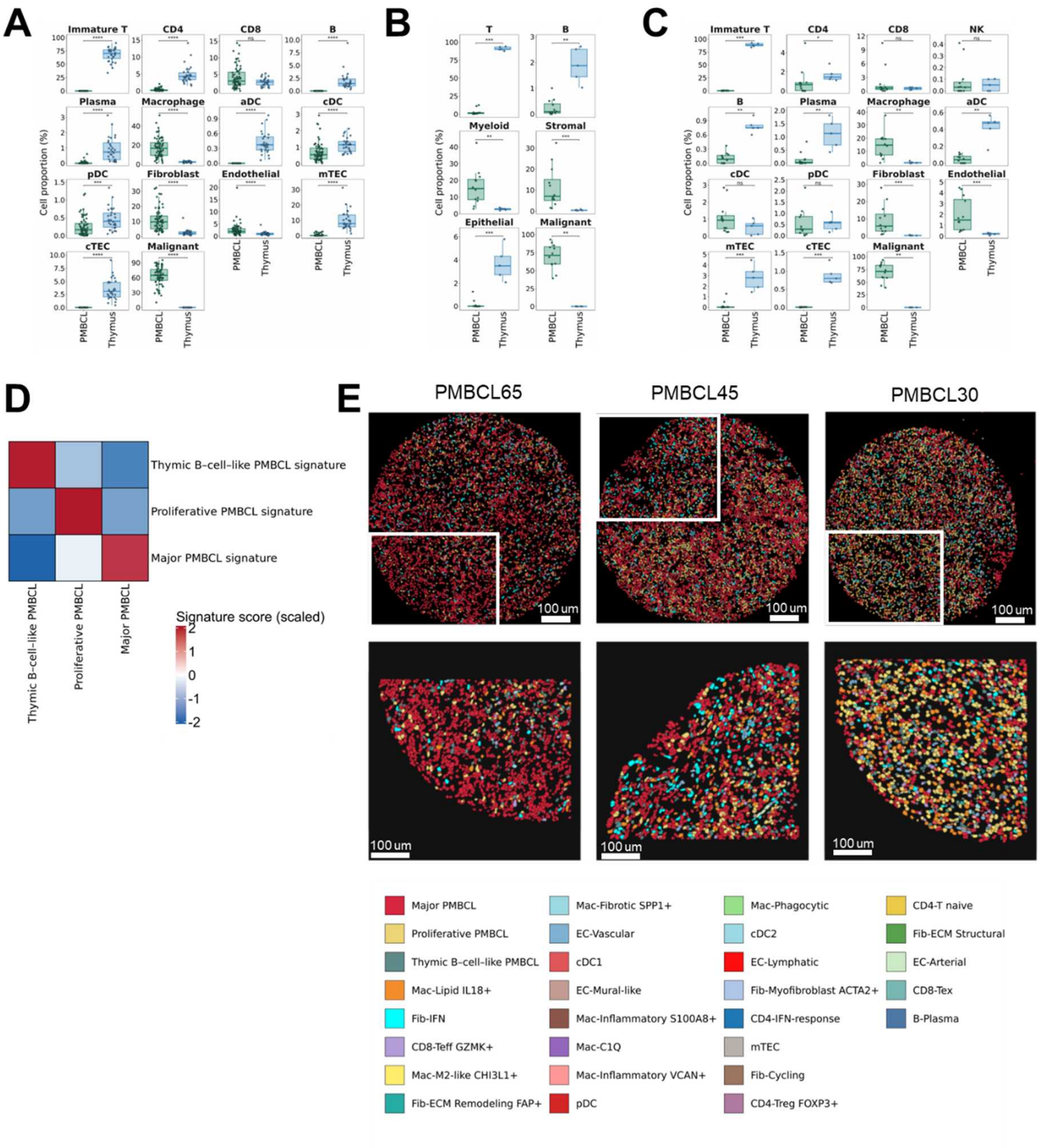


**Figure S10. Spatial compartmentalization of PMBCL states within a remodeled thymic microenvironment.**

(A) Comparison of detailed cellular composition between PMBCL and normal thymus in the CosMx spatial transcriptomic cohort. Boxplots show the relative proportions of annotated immune, stromal, epithelial, and malignant cell populations in each tissue. Statistical significance was assessed using two-sided Mann–Whitney U tests; $p < 0.05(^{*})$, $p < 0.01(^{**})$,

$p < 0.001$(***), $p < 0.0001$(****); ns, not significant.

(B) Comparison of major cellular compartments between PMBCL and normal thymus in the FFPE snRNA-seq cohort. Boxplots show the relative proportions of T cells, B cells, myeloid cells, stromal cells, epithelial cells, and malignant B cells. Statistical significance was assessed using two-sided Mann–Whitney U tests; $p < 0.05$(*), $p < 0.01$(**), $p < 0.001$(***), $p < 0.0001$(****); ns, not significant.

(C) Detailed comparison of cellular composition between PMBCL and normal thymus in the FFPE snRNA-seq cohort using refined cell-type annotations. Boxplots show the relative proportions of individual immune, stromal, epithelial, and malignant cell populations. Statistical significance was assessed using two-sided Mann–Whitney U tests; $p < 0.05$(*), $p < 0.01$(**), $p < 0.001$(***), $p < 0.0001$(****); ns, not significant.

(D) Heatmap showing scaled signature scores for the three PMBCL states identified by label transfer in the spatial dataset. Columns represent the thymic B-cell–like, proliferative, and major PMBCL states, and rows represent the corresponding transcriptional signatures derived from the reference dataset. Higher scores indicate greater enrichment of the respective state-specific transcriptional program.

(E) Representative CosMx spatial molecular imaging section illustrating the spatial distribution of annotated PMBCL cell types (scale bar, 300 μm). Individual segmented cells are colored by cell identity.
