## Supplementary Notes for "Developmental origins of thymic B-cell lymphomas"

* Authors contributed equally to this work.

This PDF file includes:

**Supplementary Notes:**

**Supplementary Note 1:** Quality assessment of FFPE snRNA-seq data

**Supplementary Note 2:** Evaluation of batch effects in FFPE snRNA-seq data

**Supplementary Note 3:** Quality assessment of fresh 5’ scRNA-seq data

**Supplementary Note 4:** Quality assessment of fresh 3’ snRNA-seq data

**Supplementary Note 5:** Evaluation of integration on 5’ scRNA-seq and 3’ snRNA-seq data

**Supplementary Note 6:** Quality assessment of fresh 3’ snATAC-seq data

**Supplementary Note 7:** Quality assessment of thymus CosMx data

**Supplementary Figures:**

**Supplementary Data Fig. 1:** *CCL17*$^{+}$ thymic B cell FACS gating strategy

**Supplementary Data Fig. 2:** B cell FACS gating strategy

**Supplementary Data Fig. 3:** Quality assessment of FFPE snRNA-seq data

**Supplementary Data Fig. 4:** Evaluation of batch effects in FFPE snRNA-seq data

**Supplementary Data Fig. 5:** Quality assessment of fresh 5’ scRNA-seq data

**Supplementary Data Fig. 6:** Quality assessment of fresh 3’ snRNA-seq data

**Supplementary Data Fig. 7:** Evaluation of integration on 5’ scRNA-seq and 3’ snRNA-seq data

**Supplementary Data Fig. 8:** Quality assessment of fresh 3’ snATAC-seq data

**Supplementary Data Fig. 9:** Quality assessment of CosMx data

**Supplementary Tables:**

**Supplementary Table 1.** Clinical characteristics of the FFPE, and fresh frozen scRNA-seq, and snRNA-seq cohorts

**Supplementary Table 2.** Clinical characteristics of the thymus CosMx cohort

**Supplementary Table 3.** Clinical characteristics of the PMBCL CosMx cohort

**Supplementary Table 4.** Differential gene expression analysis across PMBCL malignant cell states and physiological CCL17+ thymic B cells

**Supplementary Table 5.** Differential gene expression analysis between *CCL17*$^+$ thymic B cells versus thymic B–cell–like PMBCL cells

**Supplementary Table 6.** Differential transcription factor motif enrichment analysis across B-cell subsets identified by snATAC-seq

**Supplementary Table 7.** Gene ontology enrichment analysis across B-cell subsets

**Supplementary Table 8.** Ingenuity Pathway Analysis across B-cell subsets

**Supplementary Table 9.** CytoSig analysis across B-cell subsets

**Supplementary Table 10.** Differential gene expression analysis between *CCL17*$^+$ thymic B cells versus other B-cell subsets

**Supplementary Table 11.** Differential gene expression analysis between GC-like B cells versus other B-cell subsets

**Supplementary Table 12.** Differential gene activity analysis of *CCL17*$^+$ thymic B cells versus other B-cell subsets (snATAC-seq)

**Supplementary Table 13.** Differential gene activity analysis of GC-like B cells versus other B-cell subsets (snATAC-seq)

**Supplementary Table 14.** List of 200 custom add-on genes included in the CosMx SMI panel

#### Supplementary Notes

##### Supplementary Note 1: Quality assessment of FFPE snRNA-seq data

After stringent filtering to remove low-quality nuclei and multiplets, we retained 20,011 high-quality nuclei from thymus samples, 14,259 nuclei from reactive lymph node (RLN) samples, 18,073 nuclei from tonsil samples, and 43,100 nuclei from PMBCL (primary mediastinal B-cell lymphoma) samples.

Across all retained nuclei (n = 95,443), the distribution of detected genes per nucleus was unimodal, with a median of 752 genes (interquartile range [IQR], 561–1081; Supplementary Data Fig. 3a–b). Similarly, the distribution of unique molecular identifiers (UMIs) per nucleus displayed a narrow, unimodal profile, with a median of 868 UMIs (IQR, 636–1,317), reflecting robust transcript capture efficiency and limited technical variability across nuclei (Supplementary Data Fig. 3c–d). Consistent with high-quality library complexity, gene and UMI counts were positively correlated at the single-nucleus level, with no evidence of distinct subpopulations exhibiting disproportionately large or small library sizes (Supplementary Data Fig. 3e). Collectively, these quality control metrics indicate that the final dataset is composed of transcriptionally intact, well-captured single nuclei suitable for downstream analyses.

##### Supplementary Note 2: Evaluation of batch effects in FFPE snRNA-seq data

The FFPE snRNA-seq datasets were generated using two distinct sequencing chemistries (Next GEM Flex Gene Expression and GEM-X Flex Gene Expression), potentially introducing chemistry-associated technical variation. Prior to integration, cells showed partial segregation by sequencing chemistry in UMAP space, suggesting the presence of chemistry-associated batch effects (Supplementary Data Fig. 4a). To mitigate these effects while preserving biologically meaningful transcriptional differences, we performed batch correction using Harmony. Following integration, cells from the two sequencing chemistries showed improved mixing within shared transcriptional populations (Supplementary Data Fig. 4a).

To quantitatively evaluate batch correction performance and preservation of biological structure, we calculated the local inverse Simpson's index (LISI) and average silhouette width (ASW) before and after Harmony integration. Differences before and after integration were assessed using two-sided paired Wilcoxon signed-rank tests, with individual biological samples treated as the units of comparison. The integration LISI (iLISI) increased significantly following Harmony correction (P

< 0.001; Supplementary Data Fig. 4b), indicating improved batch mixing. Consistently, ASW with respect to sequencing chemistry decreased significantly (P < 0.01), further supporting the reduction of chemistry-associated technical variation. In contrast, cell-type LISI (cLISI) values remained close to 1 before and after integration, with no statistically significant difference (Supplementary Data Fig. 4b). Similarly, ASW with respect to cell type remained largely unchanged, suggesting that cell-type-specific transcriptional structure was preserved following integration. Together, these results demonstrate that Harmony effectively reduced batch-associated variation in the FFPE snRNA-seq dataset while maintaining biological cell-type separation, supporting its use for downstream integrative analyses.

**Supplementary Note 3: Quality assessment of fresh 5' scRNA-seq data**

Stringent quality control was applied to remove low-quality cells, multiplets, and residual non–B cell contamination. After filtering, 30,512 high-quality cells from thymus samples, 2,809 cells from RLN samples, 38,907 cells from PMBCL samples, and 9,931 cells from DLBCL (diffuse large B-cell lymphoma) samples were retained for downstream analyses.

Across all retained cells (n = 82,159), the distribution of detected genes per cell was unimodal, with a median of 1,718 genes (IQR, 1,285–2,456; Supplementary Data Fig. 5a–b). The distribution of unique molecular identifiers (UMIs) per cell similarly exhibited a unimodal profile, with a median of 3,939 UMIs (IQR, 2,806–6,385; Supplementary Data Fig. 5c–d).

The fraction of mitochondrial transcripts was low across the dataset, with a median of 2.14% (IQR, 1.25–3.69%; Supplementary Data Fig. 5e–f), indicating preserved RNA integrity and limited technical noise.

**Supplementary Note 4: Quality assessment of fresh 3' snRNA-seq**

For the fresh 3′ snRNA-seq datasets, nuclei were subjected to stringent filtering to exclude low-quality profiles, multiplets, and non–B cell contamination. This process yielded 15,909 high-quality nuclei from thymus samples, 27,393 nuclei from RLN samples, 21,958 nuclei from tonsil samples, 14,652 nuclei from PMBCL samples, and 9,928 nuclei from DLBCL samples.

Evaluation of transcriptomic complexity across retained nuclei (n = 89,840) revealed a unimodal distribution of detected genes, with a median of 1,194 genes (IQR, 967–1,686; Supplementary

Data Fig. 6a–b). Likewise, UMI counts per nucleus followed a unimodal distribution, with a median of 2,075 UMIs (IQR, 1,578–3,243; Supplementary Data Fig. 6c–d).

Consistent with expectations for nuclear RNA profiles, mitochondrial transcript fractions were modestly elevated relative to 5' scRNA-seq but remained low overall, with a median of 5.43% (IQR, 3.49%–8.19%; Supplementary Data Fig. 6e–f).

**Supplementary Note 5: Evaluation of integration on 5' scRNA-seq and 3' snRNA-seq data**

Gene expression matrices from all modalities were integrated using Harmony to enable joint analysis across tissues and lymphoma subtypes. Because the datasets were generated using distinct library preparation strategies (5′ scRNA-seq and 3′ snRNA-seq), integration was performed to reduce modality-associated technical heterogeneity while preserving biologically meaningful transcriptional differences between normal and malignant B cell populations.

Prior to integration, cells segregated predominantly by sequencing modality in UMAP space, with limited overlap between 5′ scRNA-seq and 3′ snRNA-seq datasets (Supplementary Data Fig. 7a), indicating substantial modality-driven technical variation. Following Harmony integration, cells from both modalities were well mixed within shared transcriptional states while preserving the global embedding structure (Supplementary Data Fig. 7a).

We applied the same batch effect evaluation framework used for the FFPE snRNA-seq dataset to assess integration performance. Following Harmony correction, iLISI increased significantly ($P < 0.0001$), indicating improved mixing between the 5′ scRNA-seq and 3′ snRNA-seq datasets (Supplementary Data Fig. 7b). Consistently, ASW with respect to sequencing technology decreased significantly ($P < 0.01$), indicating reduced modality-associated technical variation. In contrast to the FFPE snRNA-seq dataset, where cLISI remained largely unchanged, cLISI decreased significantly following integration ($P < 0.0001$), accompanied by an increase in cell-type ASW ($P < 0.0001$; Supplementary Data Fig. 7b). These findings suggest that Harmony effectively reduced modality-associated batch effects while improving cell-type separation and preserving biologically meaningful transcriptional differences. Together, these orthogonal metrics demonstrate that Harmony effectively mitigated technical heterogeneity arising from 5′ and 3′ library preparation strategies while preserving biologically meaningful cell-type structure.

**Supplementary Note 6: Quality assessment of fresh 3' snATAC-seq data**

snATAC-seq data quality was assessed using standard chromatin accessibility metrics implemented in Signac. High-quality nuclei exhibited strong enrichment of accessibility at transcription start sites, with a median TSS enrichment score of 5.37 (IQR, 4.64–6.04; Supplementary Data Fig. 8a). Fragment counts per nucleus were consistent with adequate library complexity (median, 7,765 fragments; IQR, 3,722–12,829; Supplementary Data Fig. 8b), and nucleosome signal scores indicated well-resolved chromatin fragmentation patterns (median, 0.64; IQR, 0.55–0.77; Supplementary Data Fig. 8c).

**Supplementary Note 7: Quality assessment of CosMx data**

Following cell segmentation and transcript assignment, the distribution of detected genes per cell exhibited a unimodal profile in thymus samples, with a median of 314 genes per cell (IQR, 205–451; Supplementary Data Fig. 9a). Similarly, transcript counts per cell followed a unimodal distribution, with a median of 403 transcripts per cell (IQR, 256–597; Supplementary Data Fig. 9b). In PMBCL samples, the median number of detected genes per cell was 180 (IQR, 98–311; Supplementary Data Fig. 9c), while the median transcript count per cell was 229 (IQR, 120–421; Supplementary Data Fig. 9d).

Collectively, these quality metrics demonstrate robust cell segmentation and reliable transcript detection suitable for downstream spatial and integrative analyses.

### Supplementary Figures

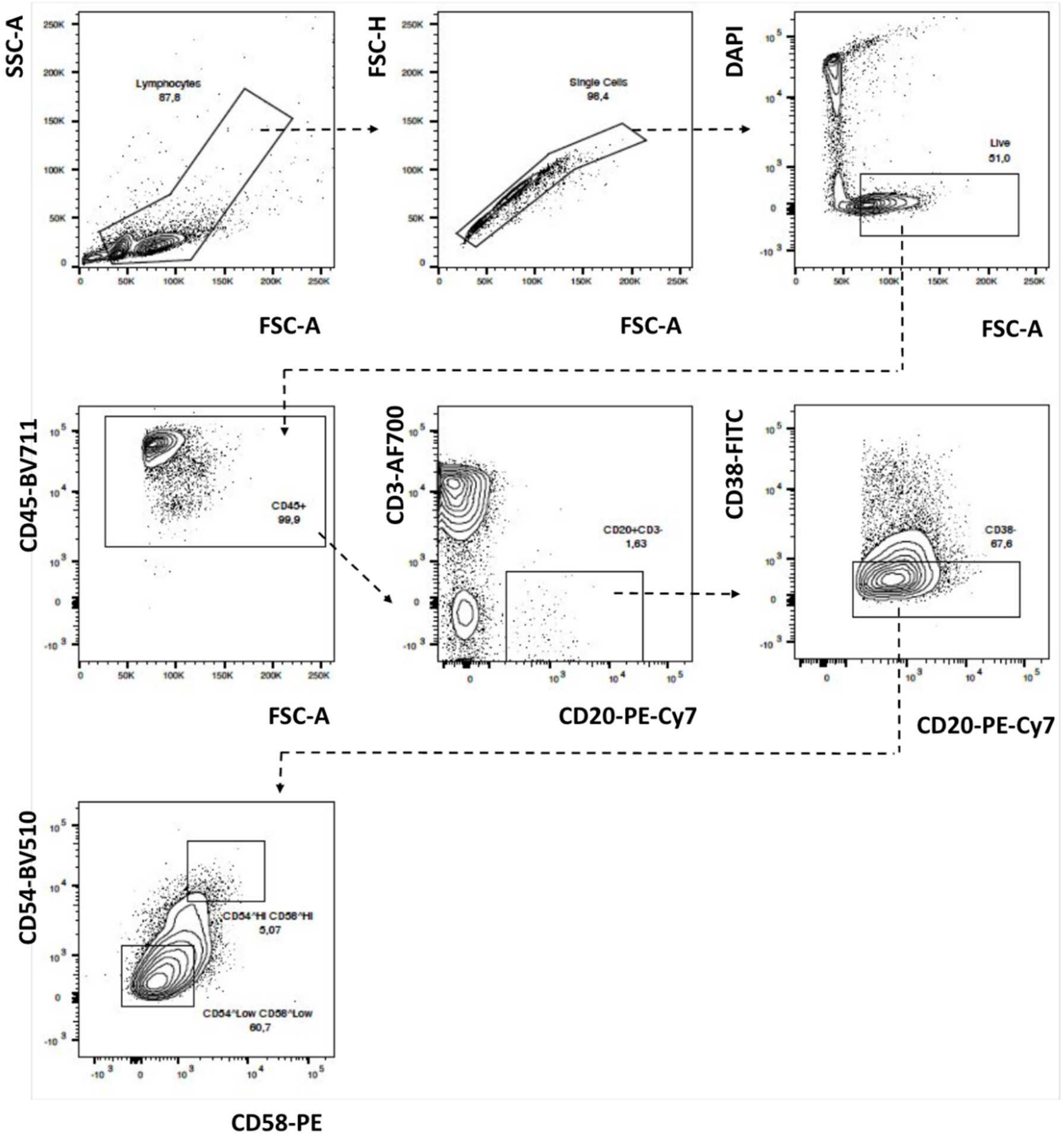


**Supplementary Data Fig. 1. *CCL17*$^{+}$ thymic B cell FACS strategy.** Representative sequential gating strategy for FACS for CD20$^{+}$CD38$^{-}$CD54hiCD58hi and CD20$^{+}$CD38$^{-}$CD54lowCD58low populations from normal human thymus samples.

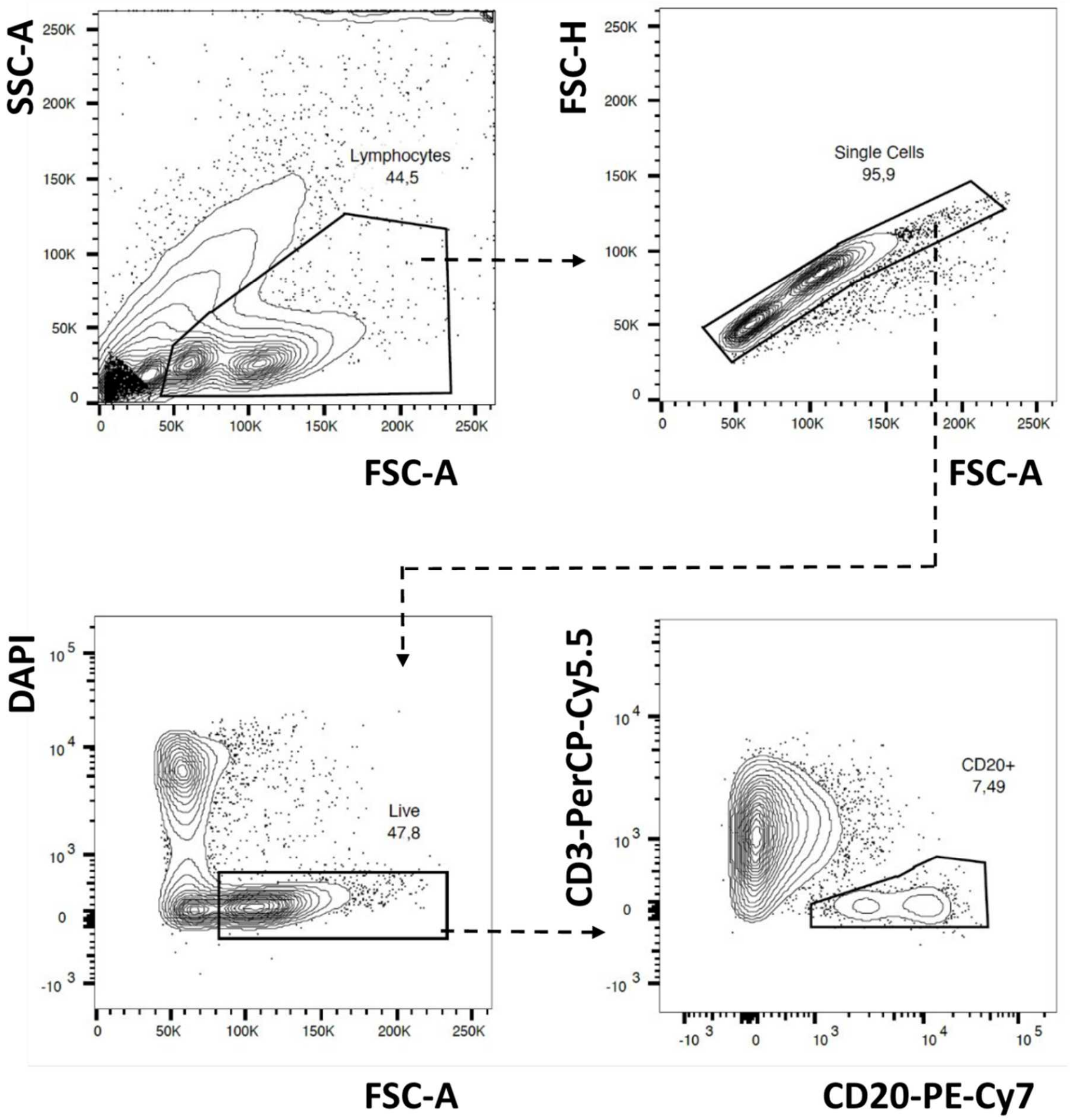


**Supplementary Data Fig. 2. B cell FACS gating strategy**. Representative sequential gating strategy for FACS for $CD20^{+}CD3^{-}$ populations for RLN, thymus, and PMBCL samples

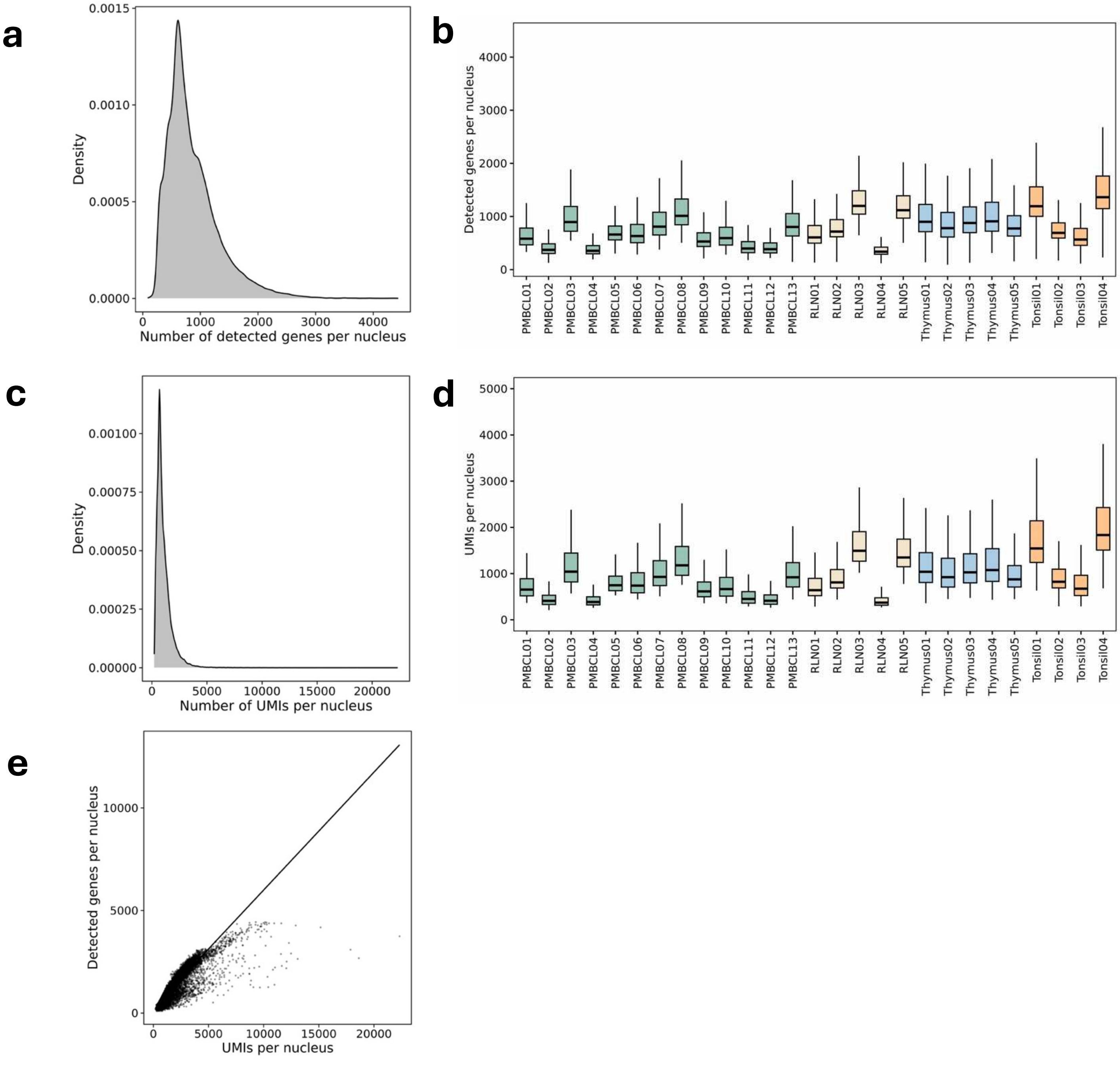


**Supplementary Data Fig. 3. Quality assessment of FFPE snRNA-seq data. a.** Density distribution of detected genes per nucleus. **b.** Per-nucleus gene detection across individual PMBCL, RLN, tonsil, and thymus samples. **c.** Density distribution of UMIs per nucleus. **d.** Per-nucleus UMIs across individual PMBCL, RLN, tonsil, and thymus samples. **e.** Relationship between UMIs and detected genes per nucleus.

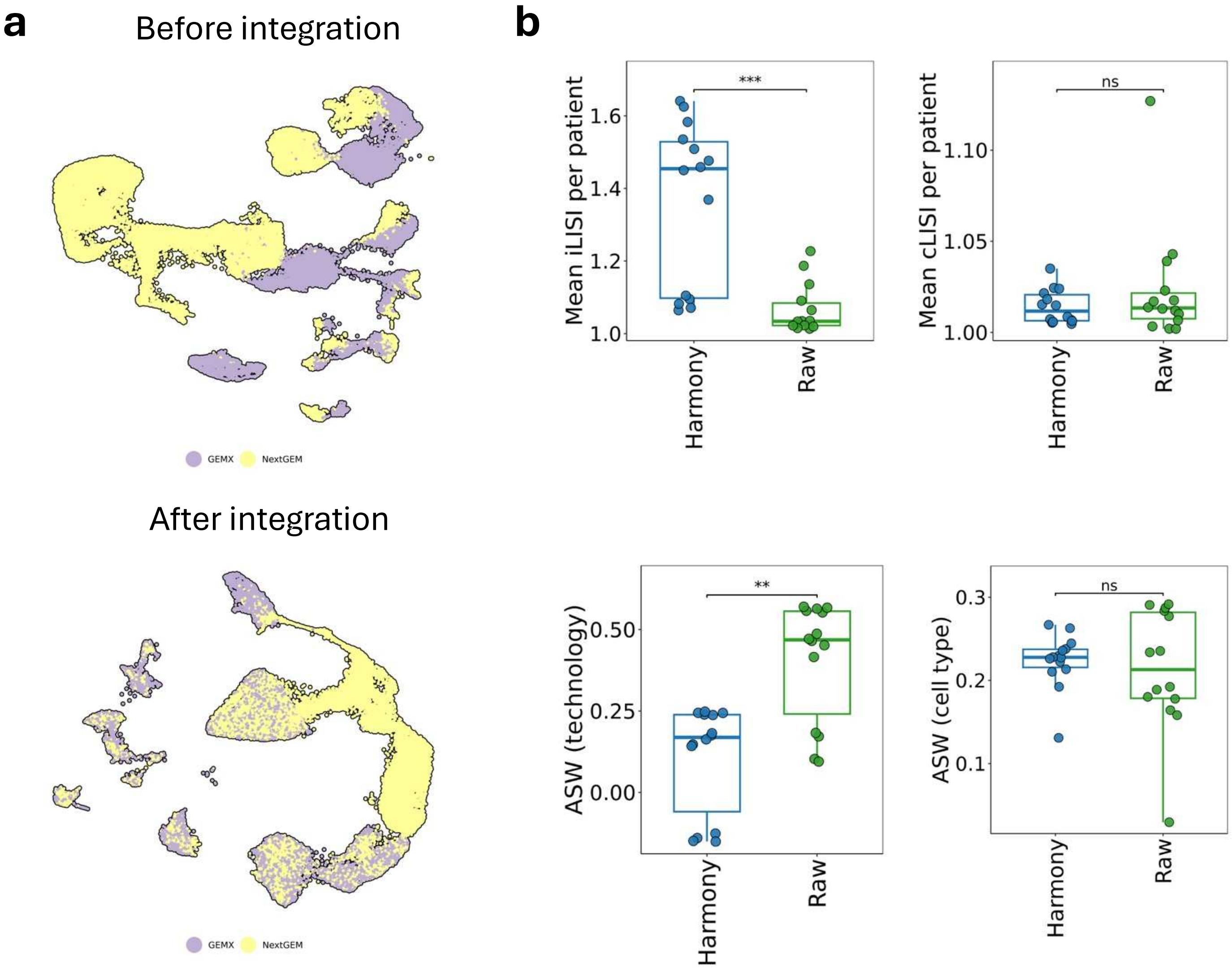


**Supplementary Data Fig. 4. Evaluation of batch effects on FFPE snRNA-seq data**. **a.** UMAP visualization of samples color coded by sequencing chemistries in the FFPE snRNA-seq dataset. **b.** Quantitative assessment of batch mixing and cell type purity using multiple metrics, including iLISI and cLISI, as well as ASW with respect to cell type and batch labels.

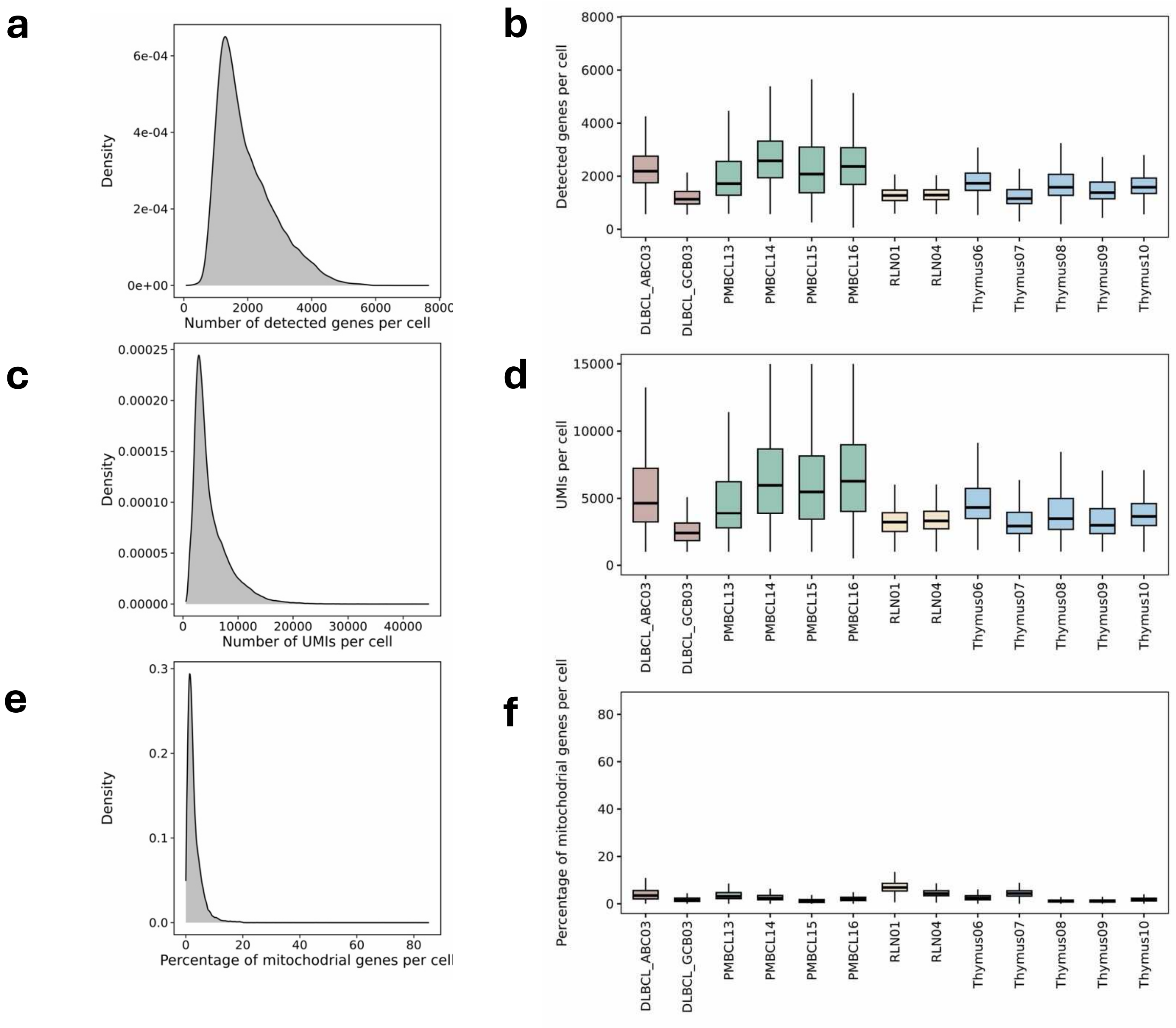


**Supplementary Data Fig. 5. Quality assessment of fresh 5’ scRNA-seq data**. **a.** Density distribution of detected genes per cell. **b.** Per-cell gene detection across individual thymus, RLN, PMBCL and DLBCL samples. **c.** Density distribution of UMIs per cell. **d.** Per-cell UMIs across individual thymus, RLN, PMBCL and DLBCL samples. **e.** Density distribution of percentage of mitochondrial genes per cell. **f.** Per-cell percentage of mitochondrial genes across individual thymus, RLN, PMBCL and DLBCL samples.

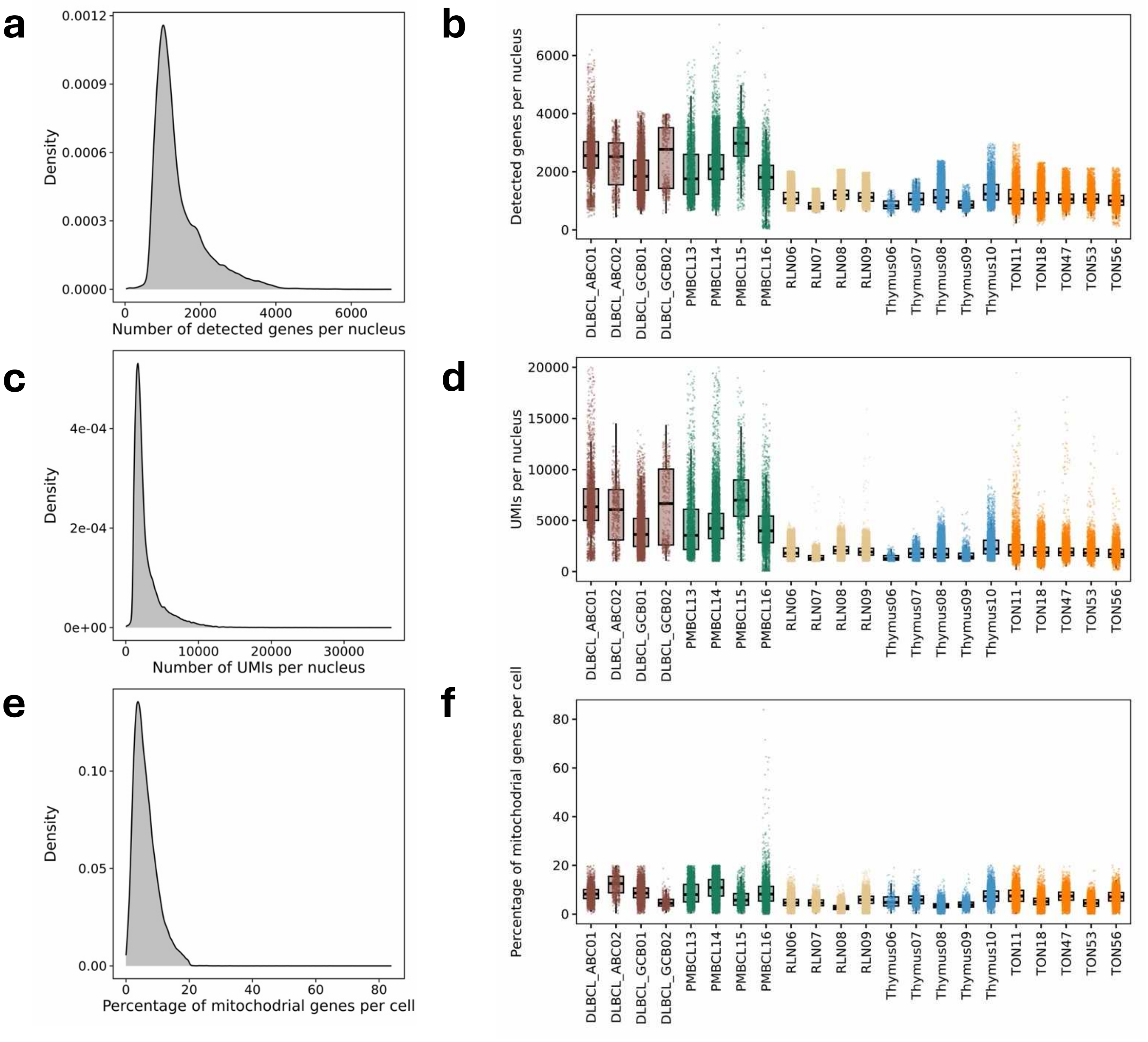


**Supplementary Data Fig. 6. Quality assessment of fresh 3' snRNA-seq data**. **a.** Density distribution of detected genes per nucleus. **b.** Per-nucleus gene detection across individual thymus, RLN, tonsil, PMBCL and DLBCL samples. **c.** Density distribution of UMIs per nucleus. **d.** Per-nucleus UMIs across individual thymus, RLN, tonsil, PMBCL and DLBCL samples. **e.** Density distribution of percentage of mitochondrial genes per nucleus. **f.** Per-nucleus percentage of mitochondrial genes across individual thymus, RLN, tonsil, PMBCL and DLBCL samples.

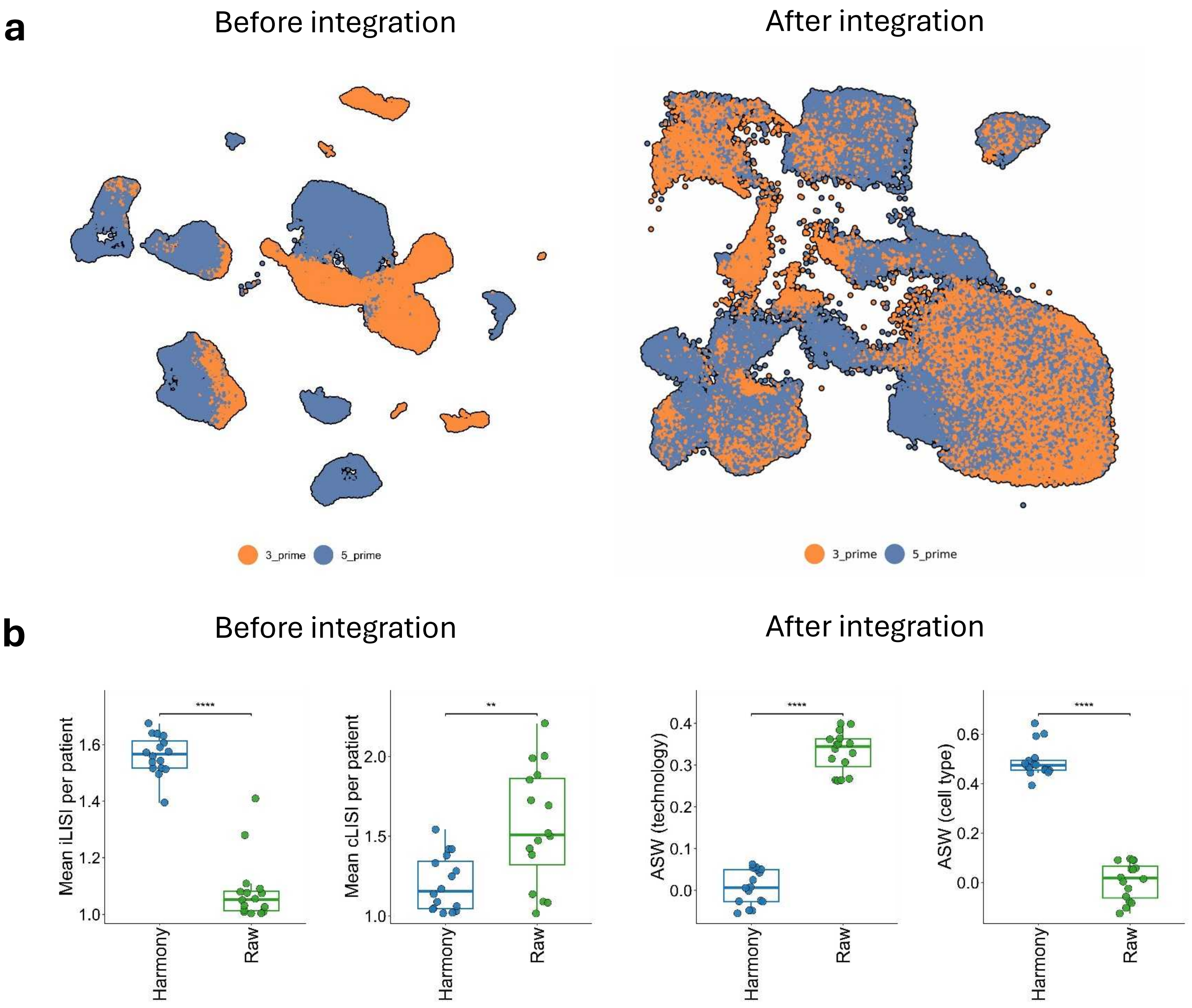


**Supplementary Data Fig. 7. Evaluation of batch effects in integrated gene expression data**. **a.** UMAP visualization of different data modalities before and after integration in the B cell atlas dataset. **b.** Quantitative assessment of integration performance and cell identity preservation before and after batch correction using cLISI, iLISI, ASW with respect to data-modalities and cell type labels.

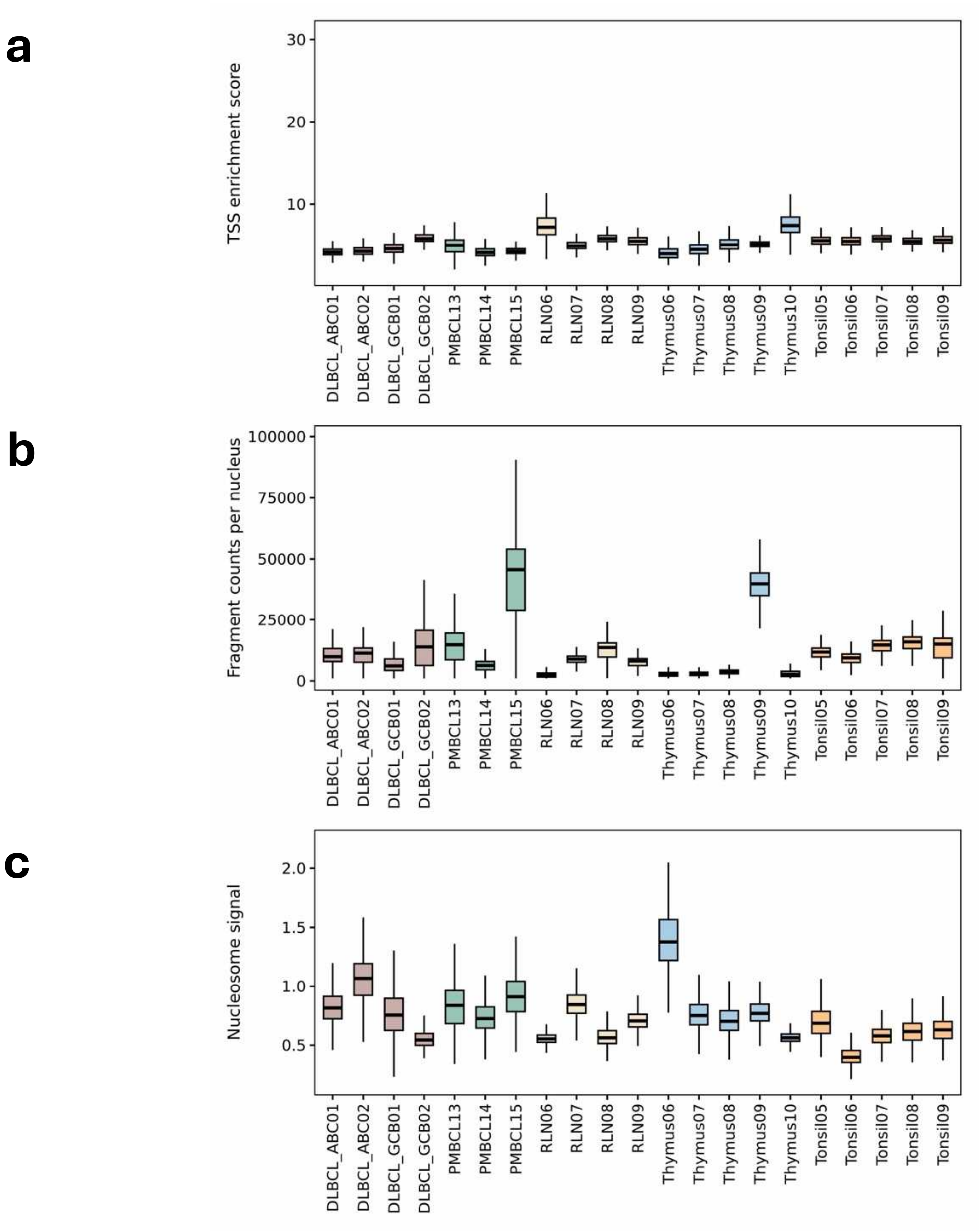


**Supplementary Data Fig. 8. Quality assessment of fresh 3' snATAC-seq data**. **a.** Per-nucleus TSS enrichment score across individual thymus, RLN, tonsil, PMBCL and DLBCL samples. **b.** Per-nucleus fragment counts across individual thymus, RLN, tonsil, PMBCL and DLBCL samples. **c.** Per-nucleus nucleosome signals across individual thymus, RLN, tonsil, PMBCL and DLBCL samples.

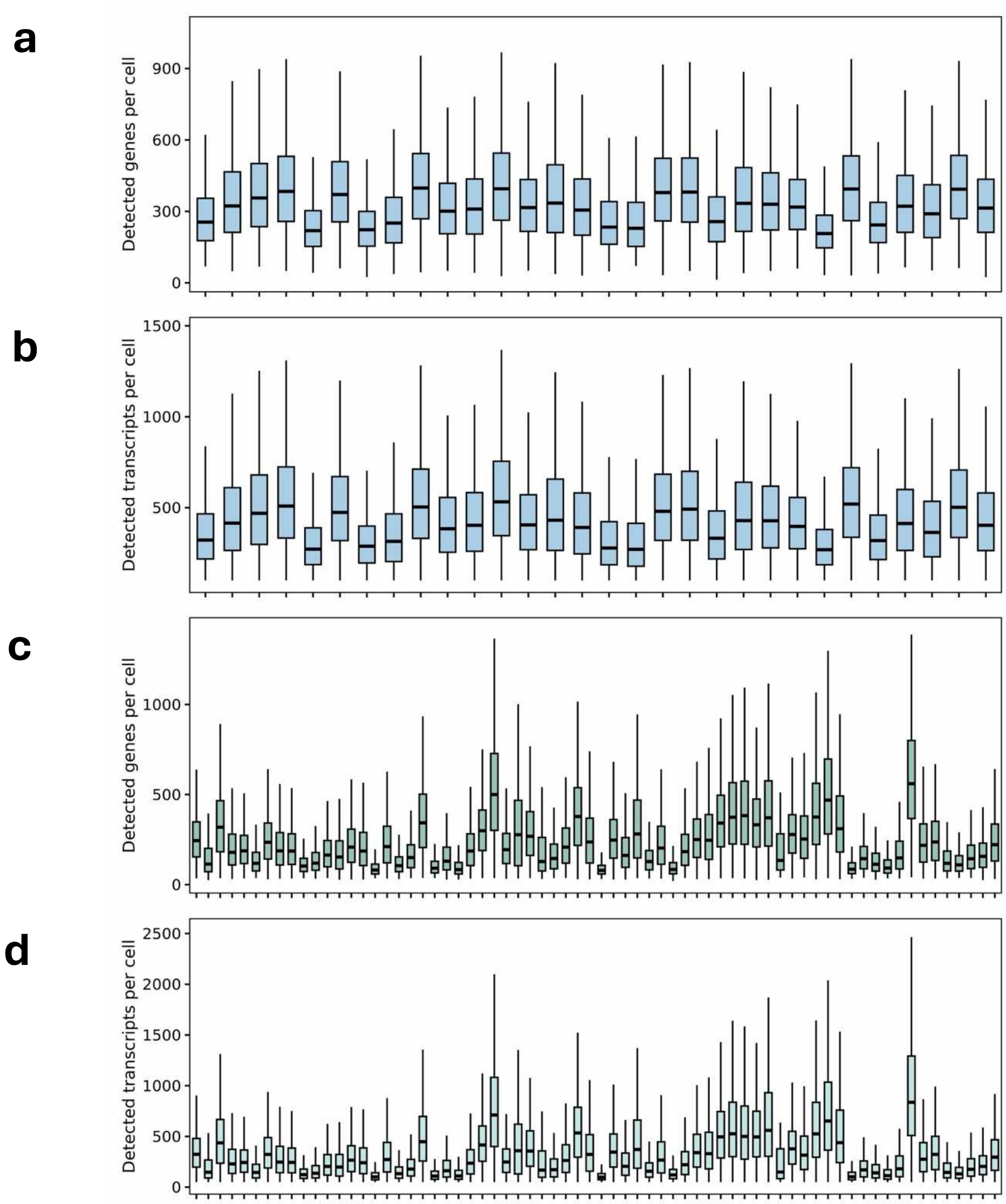


**Supplementary Data Fig. 9. Quality assessment of CosMx data**. **a.** Per-cell detected genes across individual thymus samples. **b.** Per-cell detected transcripts across individual thymus samples. **c.** Per-cell detected genes across individual PMBCL samples. **d.** Per-cell detected transcripts across individual PMBCL samples
